# Awake alpha bursting emerges as the dynamic working state in a lateral geniculate thalamocortical cell model

**DOI:** 10.64898/2026.06.08.730831

**Authors:** Kees McGahan, Michelle McCarthy, Nancy Kopell

**Affiliations:** Department of Mathematics and Statistics, Boston University, Boston, MA, USA

## Abstract

The awake thalamus is known to be able to filter primary sensory input with and without external modulation. Through the construction and analysis of a novel computational model of a lateral geniculate thalamocortical neuron, we demonstrate how the processing of sensory retinal input is influenced by the underlying thalamic dynamic state. Our model, using only currents verified against expression data from publicly available datasets, is the first to produce five experimentally established distinct dynamic firing regimes. We demonstrate that the thalamocortical cell transitions between these dynamic states in response to glutamatergic signals from the cortex or cholinergic arousal signals coming from the brainstem. We focus on signal processing in the model dynamic states associated with the awake thalamic alpha rhythm where we find that the ability of retinal inputs to generate thalamic spikes is a balance between the timing of retinal spikes, the excitability break imposed by the M-current, and the decay time of the L-type calcium current. Finally, we explore how these two currents help the thalamus process extra-retinal rhythmic inputs, showing the model produces entrainment to slower inhibitory and excitatory rhythms, as well as detailing the importance of nesting faster frequency rhythms within slow cycles for successful thalamic transmission. Our results suggest that the awake alpha rhythm is indirectly causal by acting as a marker for the interaction of these two currents. This biophysically-constrained lateral geniculate thalamocortical cell model generates predictions regarding rhythmic dynamics under different arousal states, thalamic control of retinogeniculate transmission, and the possible impacts neurological disorders, like schizophrenia, have on thalamic processing. Variations of this model could be used to explore the functions of higher order thalamic nuclei, thereby extending its use to investigating more complex cognitive processes.

**Author summary:** The thalamus generates multiple distinct brain rhythms, processes primary sensory inputs, and modulates its output using feedback signals. Previous computational models of the thalamus have typically focused on a subset of these three thalamic functions without drawing relationships among them. Here we present a novel computational thalamic cell model that unites these thalamic processes. We focus on the awake alpha rhythm, a well known thalamic oscillation, and show that it is a signature of a critical working state that enables the experimentally observed thalamic filtering of retinal signals. Additionally, we find this state is optimal for processing and passing non-sensory rhythmic signals. Our model generates testable predictions about which ionic currents control the transmission of external signals. It highlights the roles of two currents from our model that do not have specified functions in the awake thalamus in previous computational models. The work concludes with hypotheses about why neurological disorders that perturb the thalamus from this alpha rhythm working state lead to significant processing errors locally within the thalamus and globally within the brain.

## 1 Introduction

Historically the awake thalamus was perceived as a passive relay station [1–4]. Recent experimental advances have instead shown the thalamus acts as an active processing hub featuring complex dynamics. One thalamic nucleus that has been key in this perceptual shift is the dorsal lateral geniculate nucleus (LGN). While LGN thalamocortical (TC) cells receive modulatory input from numerous sources (thalamic reticular nucleus, visual cortex, intrageniculate interneurons) the dominant driver of LGN spikes comes from a few (1-6) synaptically connected retinal ganglion cells [3, 5, 6]. The efficacy of retinogeniculate transmission has been much investigated with both experiments and computation [4–13] in order to help understand what role the thalamus plays in processing sensory input.

Early experimental examinations of the LGN during different arousal states have revealed that arousal increases visually evoked LGN activity [4, 5, 14]. Later more detailed experimental studies, using s-potentials (extracellular events that represent retinal excitatory postsynaptic potentials) or dual unit recordings, were able to show that shorter retinal interspike intervals (ISIs) were more successful at driving LGN spikes [3, 5–7, 15]. In accompanying computational models, researchers showed that postsynaptic summation of the retinal inputs is likely the biggest determinant of which retinal inputs are transmitted, suggesting that presynaptic effects like facilitation and depression are not the main drivers of the ongoing filtering [5, 7, 8, 10–13, 16, 17]. However, the majority of data with this level of fine detail has been collected from paralyzed, anesthetized animals and thus there still exist a number of open questions pertaining to retinogeniculate transmission under non-anesthetized states [5–7, 9–13, 16, 17]. These include: what physiological features control transmission in arousal, why longer retinal ISIs are more likely to be transmitted in awake versus anesthetized animals, and what explains the heterogeneity between different LGN cell transfer efficacies in awake animals.

Complicating matters, in non-anesthetized animals, there are other thalamic functions/processes that occur concurrently with, and likely influence, retinal transfer. At the heart of these additional complexities are rhythmic oscillations. Across the full arousal spectrum, the thalamus is known to generate different rhythms whose timescales may be critical for understanding non-anesthetized animal retinal transmission [18–20]. Specifically, during the natural wakefulness state, where awake retinal transmission has primarily been studied [5, 13], the thalamus produces an alpha rhythm. However, how this rhythm is generated, how it interacts with sensory signals, how it interacts with rhythmic feedback from the cortex, and whether it is facilitating or impeding retinal transfer is still unclear [21–23]. Furthermore, it is well known the thalamus can be perturbed from producing this alpha rhythm in response to sensory signals, neuromodulatory signals [18, 24] or due to dysrhythmias resulting from neurological or psychological diseases and disorders [21, 25]. What happens to retinal transmission properties if either the underlying rhythmic oscillations or the cellular machinery is disrupted are also open questions.

Here we endeavor to connect two distinct areas of LGN study: retinogeniculate transmission and the possible types of rhythmic dynamics seen under different arousal states. More specifically, we investigate which cellular mechanisms dictate the dynamic arousal state of the thalamus and how these mechanisms control the transmission and filtering of incoming sensory (and non-sensory) signals. To do so, we have constructed a novel conductance-based (in the Hodgkin-Huxley formulation) LGN thalamocortical cell model that we show is capable of producing 5 distinct in-vitro firing regimes that correlate with experimentally observed arousal states. Using this conductance-based model framework we demonstrate how the thalamus transitions between states in response to neuromodulatory signals, and investigate how visual information might be gated, or transmitted, through the thalamus under different arousal states. The most interesting of these 5 states are the dynamic regions related to the natural wakefulness behavioral state (relaxed but alert) where the thalamus produces a firing type termed “high-threshold bursting” that coincides with the awake alpha rhythm [21, 26, 27]. Within these dynamic regimes we show the thalamus has the flexibility to filter retinal ganglion cell spike trains, showing a preference towards transmitting shorter retinal ISIs in a manner consistent with experimental findings [3, 5]. Our model suggests that the inclusion of an M current and an L-type calcium current in this thalamic model is instrumental for producing these observed in-vitro dynamics and the corresponding filtering of visual signals in these awake states. We use these results to suggest an explanation for the observed heterogeneity in transfer frequencies of different LGN cells [3, 5, 13] as well as to offer reasons for why high-threshold bursting is simultaneously a robust dynamic firing regime yet a largely hidden state in-vivo. Finally, we show that this level of model detail allows us to make additional predictions about the ability of the thalamus to interpret extra-retinal rhythmic inputs that may arrive as feedback directly from V1 visual cortex or indirectly via the thalamic reticular nucleus. Our model results lead to a new understanding of the awake thalamic alpha rhythm: the rhythm serves as a marker that the thalamus is in a working state. The associated filtering and entrainment capabilities that have been previously associated with changes in the alpha rhythm are dictated by interactions of the M current and L-type calcium current which happen to produce an alpha rhythm in the absence of input. In total, this study helps to unite retinal filtering findings from in-vivo LGN recordings with findings from in-vitro recordings regarding intrinsic thalamic rhythmic generation to make predictions about thalamic processing and how it is disrupted by diseases and disorders.

## 2 Results

### 2.1 Novel thalamocortical cell model is necessary to capture experimentally observed awake dynamics

Experimental studies (discussed in detail in the Methods) show that by increasing depolarization via an applied current, in the presence of a metabotropic glutamate receptor (mGluR1) agonist, a thalamocortical cell produces five distinct dynamic firing types [21, 28, 29]. These studies also reveal that at least one of these novel firing types, the high-threshold bursting alpha rhythm, is exhibited in the presence of cholinergic (mACHr) agonists [30]. Since previous models failed to capture the full observed dynamic range of these thalamocortical cells using validated ion channels (discussed in Methods), we constructed a new cellular model that contains both an M current (*I_M_*) and a high-threshold L-type calcium current (*I_CaL_*). Notably, all currents used in our model are expressed within the thalamus [31–33] and the ranges for their associated parameter values are restricted based on current clamp recordings in manners (and values) consistent with previous modeling studies (see Methods for details) [18, 31, 34, 35].

Our model produces five distinct dynamic firing types, including the elusive experimentally observed high-threshold alpha-frequency bursting state [30, 36]. The full dynamic range in the model can be found by decreasing the background potassium conductances, an action used in computational modeling to mimic either an applied cholinergic or glutamatergic agonist effect [18, 31, 35], and then continually increasing *I_app_* in order to depolarize the cell (Figure 1 and Supplemental Figure 1). Additional key pharmacological blocking experiments are replicated within the supplement [30, 36, 37] to show further relationships between the different ionic conductances and the dynamic firing regimes of the model (Supplemental Figure 2).

**Fig 1.**
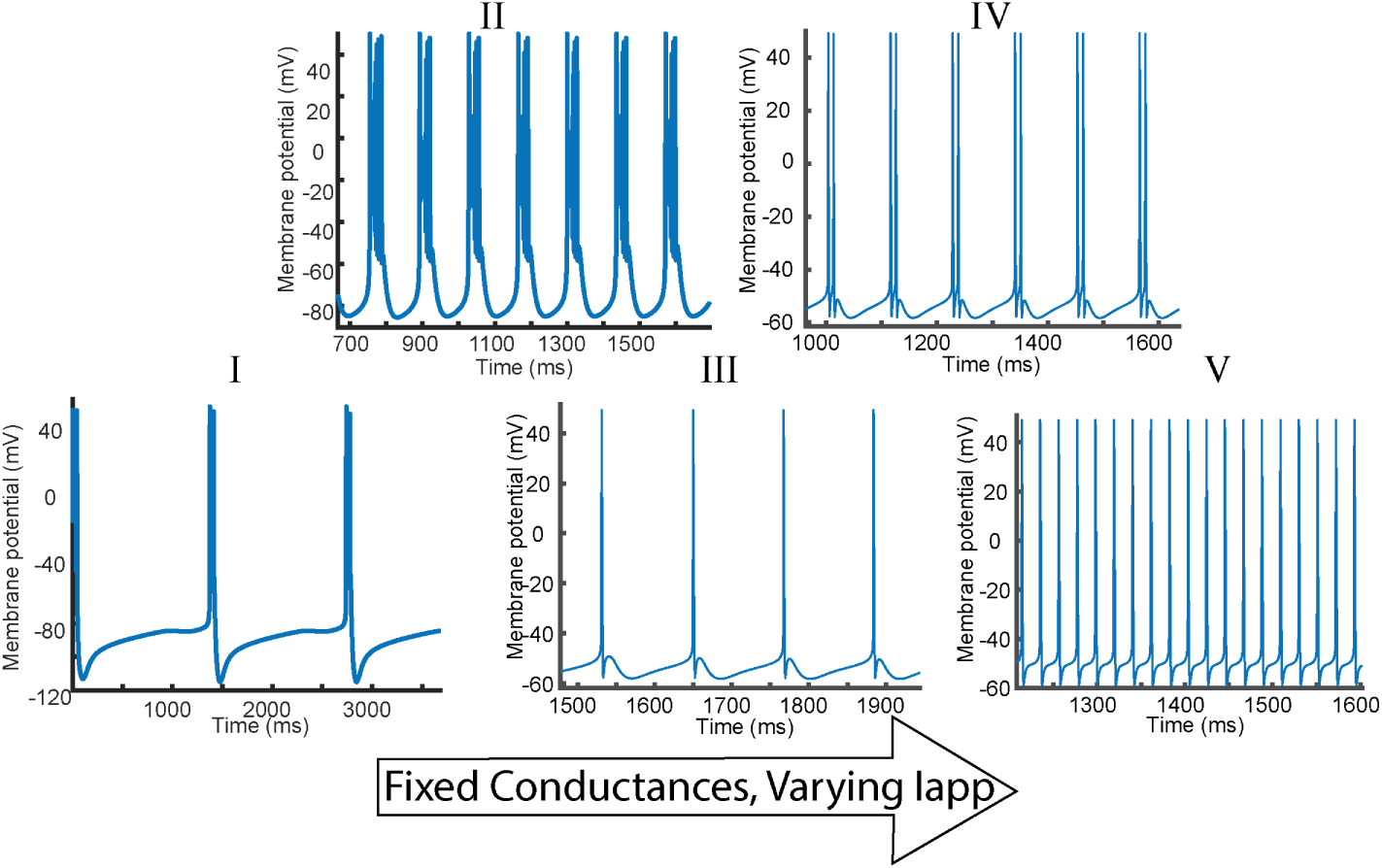
Thalamocortical cell model produces five distinct complex firing dynamics. With fixed ionic conductance values and increasing applied current (*Iapp*) from left to right, the model produces some notable firing patterns highlighted in the experimental work of Crunelli, Lorincz, and Hughes [27, 28]. In order the model dynamics are: **I** and **II** low-threshold bursting, **III** slow tonic firing, **IV** high-threshold bursting and **V** fast tonic spiking. A more detailed depiction (for 12 fixed values of *Iapp*) of the possible neuron dynamics is presented in (Supplemental Figure 1). Note, the distinction between low-threshold bursting dynamic states **I** and **II** are laid out in the next subsection. For comparison with experimental literature see: Figure 1 in [28] and Figures 1 and 2 in [37].

Critically, in addition to resembling experimentally observed firing types, these states can also be related to levels of arousal [18, 27, 28, 38]. Figures 1**I** and 1**II** show two low-threshold bursting states. The range of frequencies of low-threshold bursting includes *<* 1Hz frequencies that correspond to the slow wave sleep rhythms and between 1 − 4Hz frequencies that aligns with observed delta (1-4Hz) sleep rhythms (Supplemental Figure 1) [38]. The model also suggests that low-threshold bursting at greater than delta frequencies is possible (Supplemental Figure 1), which is discussed in greater detail in the next subsection in the context of the experimental literature. In addition to the sleep states, the model produces three distinct awake firing dynamics (Figure 1). Note that these states are categorized as “awake” firing based on minimum observed membrane potentials (*>* −60mV, discussed in Methods) and the lack of hyperpolarization events [27, 29, 37]. The first awake related firing type is a slow tonic spiking mode (missing from previous models [28]) that happens at theta (4-8Hz) to alpha frequencies (8-14Hz) (Figures 1 and Supplemental Figure 1). With slightly more applied current (*I_app_*), in Figures 1**IV** and Supplemental Figure 1 the model generates high-threshold bursts at theta and alpha frequencies, with experimentally matching intra-burst frequencies [27, 30, 36] (See section 2.9 for more details). Finally, additional increases in *I_app_* results in a fast tonic firing state at frequencies beta (20Hz) and above. The first two awake states (Figures 1**III** and **IV**) correspond to what we term “natural wakefulness” and correspond to an experimental awake behavioral state in which prominent alpha rhythms can rapidly emerge in occipital EEG by closing the eyes [21, 29]. The fast tonic firing seen in Figure 1**V**, is the most excited state, with a firing frequency that is associated with attentional processing [18, 39, 40].

While the five states in Figure 1 were found under fixed ionic conductance levels by varying the external excitation *I_app_*, experimental work has shown some of these state transitions occur in the presence of neuromodulation [41–44]. Specifically, glutamate (Glu) and acetylcholine (ACh) are implicated in the transition from sleep to waking states via their respective activation of metabotropic glutamate receptors (mGluRs) and muscarinic acetylcholine receptors (mAChRs) [42, 44–48]. The next subsection investigates how these neuromodulatory signals act in the model, both independently and in tandem with changes in external excitation.

### 2.2 Neuromodulatory arousal signals switch the thalamic cell from low threshold bursting to awake dynamic states

It is well documented that glutamatergic signals from the cortex and cholinergic signals from the brainstem play a strong role in transitioning the thalamus from sleep states to wakeful states [42, 44–48]. These arousal signals are typically instantiated in conductance-based thalamic models by decreasing the background potassium channel conductances of *I_kl_* or *I_M_* [18, 35, 43, 49]. This action replicates one (of many) downstream effects of glutamate and acetylcholine via distinct signaling pathways involving mGluR and mAChR receptor (both expressed throughout the thalamus) activation [32, 33, 44, 50–53]. Depending on receptor and ion channel expression levels these two neuromodulatory signals may alter one, or both, of *I_kl_* or *I_M_* to varying degrees; for simplicity, moving forward we refer not to the specific neuromodulator but to the modified ionic current. In this subsection, the M-current conductance is fixed and *g_kl_* is varied. In a later subsection the M-current conductance is varied while *g_kl_* is fixed, showing that *I_M_* has a more nuanced impact on the observable dynamics. The differences between glutamatergic and cholinergic arousal signaling (and their impacts on modeling) are examined in greater detail in the discussion section 3.2.

Figure 2 shows how model arousal state varies with changes in potassium leak conductance and applied external current. The center diagram in Figure 2 plots the dynamic behavior of the TC cell as a function of *I_app_* (x-axis) and *g_kl_* (y-axis). Each solid curve (black, purple) denotes a transition in parameter space from one dynamic firing state to another. The purple curve is highlighted as a key transition from sleep to wakefulness (STW). Each sectioned region (0-V) is categorized by their exemplified firing dynamics (i.e. quiescent, low-threshold bursting, high-threshold bursting. tonic spiking, or some combination of these) provided in the accompanying voltage plots. Dynamic state labels are consistent with Figure 1.

**Fig 2.**
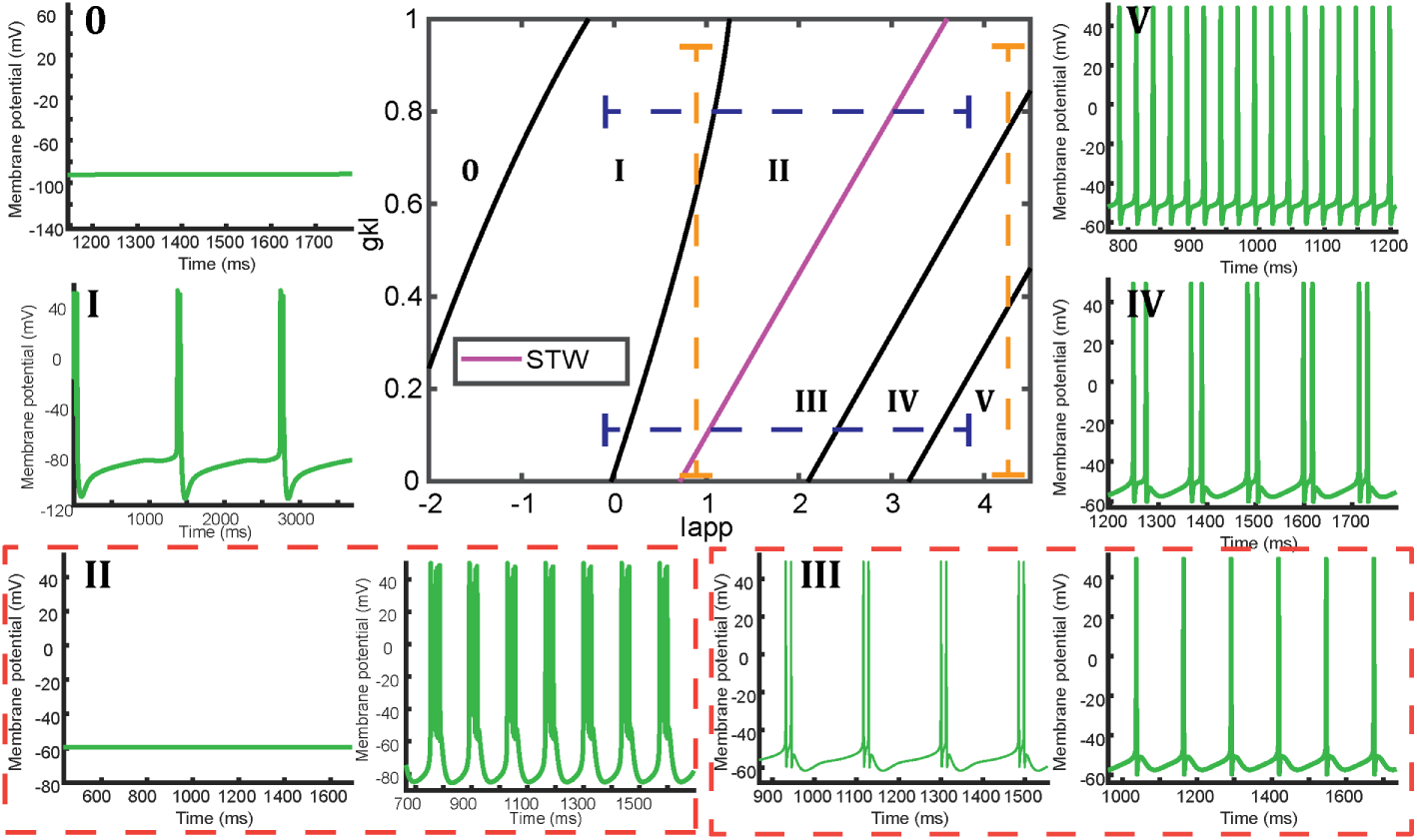
Description of the relationship between the thalamocortical model state of arousal, potassium leak conductance and external excitation. (*Iapp*). Solid lines in the central plot are used to divide parameter space into a total of six dynamically distinct regions (0-V). The magenta curve denotes the sleep to wake (STW) transition. Dashed orange and blue lines show transitions through parameter space either vertically or horizontally by fixing the corresponding parameter. Sample voltage traces for the TC cell from within each region are provided. The order of firing dynamics for these regions corresponds to the dynamics of Figure 1. Two differences from Figure 1: the quiescent hyperpolarized state (0) is a newly described state, and regions II and III have two different corresponding membrane potential plots (depicted in red dashed boxes) depending on the cell’s initial conditions. Note the complete bifurcation diagram with labeled mathematical bifurcation curves can be found in Supplemental Figure 3.

Figure 2 also reveals additional dynamic features not captured by Figure 1, namely that regions II and III are bistable; different initial conditions can lead to 2 different firing type outputs. Importantly, one of the bistable states in region II is a depolarized quiescent state that is commonly observed during slice experiments when transitioning from low-threshold bursting to tonic spiking with increasing *I_app_* [28]. This bistability (and existence of the depolarized quiescent state) of the model is critical for explaining why frequencies higher than delta (1-4Hz) can be produced by our model even though they are not typically seen in-vitro. The model shows that experimentally observed low-threshold bursting frequencies (between 0-6Hz) occur in region I as the only stable state, while the higher (not typically experimentally observed) frequencies occur only in the bistable region II (Supplemental Figure 4). However, in the absence of external inputs that can activate the T-type calcium channel, the model suggests that low-threshold bursts at frequencies greater than delta will be rare events: instead the silent depolarized state will emerge as is commonly seen with increasing *I_app_*in experiments [28].

Experimentally, the full dynamic range was observed in slice experiments only after application of a glutamatergic agonist [29, 30]. Without glutamatergic agonist application, experimentalists observed a smaller number of distinct dynamic states [28]. The model could help explain this experimental observation with the central plot of Figure 2 by considering two levels of modulated *g_kl_* (proposed conductance altered by the applied agonist) and varying *I_app_*across an identical fixed range (blue horizontal dashed lines). At high *g_kl_*(low levels of agonist), the model reaches only three states whereas all 5 states are reached with low *g_kl_*(high levels of agonist) for an identical range of *I_app_* (Figure 2). Furthermore, the bistability of region II (combined with the increased width of region II for high *g_kl_*) could help explain why, prior to agonist application, experimentalists could observe a quiescent state that was nonexistent (model quickly transitions from state I to state III low *g_kl_*) afterwards [28]. These results indicate that neuromodulatory changes of *g_kl_* could increase the total number of possible observed dynamic states, while also impacting which state transitions are possible.

Figure 2 also shows the impact of modulating *g_kl_* if external excitation (*I_app_*) is kept constant (orange vertical dashed lines). Starting in any of the awake states (III,IV,V), for any fixed value of *I_app_*, increasing *g_kl_*from 0 brings the neuron to a lower state of arousal. In spite of this, one subtle observation is that, with fixed excitation, the neuron cannot transition through the full range of states purely with *g_kl_* manipulation (Figure 2 orange lines). While modulation of *g_kl_* is sufficient to transition the thalamus between light sleep and resting awake states, reaching the slowest low-threshold bursting frequencies (*<* 1Hz) associated with slow wave sleep (found at the left edge of region I in the model), from any of the awake states (III,IV,V), requires additional changes in external thalamocortical cell input (Supplemental Figure 4) [42].

This model is uniquely positioned to explore how retinogeniculate transmission changes with arousal. It is crucial that the model captures the full dynamic range in order to identify differences in transmission during sleep states (1**I** and 1**II**), the natural wakefulness states (1**III** and 1**IV**), and the highly excited state (1**V**). How transmission probability changes as a function of LGN state is the focus of the next subsections.

### 2.3 LGN transmission of retinal signals depends on model arousal level

It is well known the retina can send spiking signals to the LGN during each of the different states of arousal [54, 55]. To investigate how these signals are processed as arousal level is changed, a spiking retinal ganglion cell is connected via an excitatory AMPA synapse to the model LGN thalamocortical cell (see Methods). The interspike intervals for the simulated retinal spike trains are then generated from a gamma distribution (see Methods) [10, 56]. Figure 3 shows how retinal input is processed by the LGN during the different LGN arousal states.

**Fig 3.**
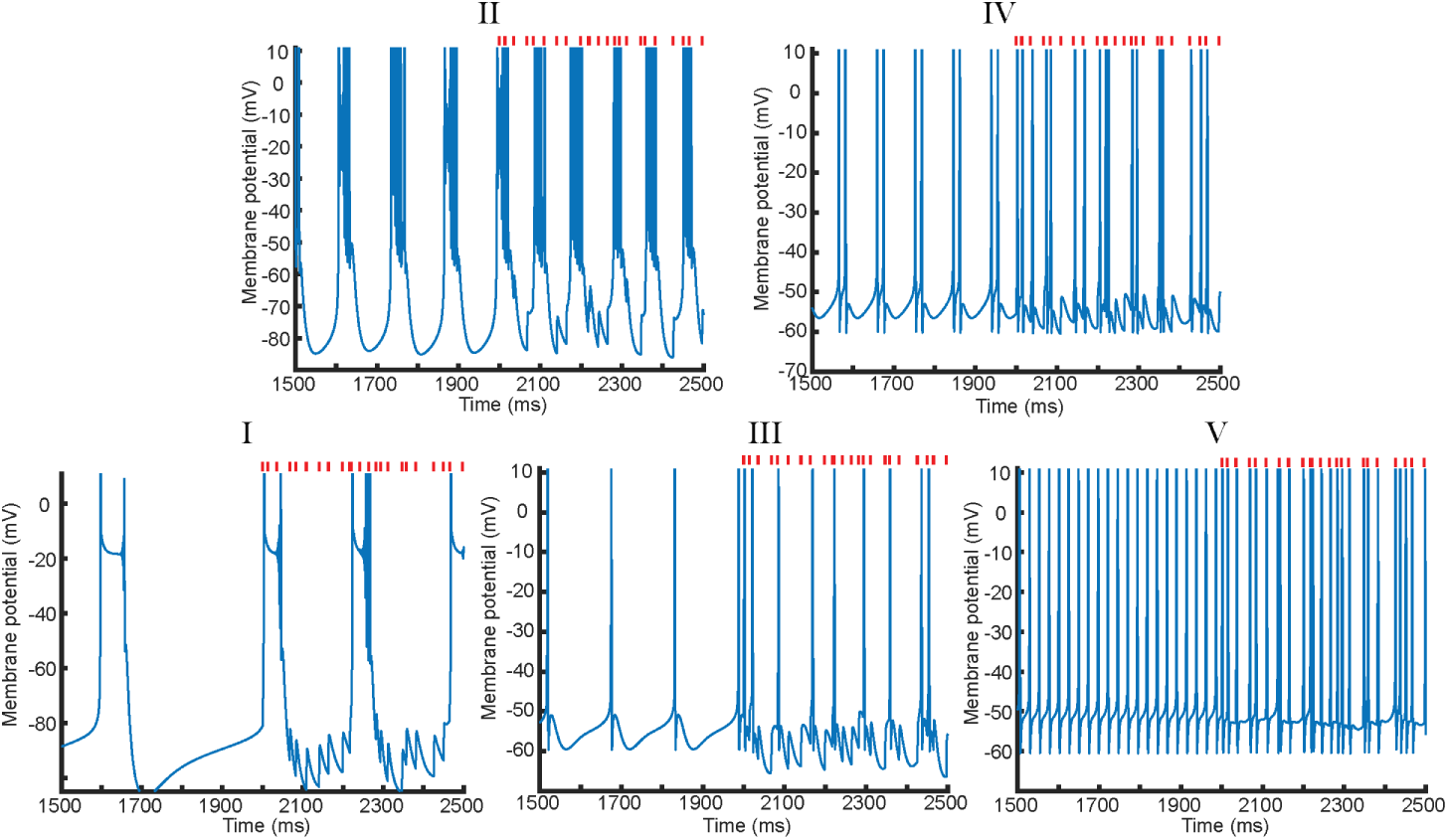
LGN response to retinal input changes with arousal state. The five arousal states match the order and meaning from Figure 1. Each plot shows how LGN spiking (in the 5 different arousal states) responds to an identical retinal input that is turned on at time= 2000ms. Retinal spikes occur at time points denoted by red tick marks. Note, retinal ISIs are drawn from a gamma distribution with a mean of 60 spikes/sec.

The five distinct model arousal states differ in their processing of visual information from the retina. In states I and II, retinal spikes are not patterning the output but instead act as blanket excitation by increasing the low-threshold bursting frequency. This is confirmed when replacing the retinal inputs with a fixed increase in *I_app_* during states I and II yields a similar thalamocortical cell output (Figure 3II and Supplemental Figure 5A and Supplemental Figure 5B). While there is an increase in frequency of the low-threshold bursting, the timing of the retinal spikes is irrelevant for patterning the LGN output as they are not strong enough to transition the model out of the sleep-like states. In contrast, in the natural wakefulness states III and IV (Figure 3) the model enters what we call a filtering mode. The TC cell is responsive to the retinal inputs but only a subset of the retinal spikes are transmitted; as the model progresses from state III to IV there is a continuous increase in retinal transfer. Finally, in the fast tonic firing regime (state V), [18, 57], there is a nearly faithful transmission of retinal spikes regardless of the spike timing. This is corroborated by the accompanying delay time and transfer frequency (between retinal and LGN spikes) statistics: I (8.5ms, N/A), II (9.3ms, N/A), III (3.0ms, .4), IV (3.2ms, .63), and V (2.5ms, .99) (Figure 3). Note, transmission percentages for state I and II are given as N/A as the cell remains in low-threshold burst mode with average delay times greater than 5ms (section 2.5 will provide detailed descriptions of the delay time and transfer frequency metrics [9, 58]).

There are a couple of nuanced points regarding the response of state II to retinal inputs that result from the bistability of the region and the interactions with the Sleep-to-Wake border (Figure 2). First, if parameters are set such that the model is in State II but very close to the sleep to wake border, small amounts of excitation, such as provided by incoming retinal input, may allow the model to transition into State III and thus filter the input (Supplemental Figure 5C and Supplemental Figure 5D). Second, we predict that the quiescent state of region II is more likely to be observed (as it is the stable state found after noise cessation) for low neuromodulatory tone (high *g_kl_*) and closer to the Sleep-to-Wake border (Supplemental Figure 3B and Supplemental Figure 6). This reveals two ideas: first, it provides further evidence for why, in the Crunelli experiments, the quiescent state was observed only prior to agonist application [28]. Second, it suggests that during slice experiments, the use of noisy stimuli could help correctly characterize firing regimes, uncover if there is any type of bistability and decipher stable versus unstable steady states (Supplemental Figure 6).

Figure 3 provides individual representations of the LGN response to retinal inputs in the different dynamic regions. The following subsection explores how variable the thalamic outputs can be, particularly during the filtering states III and IV.

### 2.4 Novel awake dynamic states of the model reproduce experimental variability in mean retinal transmission

In-vivo experimental studies have shown that a given LGN-retinal pair can exhibit transfer frequencies from twenty to one hundred percent [5, 7, 13]. The source of this variability is unknown. This subsection shows that the novel awake states of the model might be a source of such variability.

By examining how the retinogeniculate filtering statistics change throughout the five thalamic dynamic states, Figure 4 reveals that the vast majority of filtering occurs in the awake states III and IV. Figure 4A is a version of the two parameter plot from Figure 2 that depicts the underlying state dynamics; it is there to compare with Figures 4B and 4C, which convey the retinal transmission statistics. Figure 4B shows the ratio of LGN output spikes to incoming retinal spikes. Figure 4C plots the average delay time between an LGN spike and the nearest preceding retinal input. Figures 4B and 4C together determine whether an area in parameter space is filtering or not. To be in this filtering mode requires two conditions based on experimental findings: 1) the ratio of output LGN spikes to input retinal spikes is less than one, and 2) the average delay time between the LGN spike and retinal spike should be under five milliseconds, suggesting a causal relationship between retinal input and LGN spiking [9, 58]. This overlap is depicted in Figure 4D, confirming that filtering is primarily restricted to states III and IV. Figure 4 also confirms that retinal spikes are not patterning thalamic output in states I and II (delay time *>* 5*ms*) and that all retinal inputs are faithfully transmitted in state V (Transfer Ratio ≥ 1 and delay time *<* 5*ms*). Note that there is a small region of filtering to the left of the sleep-to-wake curve (dashed purple line). This is consistent with the results of the previous subsection showing that retinal input near the sleep to wake border can push the model into a filtering state (Supplemental Figure 5).

**Fig 4.**
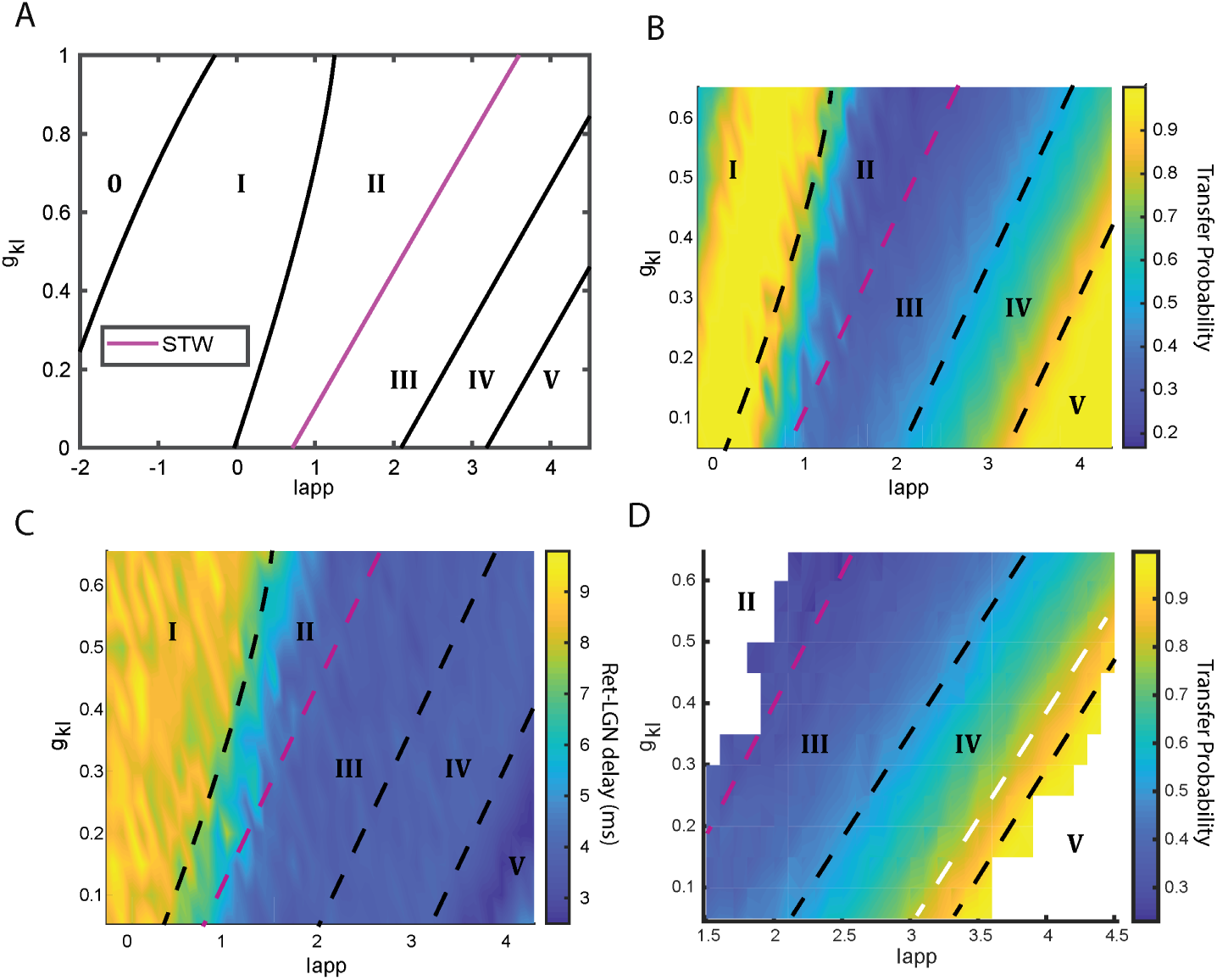
The filtering of retinal signals coincides in parameter space with the novel awake dynamic regimes and captures observed heterogenity in transmission percentages. A) Reproduced diagram from Figure 2 for visual comparison with the other panels. B) Transfer probability (colorbar value) between incoming retinal spikes and outgoing LGN neuron spikes plotted as a function of *I_app_* and *g_kl_*. C) The delay time in milliseconds (colorbar value) between LGN spikes and the nearest driving incoming retinal spike plotted for the same *I_app_*and *g_kl_* parameter space. D) Overlap between B and C where filtering occurs: transfer probability *<* 1 and delay time *<* 5ms. The details for calculating transfer frequency and the delay time are provided in Section 5. The state transition curves (dashed black and purple lines) have been redrawn in 4B, 4C and 4D for further clarity. The white dashed line in 4D denotes where transfer frequency in the filtering mode reaches .75. For experimental comparison with mean retinal transfer efficacy in resting awake animals see Figure 7 of [5].

Of note, our model also reproduces the experimentally observed distributions of retinal transfer ratios [5, 7, 13, 15]. Numerous experimental studies have shown that the average overall transfer efficacy rate is near fifty percent with typical values ranging from twenty to seventy percent [5, 15]. Figure 4D shows that the model produces a similar distribution of transfer ratios: the majority of parameter space (area between the purple and white dashed lines) where filtering occurs is dedicated to transfer frequencies between .25 and .75. Only for the small area of parameter space to the right of the white line in Figure 4D do transfer ratios reach above .75, indicating that observing these high transmission ratios should be rare.

This subsection shows that the model compares well with experimentally observed mean retinal transfer percentages. The next subsections will examine two more granular details of retinogeniculate transmission: what percentage of LGN spikes are driven by the primary retinal ganglion cell (RGC) and which retinal spikes are preferentially transmitted versus filtered out.

### 2.5 High-threshold bursting states III and IV preferentially transmit short retinal interspike intervals similarly to experimentally observed awake animal retinal transfer

Retinogeniculate experimental studies have shown that the majority (*>* 90%) of LGN spikes are driven by a primary retinal ganglion cell [9, 58]. These studies revealed that the driving RGC spike preceded the LGN spike with a delay time between 0ms and 5ms [9, 58]. To compare with these experimental results, Figure 5A plots a histogram of the state IV model delay times between recorded LGN spikes and the immediate closest preceding retinal spike. In agreement with the experimental literature, the majority (90%) of the model delay times are between 0ms and 5ms [9, 58]. LGN spikes occurring more than 5ms after the closest retinal spike (10%), can either be attributed to integration from other non-modeled retinal cells whose inputs are implicitly contained within the excitation term *I_app_* [3, 4], or the underlying intrinsic spiking dynamics from state IV. This finding indicates that retinal input largely is sufficient to override the intrinsic model dynamics and dictate the thalamic spike timing.

**Fig 5.**
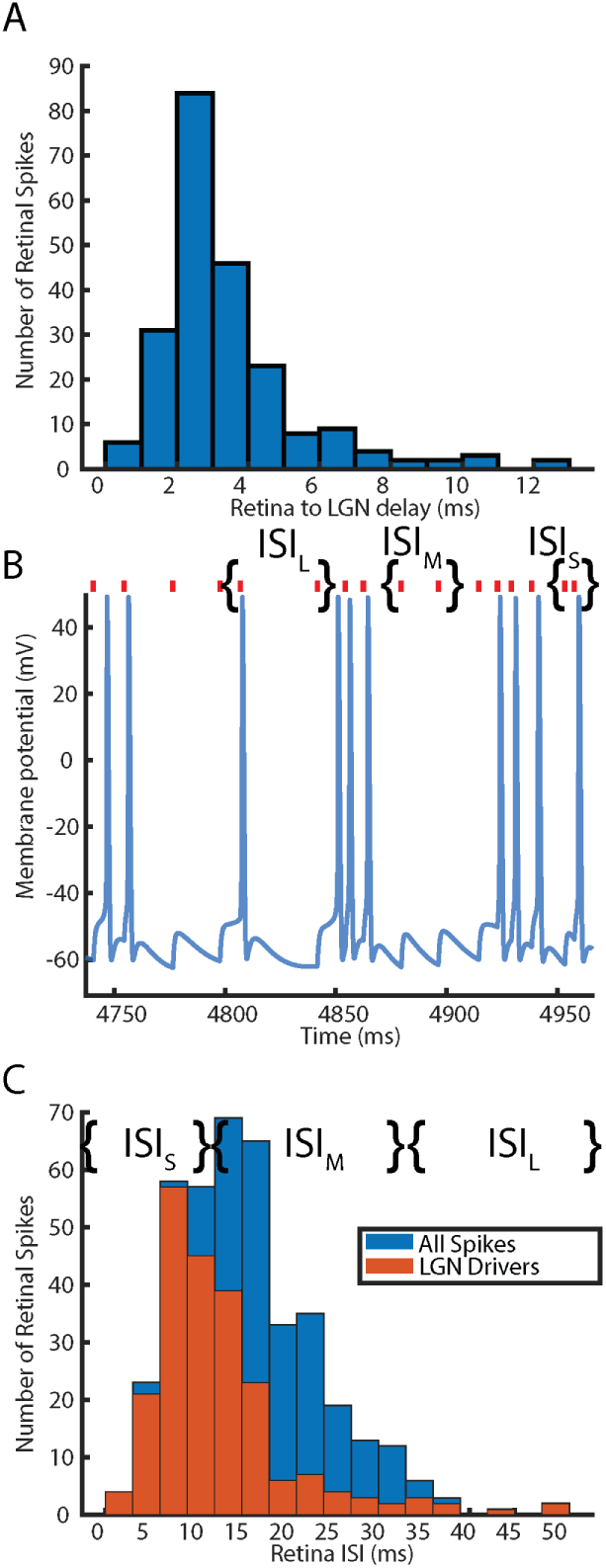
LGN spike generation in the awake state is heavily influenced by retinal ISI. A) Distribution of delay times between each LGN spike and the preceding retinal spike. B) Membrane potential of the LGN TC cell plotted against time. Red tick marks denote timestamps of the incoming retinal spikes. Curly braces depict example retinal ISIs from the three distinct time length categories: short (*ISI_S_*, 0-10ms), medium (*ISI_M_*, 10-30ms) and long (*ISI_L_*, *>* 30ms). C) Distribution of retinal interspike intervals. The ISIs for the full retinal spike train are plotted in blue. The overlapping orange histogram depicts the ISIs between two retinal spikes where the second spike is an LGN spike driver. Curly braces here depict the three distinct time length categories.

It known that the successful transmission of RGC spiking inputs is dependent on ISI length [5, 6, 8, 11–13]. Notably, the increased transmission probability for short ISIs compared to long ISIs is more pronounced in anesthetized animals; ISIs greater than 30ms have a near zero percent chance of generating a spike in anesthetized animals as opposed to between 10 − 40% chance of producing an LGN response in awake behaving animals [5]. The remaining two panels of Figure 5 examine how retinal spike transmission in the model TC cell during state IV depends on retinal ISI. Figure 5B shows a portion of the voltage trace from the TC cell and the corresponding retinal spike times (red circles). Note, Figure 5B labels examples of three distinct retinal ISI categories: short (0-10ms), medium (10-30ms), and long (*>* 30ms). In the accompanying plot of Figure 5C two histograms are overlaid. The first histogram depicts the retinal interspike interval for the full retinal spike train (shown in blue). The second histogram depicts only the interspike intervals such that the second retinal spike drives an LGN spike (shown in orange).

The main takeaway from Figures 5B and 5C is that, in general, only those retinal spikes that come within quick succession of the previous retinal spike (*ISI_S_*) are nearly guaranteed to generate an LGN output [5, 6, 8, 11–13]. Matching experimental observations [5], Figure 5C shows a critical drop off of retinal to LGN spike transmission after the 15ms bin (*ISI_M_*). The histogram in Figure 5C, also shows that in the awake state there can be a high percent transmission of long ISIs (*ISI_L_*) (even if they occur infrequently) [5, 6]. Interestingly, this increased probability of successful transmission happens for retinal spikes near the beta (16-35Hz) frequency range, a set of frequencies implicated in LGN feedforward signaling to cortex [59, 60].

The voltage trace in Figure 5 suggests a partial mechanism for each of the above results. The near faithful transmission of short ISIs and failure of medium length ISIs appears to be at least partially a result of EPSP summation, as noted in previous works studying anesthetized data [3, 7, 8, 11]. In the case of *ISI_L_* the voltage trace in 5B hints that there has been sufficient decay of the membrane potential to steady state such that any input is enough to generate a spike. A natural follow-up is to ask which model components control these awake dynamics, and their subsequent observed relationships with retinal ISIs during retinal transmission. This relationship between retinal ISI, LGN spike probability, and the responsible ionic currents will be the focus of the next results subsection.

### 2.6 Filtering different retinal ISIs depends on the timescales provided by *I_M_* and *I_CaL_*

The first observed relationship between ionic currents and retinal filtering is that *I_M_* is necessary to produce the awake dynamics of states III and IV where filtering is actually possible. While both the potassium leak current (*I_kl_*) and the M current (*I_M_*) (a non-inactivating slowly decaying potassium current) are controlled by neuromodulatory attention and arousal signals [32, 33, 44, 50–53], the M current has a much stronger impact on the existence and transitions between the awake states. This is shown by contrasting the results of Figures 4 and 6, where Figure 6 plots the relationship between dynamic firing regions (Figure 6A) and the mean filtering properties (transfer frequency in Figures 6B and delay time in Figure 6C) as a function of *I_app_*and *g_M_*. Consistent with the findings for *g_kl_*, the majority of parameter space that satisfies the filtering requirements overlaps with states III and IV (Figures 4D and 6D). However, unlike when *g_kl_*is varied, Figure 6A shows that the existence of states III and IV requires a nonzero *g_M_*. This implies that for high acetylcholine (the main modulator of *I_M_*), and thus low *g_M_*, state V becomes the dominant awake dynamic regime, meaning high arousal signaling ensures all inputs are successfully passed regardless of timing. Further differentiating *I_M_* from *I_kl_*, increasing *g_M_*increases the width of states III and IV thus creating a larger range of external excitation values where retinal filtering is possible than was observed for increasing *g_kl_*.

**Fig 6.**
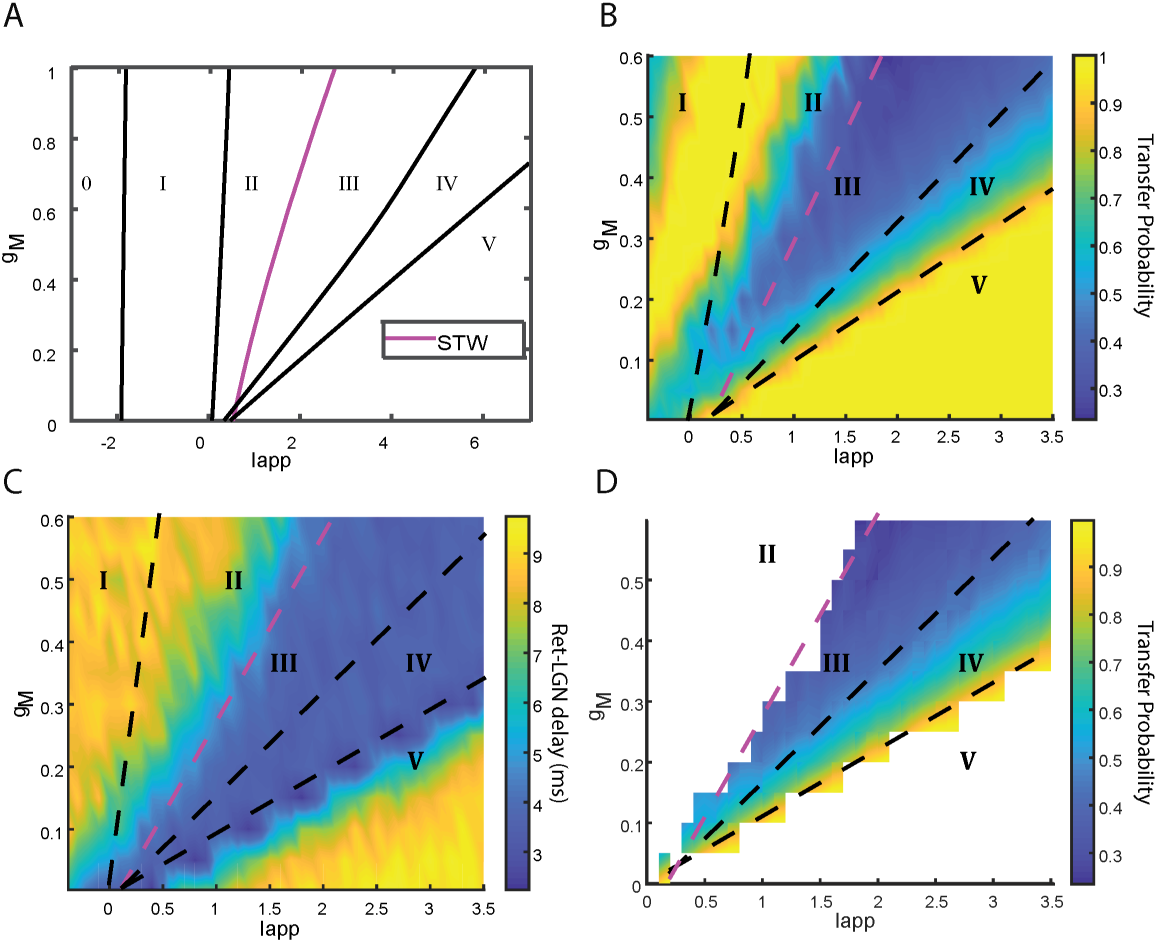
M current is necessary for resting awake state dynamics and corresponding retinal filtering. A) Depiction of the relationship between thalamocortical model state of arousal, cholinergic modulation of the M current conductance (*g_M_*) and external excitation (*I_app_*). State labeling and curve color are identical to Figure 2. Complete bifurcation diagram with labeled mathematical bifurcation curves can be found in Supplemental Figure 7. B) Transfer ratio (colorbar value) between incoming retinal spikes and outgoing LGN neuron spikes plotted as a function of *I_app_* and *g_M_*. C) The delay time (colorbar value) between LGN spikes and the nearest driving incoming retinal spike plotted for the same *I_app_* and *g_M_* parameter space. D) Overlap between B and C where filtering occurs: transfer ratio *<* 1 and delay time *<* 5*ms*. The details for calculating transfer frequency and the delay time are provided in the Methods. Awake state transition curves (dashed black and purple lines) have been redrawn in 4B, 4C and 4D for further clarity.

An examination of **how** the different ionic currents combine to determine the relationships between retinal ISI length and successful transmission reveals that *I_M_* and *I_CaL_*are the most important currents in the awake filtering process. Overlaying the LGN membrane potential with both the timing events of a retinal spike train, and each of the non-spiking, non-background ionic currents (Figure 7A), shows that *I_M_* and *I_CaL_*are the two most active of the four non-spiking currents. In the awake states the M current is always active and responds to all incoming retinal inputs with a rapid increase and a slow decay (Figure 7A). Meanwhile, *I_CaL_* is silent in the absence of inputs, activates if the TC cell crosses the spiking threshold, and inactivates rapidly without additional incoming input (Figure 7A). The other two currents are silent (*I_H_*, a hyperpolarization-activated cyclic nucleotide-gated current) or minimally active (*I_CaT_*(a low-threshold T-type calcium current) with or without retinal input (Figure 7A) due to the sufficiently depolarized membrane potential.

**Fig 7.**
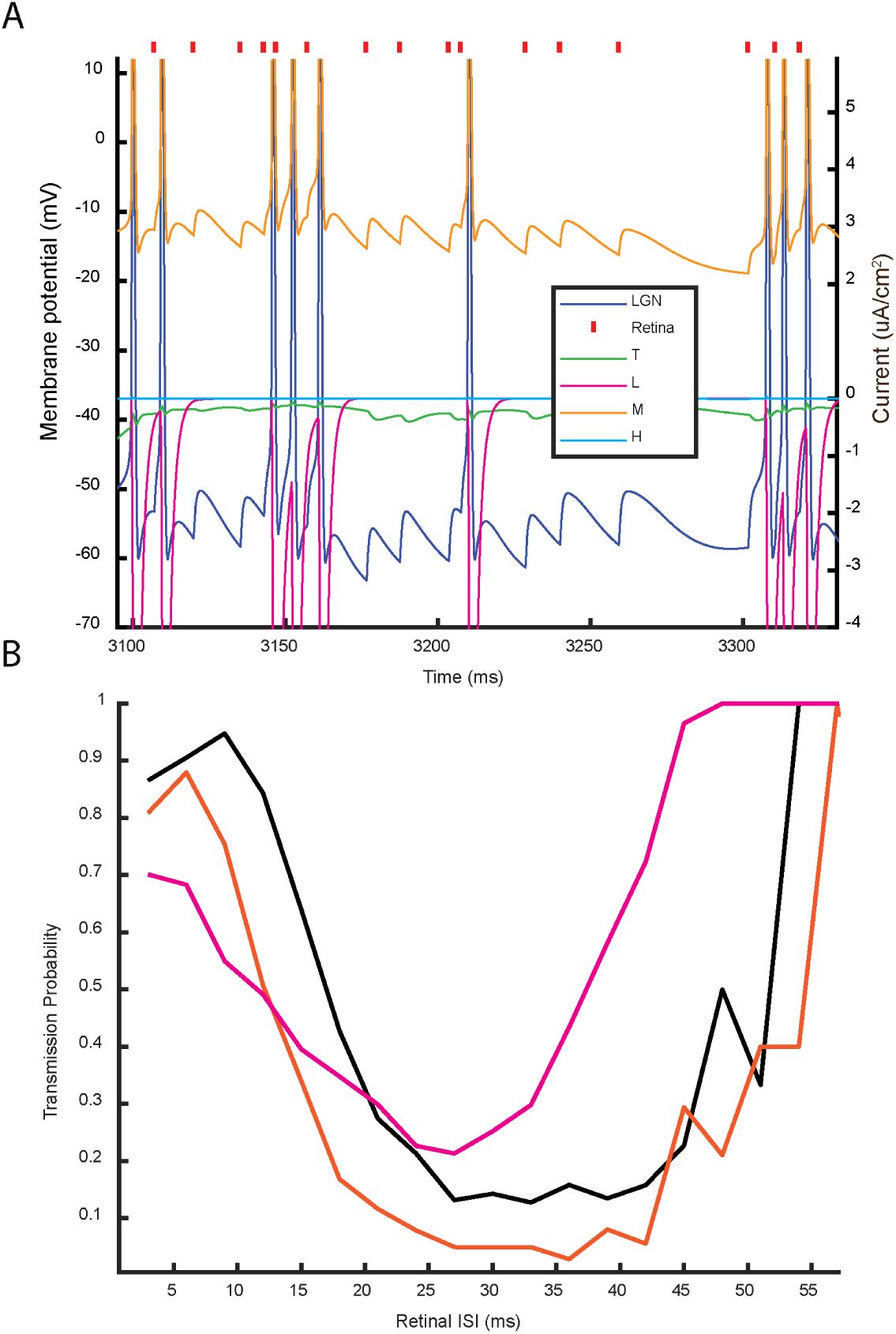
The relationships between key ionic currents and the successful transmission of retinal spikes. A) Membrane potential of the LGN neuron is shown in dark blue with corresponding values recorded on the left axis. Retinal spike times are denoted by red tick marks. The ionic currents *I_CaT_*, *I_CaL_*, *I_M_*, and *I_H_* are plotted in green, magenta, orange, and light blue respectively with values recorded on the right y-axis. B) Changes in the transmission probability of different retinal ISIs as a result of increasing *g_M_* (.12 to .165) (orange) or decreasing *g_CaL_*(.23 to .15) (pink) when compared with the control (black) parameter set’s response. In total 10000 retinal spikes were simulated with ISIs grouped into 3ms bins. Plots here are for a TC cell in state IV, but the same trends are present for the less excited state III (see Supplemental Figure 8 and section 2.9). For comparison with experimental recordings of retinogeniculate transmission probabilities see Figure 9 in [5].

Figure 7 reveals that the inactivation time constant of *I_CaL_*and the decay time of *I_M_* are key factors for determining the transmission probability in the awake state for short and medium length retinal ISIs respectively. At baseline, the TC cell shows a high passage of short (0-10ms) ISIs, a poor transmission of 10ms to 30ms ISIs (medium length), and good transmission probability for retinal ISIs beyond 30ms (long) (Figure 7B, black line). The activation of *I_CaL_* is partially responsible for the high passage of short ISIs while the inactivation time scale of *I_CaL_*dictates that this facilitatory window is between 0-10ms (Figure 7). The poor transmission of medium length ISIs is due to the slowly decaying perpetually active M current decreasing the membrane potential *V* fast enough such that the next incoming excitatory input is insufficient for *V* to cross the spiking threshold (Figure 7). After 30ms, the M current has decayed sufficiently following a thalamic spike such that long retinal ISIs can be transmitted. Increasing *g_M_* decreases the transmission probability for medium length ISIs by at least 20%, and up to 50% while lowering *g_CaL_*decreases the transmission probability for short ISIs by 25 − 40% (Figure 7B). One unexpected result from Figure 7B is that decreasing *g_CaL_* also increases the transmission probability for long ISIs. However, this is simply a byproduct of decreasing short ISI transmission: fewer clustered LGN outputs results in a smaller M-current increase thereby shortening the M-current imposed filtering window. Finally, note that the relationships between currents and filtering properties are consistent across dynamic regions III and IV (See Supplemental Figure 8 and section 2.9).

Combining the results of this subsection with previous ones we conclude that, in the awake state, the ionic currents of the thalamus are capable of filtering out, or relaying, inputs according to their timing. In agreement with experimental literature [5, 7], the thalamocortical cell model in states III and IV filters out inputs received with ISIs between 15-30ms, corresponding to a gamma timescale. However, the model predicts that longer retinal ISIs (*>*30ms) may be successfully passed in these dynamic states. These two observations lead us to hypothesize that the thalamus may pass slower periodic frequencies **and** that faster gamma rhythms may be successfully passed only if they are embedded within a slower frequency oscillation. These investigations will be the subject of the following subsection.

### 2.7 Thalamus entrains to alpha and beta periodic frequencies but theta and gamma are best transmitted as nested oscillations

Upwards of 90% of the synapses made with LGN thalamocortical cells are modulatory, with inputs arriving from layer 6 in V1 cortex, the brainstem, the TRN, and local inhibitory interneurons [3, 60]. Thus the LGN receives a variety of feedback inputs, many of which are oscillatory rhythmic signals normally generated within the above brain areas including: theta (4-8Hz), alpha (8-14Hz), beta (14-30Hz), and gamma (30-100Hz) frequencies [61–64]. Signals occurring at these different frequencies, both as periodic inputs and nested inputs, have been implicated in numerous brain functions including: spatial memory (theta), attention (alpha and theta-gamma), working memory (gamma, theta-gamma) [65–68]. To generate predictions about how the awake thalamus may process non-sensory rhythmic signals occurring at these functional frequency bands, modulatory or otherwise, we stimulate the model with a variety of different rhythmic inputs. We stimulated the TC cell with periodic (5-70Hz) and nested (25-70Hz embedded within 4-12Hz) excitatory and inhibitory rhythmic signals. This subsection focuses primarily on signal processing during the alpha generating awake state IV (as this is the experimentally observed thalamic baseline awake state/rhythm), but will discuss rhythmic signal processing in states III and V at the end.

The model preferentially transmits alpha and beta excitatory periodic frequencies over faster gamma or slower theta/delta (1-7Hz) frequency inputs. Figure 8A shows an example of the thalamus successfully transmitting a high beta (27Hz) frequency input: in fact, inputs between 7Hz and 30Hz result in perfect entrainment (overlap with orange 1:1 line in Figure 8B), albeit with different output firing types (single spikes, doublets, or mixed mode) (Supplemental Figure 9). In contrast, for all frequencies between 30Hz and 70Hz, the ratio of input to output frequency falls beneath the orange 1:1 line, indicating that gamma frequency inputs are not faithfully transmitted. We find that the thalamic output is limited to frequencies below 30Hz in response to any periodic input at frequencies between 30Hz and 70Hz (Figure 8B). Finally, input frequencies in the theta range (4-7Hz) result in a partial 2:1 (output frequency is twice the input frequency) entrainment (overlap with green 2:1 line in Figure 8B), where additional non-stimulus driven spikes cause the output to fire at twice the input frequency, thereby failing to produce a theta rhythm.

**Fig 8.**
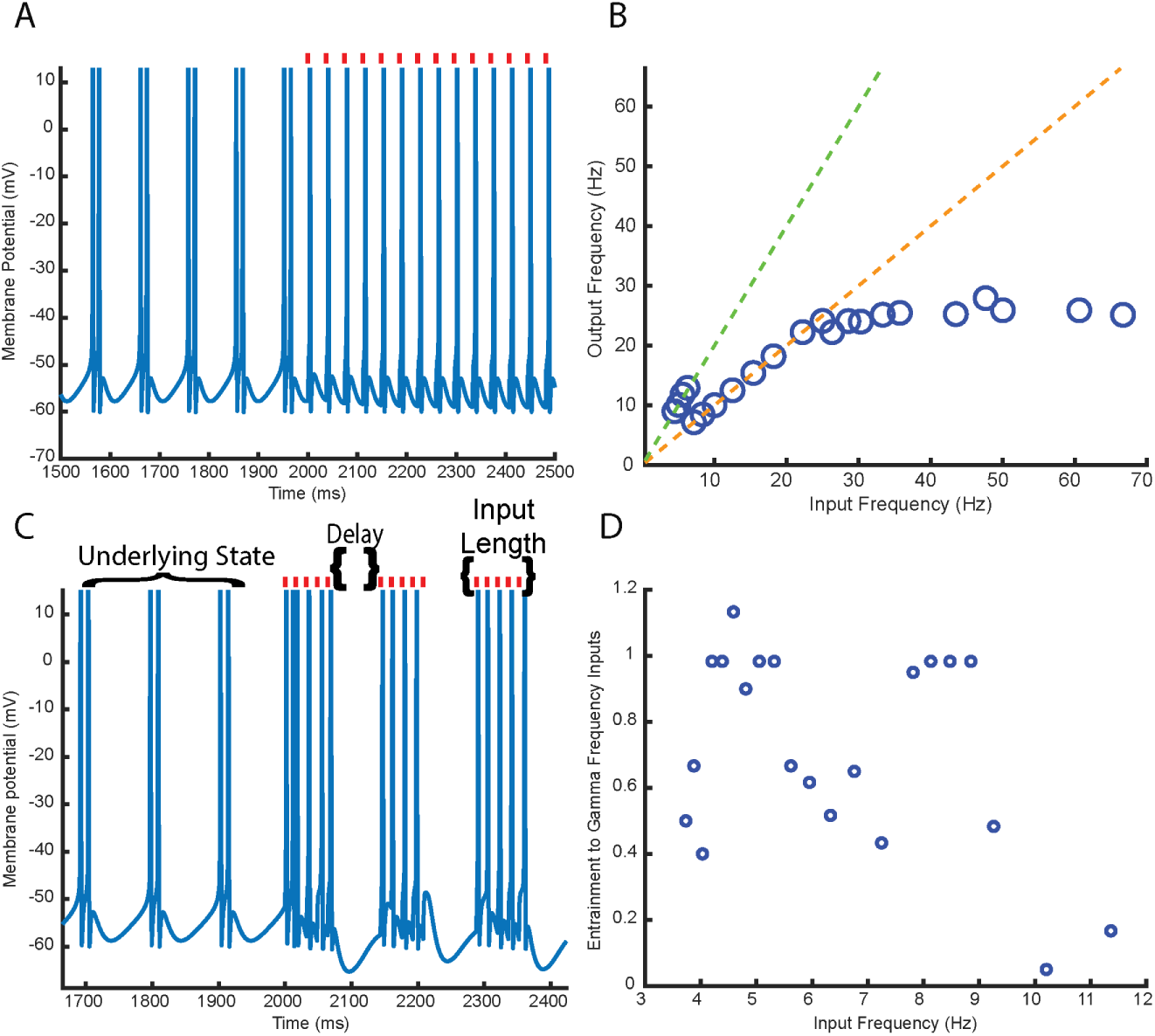
Awake thalamus successfully entrains to periodic excitatory alpha and beta rhythms but passes theta and gamma only as a nested rhythm. A) Sample trace of the TC cell (in state IV) response to a periodic 27Hz spiking frequency input; the thalamus is perfectly entrained to the signal. Red tick marks denote timestamps of incoming excitatory spiking inputs. B) Summary plot showing the number of thalamic burst (or single spike) outputs per second in response to periodic rhythmic excitatory input. Line colors correspond to 2:1 partial entrainment (green) and 1:1 entrainment (orange). C) Sample trace of neuron’s response to a nested theta-gamma (8/66.5Hz) input showing near perfect entrainment to incoming gamma spikes. Entrainment depends on three features that are labeled here: the silent period between the incoming bursts of spikes (labeled as Delay), the combined frequency and number of the incoming fast inputs (labeled as Input Length) and the underlying state of the neuron (labeled as Underlying State). The impact of each of these features is analyzed/discussed within the text. Red tick marks denote timestamps of incoming excitatory spiking inputs. D) TC cell entrainment to the spiking gamma frequency inputs as a function of the incoming slow rhythm frequency. The dependent variable is calculated as the number of TC spikes that match the incoming gamma frequency divided by the number of incoming spikes at the gamma frequency. A value of 1 here indicates 1:1 entrainment to the gamma frequency inputs. Gamma frequency inputs are fixed at 66.5Hz, while the slow timescale (denoted by Delay in panel C) is varied to generate a slow rhythm frequency between 3Hz and 12Hz.

While the model fails to pass excitatory periodic spiking gamma or theta faithfully, the thalamus can transmit these frequencies when they are nested. Figure 8C shows that, in response to an excitatory 66Hz spiking rhythm nested within an 8Hz oscillation, the thalamus entrains to the nested 66Hz signal. Excitatory gamma transmission is most successful when embedded within theta frequency oscillations and shows decreased effectiveness if embedded within alpha (*>* 9) or delta (*<* 4) rhythms (Figure 8D). Interestingly, within the theta frequency band, there are two distinct peaks for perfect 1:1 gamma entrainment found at 4-5.5Hz and 7.5-9Hz (Figure 8D, Supplemental Figure 10). This is noteworthy, as distinctions between high and low theta have been made previously in relation to working memory, behavioral tasks and spatial cognitive processes [67, 69, 70].

Since the thalamus also receives inhibitory feedback via the TRN and local interneurons, we repeat the above analyses for inhibitory signals, starting with periodic inputs. For periodic input frequencies in the range of 5-28Hz, the TC cell produces three distinct forms of entrainment: 2:1 partial entrainment (5-8Hz), 1:1 perfect entrainment (8-16Hz), and 1:2 (output frequency is half the input frequency) partial entrainment (18-28Hz) (Figure 9A and 9B). Although inhibitory and excitatory periodic inputs between 5-28Hz both produce a combination of full and partial entrainment, the thalamic output for inhibitory inputs is limited to between 8Hz and 16Hz. (Figure 9B and Figure 8B). A noteworthy similarity between inhibitory and excitatory periodic inputs over their entrainment ranges is that, as input frequency is increased, the output firing mode changes from multi-spike to single spike (Figure 9A). In both cases, gamma frequencies (*>* 30Hz) are not transmitted successfully in state IV. However, unlike the response to excitatory inputs, increasing the periodic inhibitory input frequency beyond 30Hz results in a sharply decreasing output frequency trending towards full output suppression (Figure 9B). Additionally, the phase relationship between periodic input and output is different for excitatory and inhibitory signals; for 1:1 entrained excitatory signals each input spike always sharply precedes an output spike, whereas the inhibitory signal occurs at varying phases in relation to the output spike, depending on input frequency (Figures 9A and 8B).

**Fig 9.**
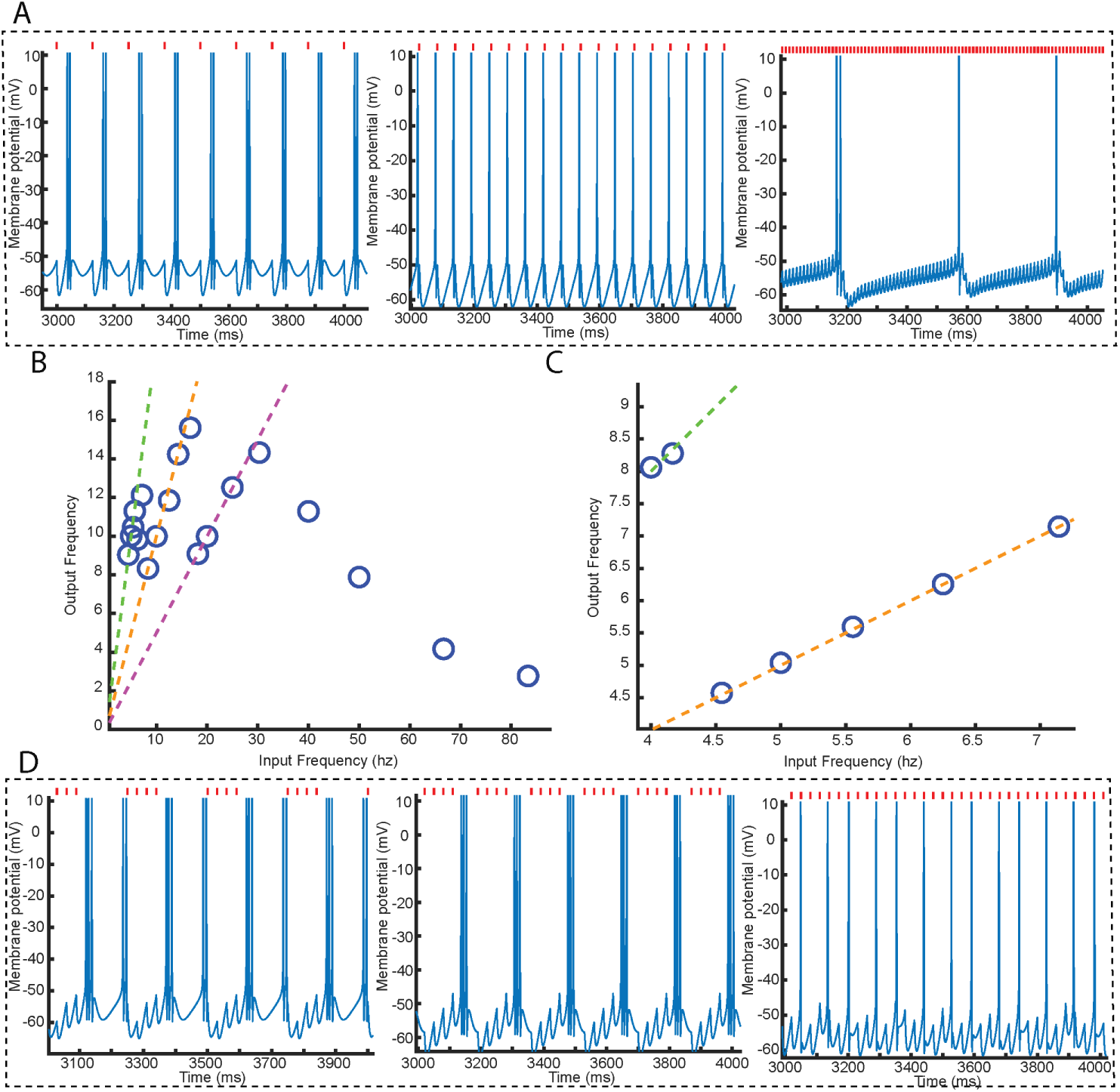
Thalamus preferentially entrains (1:1) to alpha and beta periodic inhibitory rhythms while transforming nested theta-gamma rhythms into a faster frequency nested output. A) Sample responses of a thalamic neuron to periodic inhibitory input: 1:1 entrainment with doublets (8Hz input, left), 1:1 entrainment with single spikes (15Hz input, middle), non-entrained output at a slower frequency (82Hz input, right). Red tick marks denote the timestamps of the inhibitory input spikes. B) Summary plot showing number of thalamic burst (or single spike) outputs per second in response to periodic rhythmic inhibitory input. Line colors correspond to 2:1 partial entrainment where output frequency is twice the input frequency (green), 1:1 pure entrainment (orange), and 1:2 partial entrainment where output frequency is half the input frequency (magenta) C) Number of thalamic input-driven-bursts per second in response to a nested theta-beta frequency. Gamma frequency inputs are fixed at 35Hz, while slow theta timescale is varied. D) Responses of a thalamic neuron to 35Hz inputs nested in a 4Hz theta (left), nested in a 6Hz theta (middle), non-nested for comparison (right).

If inhibitory spiking gamma inputs are coupled to a slow oscillation, the thalamus entrains to the slow frequency but not to the fast frequency. Provided a gamma (33Hz) is nested within a slow oscillation between 4Hz and 8Hz, the TC cell responds to the incoming slow rhythm with 2:1 partial entrainment (4-4.5Hz) or 1:1 entrainment (4.5-8Hz) (Figure 9C and 9D). Unlike with excitatory nested rhythms, when 1:1 perfect entrainment to the slow rhythm occurs, the spiking output is at a much faster frequency than the incoming gamma input (Figure 9D). Furthermore, as was observed with inhibitory periodic gamma frequency inputs, the nested spiking inputs have a different timing relationship to the outputs than was observed with nested excitatory rhythms (Figure 9D); instead of each input spike preceding an output spike, the inhibitory input spikes are grouped and precede the full set of grouped output spikes. Thus, while inhibitory theta nested gamma inputs can be used to entrain to the slow rhythm, the relationship between the incoming gamma spikes and the gamma frequency of the output spikes is vastly different from the response to excitatory nested inputs. Note that, for an identical non-nested gamma (33Hz) frequency input, the TC cell output is a 13Hz alpha rhythm, indicating that nesting an incoming gamma within a slower rhythm is crucial for theta generation (Figure 9D).

While the above simulations for nested rhythms were all run for fixed gamma inputs (4 spikes at 66Hz for excitatory or 4 spikes at 35Hz for inhibitory), faithful transmission of the theta rhythm, with a singular clear nested gamma frequency output, also demands a fast enough spiking nested input. Notably, for excitatory nested inputs, the percent entrainment to the incoming gamma frequency, and percent transmission of total inputs regardless of output frequency, is maximized for faster (100Hz and 66Hz) versus slower (50Hz and 40Hz) gamma frequencies with fewer spikes per slow cycle (Supplemental Figure 11 and Supplemental Figure 12). Inhibitory nested rhythms also require a threshold for the gamma frequency, with entrainment to the theta rhythm occurring only for frequencies faster than 30Hz (Supplemental Figure 12). Interested readers are directed to the supplement for a more detailed analysis of the relationship between the nested gamma input properties and the thalamic output.

Perturbations of the underlying dynamic state suggest that state IV is optimal, when contrasted with states III and V, for entraining to slower (alpha and beta) periodic excitatory rhythmic signals. More specifically, shifting to a faster high-threshold bursting underlying frequency by increasing excitation (moving from state III to state IV through increases of *I_app_* or decreases of *g_kl_*) results in a broader range of excitatory periodic input frequencies that the model can entrain to. Across the entire parameter space devoted to both regions III and IV, the lower bound for entrainment is roughly in the theta band (6-8Hz) until the IV-V border (Figure 10A). However, moving from state III into state IV, the upper bound of entrainment drastically increases from near 15Hz (region III sample point) to near, or above, 25Hz (region IV sample point) (Figure 10B). Note that once the model reaches state V, both the minimum and maximum entrainment frequencies drastically change and are constrained to be in the gamma frequency (30-70Hz), no longer entraining to alpha or beta (Figures 10A and B).

**Fig 10.**
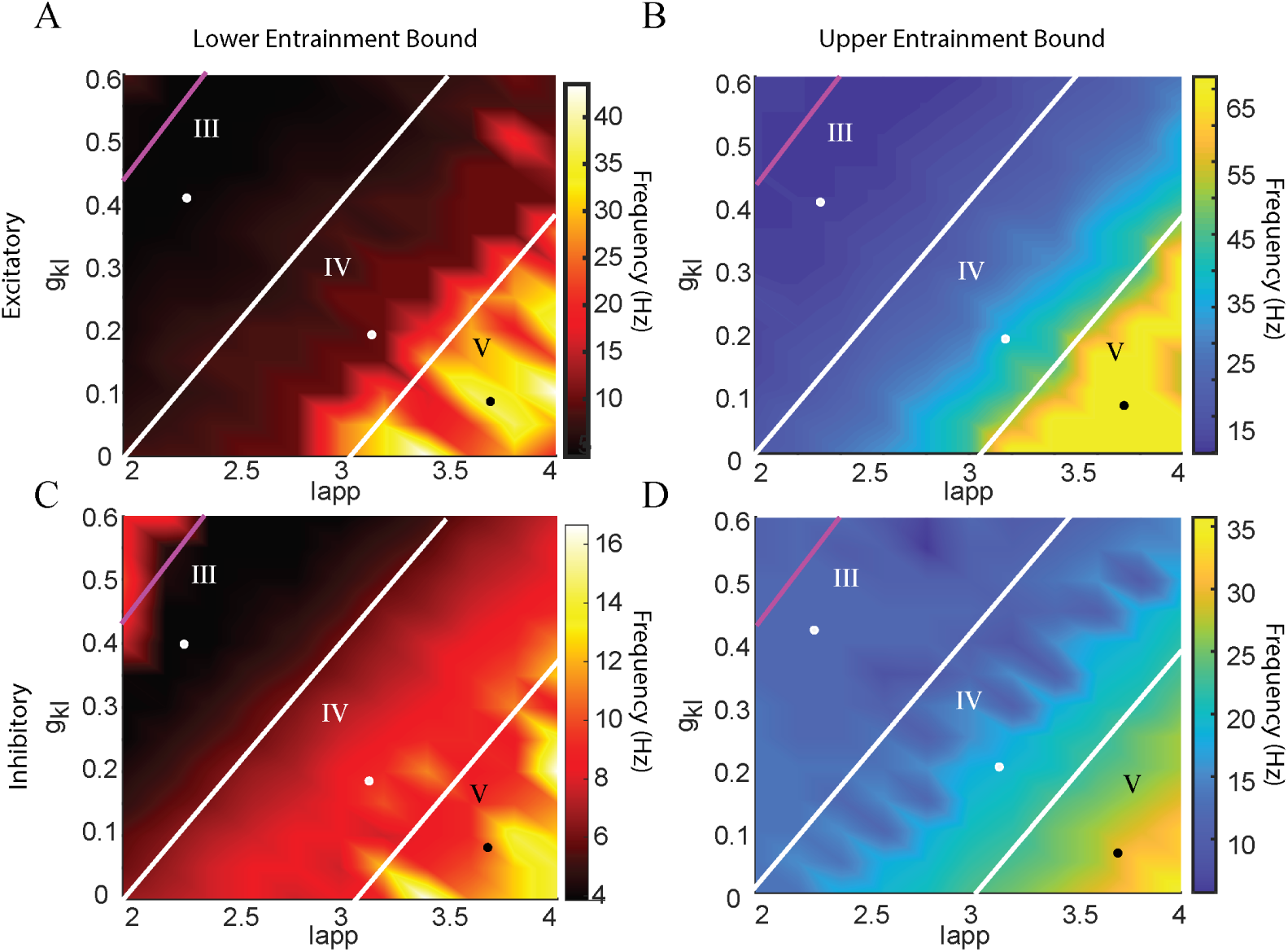
Possible entrainment range widens as the underlying dynamics of the model are moved from state III to state IV. A) Lower frequency limit that the model is capable of entraining to for excitatory periodic inputs. B) Upper frequency limit that the model is capable of entraining to for excitatory periodic inputs. C) Same as A, but for inhibitory periodic signals. D) Same as B, but for inhibitory periodic signals. All entrainment frequencies (given in Hz) are presented as color intensity (depicted on colorbar) for varying values of *I_app_*and potassium leak conductance (*g_kl_*). All awake state regions (III, IV and V) and delineating borders are redrawn for clarity. A representative point in each region is highlighted for ease of description and comparison of the entrainment properties within and between regions. Model response to periodic inputs outside of the entrainment limits in states III and IV can be found in Supplemental Figure 13.

For periodic inhibitory signals across regions III and IV, we find that increasing external excitation (*I_app_*) increases both the upper and lower bound for possible entrainment frequency. The lower bound is increased minimally from ∼ 5HZ (region III sample point) to ∼ 8*Hz* (region IV sample point) (Figure 10C). However, the upper bound is limited to ∼ 10Hz in region III but can reach ∼ 20HZ in region IV (Figure 10D). This implies that an increase in baseline high-threshold bursting frequency (moving from state III to IV) broadens the range of possible entrainment frequencies, critically allowing for the passage of inhibitory low beta signals. Once again, note that upon transition to state V, there is a large shift in entrainment properties: the model no longer entrains to inhibitory theta or alpha frequencies, but can reproduce an incoming inhibitory gamma (Figures 10C and D).

Regarding nested inputs, although all three awake states produce a theta rhythm in response to excitatory and inhibitory theta-gamma inputs (Supplemental Figure 14 and Supplemental Figure 15), there are major differences between the spiking outputs. For instance, in state V, both excitatory and inhibitory nested inputs results in the output of two distinct spiking gamma frequencies, while only one gamma frequency output is generated in states III and IV (Supplemental Figure 14 and Supplemental Figure 15). Furthermore, while inhibitory nested inputs appear to produce a similar output in states III and IV, only state IV successfully transmits all of the incoming excitatory gamma spikes (Supplemental Figure 14).

These results, coupled with the results from previous subsections, imply that arousal state IV, the state producing the baseline awake alpha rhythm, is an ideal sensory filtering state and is also optimal for rhythmic transmission.

### 2.8 Implications for Neuropsychiatric diseases

The L-type calcium current gene CACNA1 is one of many risk genes for a variety of psychiatric disorders including schizophrenia [71, 72]. It has been proposed that mutations of this gene could be partially responsible for the modulation of EEG alpha to theta/delta frequencies in the awake state [73]. Here we explore the effects that a known Single Nucleotide Polymorphism (SNP) mutation of the CACNA1 gene, which increases the L-type calcium channel conductance in induced (reprogrammed stem cells) human neurons [74], might have on the thalamus and thalamic processing.

The model suggests that increasing *g_CaL_* alters the underlying baseline firing properties and eliminates the robust entrainment features of the state IV thalamus. The first observation from Figure 11A is that the increase in L-type calcium conductance changes the baseline high-threshold alpha frequency bursting of state IV into a bursting output occurring at theta frequency. While the theta bursting output has a faster ISI and more spikes per burst in comparison with the high-threshold alpha, it also does not fully resemble low-threshold bursting, implying an intermediate bursting type. This alteration in firing behavior also affects the response of the thalamus to periodic excitatory and inhibitory inputs. While the unaffected state IV TC cell produced 1:1 entrainment to frequencies from 7-30Hz (excitatory) and 8-16Hz (inhibitory) (Figures 8 and 9), here the model fails to entrain to any input above 12Hz (Figure 11B). Furthermore, in the *g_CaL_* altered TC neuron, the output frequencies (y-axis value) are restricted to values between 4Hz and 12Hz, regardless of input type (excitatory or inhibitory) and frequency (Figure 11B). This result suggests that this genetic mutation in the L-type calcium channel gene CACNA1 could lead to a rigidity in thalamic processing that may help explain schizophrenia deficits in cognitive flexibility and task switching [75, 76].

**Fig 11.**
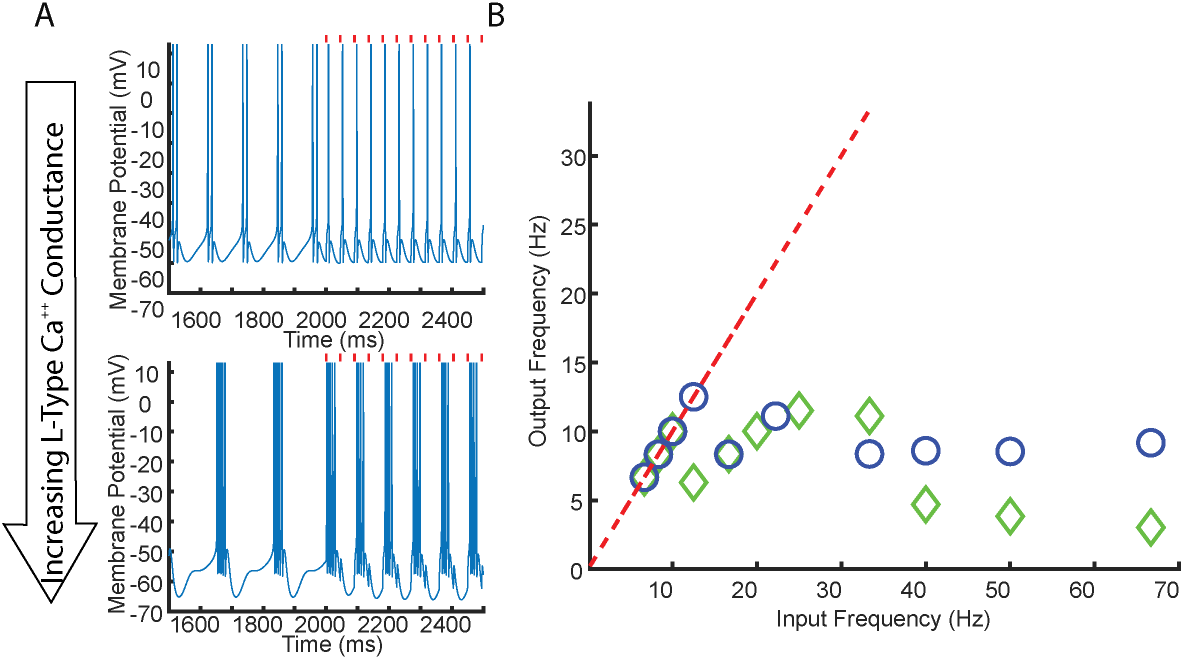
Increases in L-type calcium conductance change the baseline firing type and entrainment properties of the state IV thalamus. A) Membrane potential plots showing the underlying state dynamics and response to periodic excitatory inputs (at 24Hz) for an unaltered thalamocortical neuron (top) and a thalamocortical neuron with increased *g_CaL_* (bottom). External spiking input is turned on after 2000ms, with the timing of the inputs denoted by the red tick marks. B) Summary plot showing number of thalamic input-driven-burst (or single spike) outputs per second in response to incoming periodic rhythmic excitatory (blue circles) or inhibitory (green diamonds) signals. These responses are for a neuron initialized in state IV but with increased *g_CaL_*.

### 2.9 Model is most sensitive to changes in ***I_M_*** and ***I_CaL_***

This section will address model robustness and sensitivity to changes in parameters for states III and IV (high-threshold bursting states), which we have shown emerge as the dynamic working states. Readers interested explicitly in sleep dynamics corresponding to states I and II in out model, and their dependence on parameters, are directed to previous modeling studies [31, 34, 77]. The central result of this analysis is that the alpha high-threshold bursting of the model and its associated functional properties are most sensitive to changes in *I_M_* and *I_CaL_*, currents implicated in neuromodulatory signaling pathways and disease/disorder developments (see previous subsections).

We first investigate how the novel high-threshold bursting properties (burst frequency, ISI, minimum voltage) depend on the conductance values of the four non-spiking, non-background currents (*g_CaL_, g_M_, g_H_* and *g_CaT_*). To do so, we select four model parameters sets from within regions III and IV of the *I_app_* vs *g_kl_* parameter space (to show these results hold for different combinations of excitation and neuromodulation levels) and then independently vary the listed conductances (see Methods).

Figure 12 shows that high-threshold bursting is robust to changes in *g_H_* and *g_CaT_*, but sensitive to changes in *g_CaL_* and *g_M_*. Variations in *g_H_*or *g_CaT_* by 10% resulted in less than a 5% change for all three of the high-threshold burst properties. However, varying *g_M_* by 10% resulted in a near equivalent percent change (with opposite sign) in the high-threshold bursting frequency. Notably, we find the model was most sensitive to changes in *g_CaL_*; a 5% change in *g_CaL_*consistently resulted in a drastic shift in firing properties, or eliminated high-threshold burst firing altogether (Figure 12 and Supplemental Table 1). These results are striking as *I_M_* is modulated during attention and arousal while *I_CaL_* is implicated in the development of neuropsychiatric diseases (see previous subsection). The extreme sensitivity of model properties to changes in *g_M_* and *g_CaL_* suggests that the thalamus tightly regulates these currents to allow for different properties depending on functional context. For example, high sensitivity of the model to *g_M_* and *g_CaL_* may permit alterations of the filtering window or entrainment ranges established by these two currents to flexibly enable the passage of signals which are normally not transmitted.

**Fig 12.**
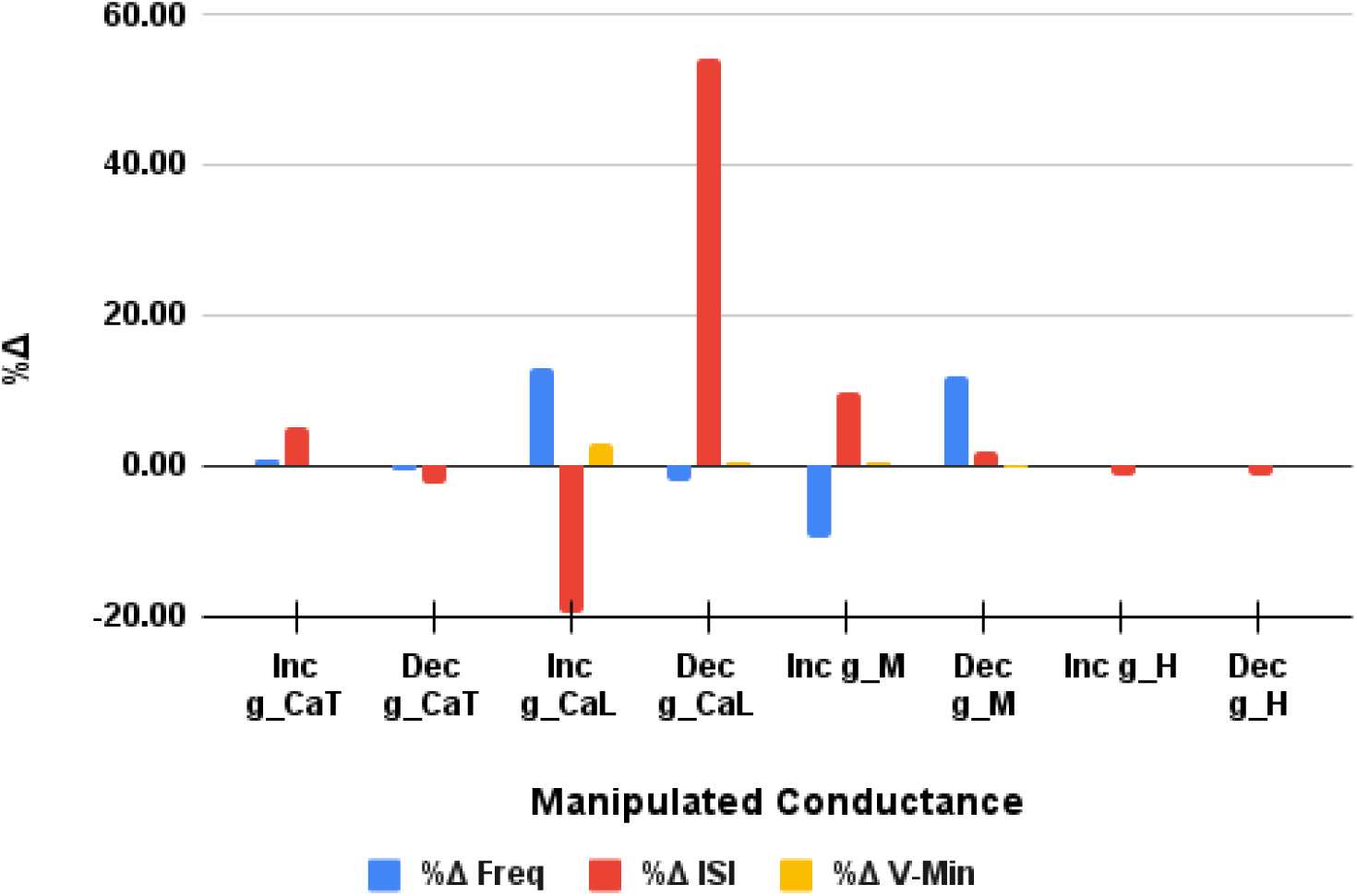
High-threshold bursting exhibits high sensitivity to changes in *g_Cal_* and *g_M_* but not *g_CaT_*or *g_H_*. The average (across four parameter sets) percent change (%Δ) from the recorded value for a control parameter set of each of the three key high-threshold bursting properties (frequency:blue, ISI:red, and minimum voltage:yellow) is plotted as a function of manipulated ionic conductance. Conductances were varied in both directions (labeled here as Inc for increased or Dec for decreased). The full non-averaged data set containing the values for each of the four chosen parameter sets is presented in Supplemental Table 1.

Sections 2.4-2.6 characterize mean retinal transmission properties and how the filtering of different retinal ISIs is influenced by *g_CaL_* and *g_M_*; here we show that these features are robust to changes in *g_CaT_* and *g_H_*, and that the conclusions of Sections 2.4-2.6 hold under different levels of baseline model excitation (different parameter sets from the *I_app_* vs *g_kl_* parameter space are chosen). Supplemental Figure 16A shows that the transmission probability curve for different length ISIs is largely unaltered by changing either *g_CaT_* or *g_H_*. More specifically, under these parameter variations, the model still predicts a high passage of short and long ISIs with a strong filtering out of the medium length retinal spike times. Additionally, although the total transmission probability increases with increased excitation (moving from state III to state IV), the timescales of retinal ISIs preferentially passed through the thalamus are similar throughout regions III and IV of *I_app_*vs *g_kl_*parameter space with the greatest percentage of filtered out ISIs occurring in the 20-40ms range Supplemental Figure 16B. These sensitivity analyses, in tandem with results of the previous subsections, show that all three emphasized model properties explored here (dynamic firing state, retinal filtering, rhythmic entrainment) show some robustness to variations in parameters within states III and IV. We see that regions III and IV both operate, and respond to parameter changes, similarly, with the caveat that a more depolarized neuron (further into state IV) passes (or entrains to) more signals (both rhythmic and non-rhythmic). And finally, across all analyses, we find the model properties are sensitive primarily to changes in the M current and L-type calcium current, further highlighting the importance of these channels for awake thalamocortical signal processing and their possible impacts on disease development.

## 3 Discussion

### 3.1 Summary of results

We constructed a biophysical conductance-based LGN model, with model components constrained using genomic databases [32, 33]. The additions of the M-current (*I_M_*) and the L-type calcium current (*I_CaL_*) to the canonical currents from previous thalamic models [34, 35, 77], were critical for reproducing the full range of experimentally observed LGN firing dynamics. In particular, in the absence of noise, the interaction of *I_M_*and *I_CaL_*produces a firing state of alpha frequency high-threshold bursting. Our model suggests that this dynamic state does not require a special thalamocortical cell subtype as suggested by previous works [18, 35, 36]. In total, the model finds five different firing modes, each of which can be associated with a state of arousal (Figure 1) [18, 37, 38]. Two of these LGN states (I and II) are associated with low-threshold bursting sleep dynamics, the two new states (III and IV) correspond to the theta and alpha rhythms seen from freely moving animals during natural wakefulness [78], and the final state (V) describes the most excited state of focused attention. Importantly, the model transitions between these different states dependent on neuromodulatory arousal signals and/or external excitation (Figure 2).

The model reveals that incoming retinal signals interact with the five thalamic states in different ways; retinal spike train inputs are largely ignored during sleep-like states (states I and II), filtered during the theta and alpha generating awake dynamics (states III and IV), and faithfully transmitted only during the most excited state V (Figure 3). Notably, the area of parameter space where the model exhibits retinal filtering precisely overlaps with the newly modeled dynamic regimes III and IV specific to our model (Figures 4 and 6). Our model’s version of filtering compared with key retinal transfer studies in awake and anesthetized animals shows agreement across the mean and granular properties of retinal transmission (Figures 5) [5, 7, 9, 15]. Our analysis shows that within the novel states III and IV, the model reproduces the experimentally observed heterogeneity in the mean percentages of transmitted retinal spikes. A deeper examination shows that the timescales governing *I_M_* and *I_CaL_* interact with the timing of retinal inputs to specifically control which retinal ISIs are faithfully transmitted in the awake states (Figure 7).

Finally, the model examines how the thalamus might process and interact with non-sensory signals coming from separate brain areas such as the cortex, basal ganglia, and thalamic reticular nucleus. This was done by considering a wide variety of different incoming rhythmic oscillations that have been suggested to play a significant role in inter-area brain communication. Input types investigated include both periodic, and nested, excitatory and inhibitory rhythmic input. Our analysis revealed that in the filtering states thalamic 1:1 entrainment is possible for periodic excitatory and inhibitory inputs at alpha and beta frequencies but that the passage of periodic gamma and theta frequency inputs is limited. Successful transmission of both gamma and theta frequency inputs was achieved when a spiking gamma signal was embedded within a slower theta cycle. Investigating rhythmic signal transmission during the different awake states suggests that state IV is optimal for rhythmic signal processing. We demonstrate that perturbations from this state in manners consistent with different neurological disorders leads to significant decreases in thalamic processing power. For example, we show how an experimentally observed alteration of the L-type calcium current associated with schizophrenia disrupts the awake alpha rhythm thereby limiting the rhythmic entrainment range of the thalamus.

### 3.2 Neuromodulatory inputs are necessary to understand the complete dynamic range of the thalamus

Neuromodulatory signals from the brainstem and cortex have been shown experimentally to be critical in transitioning the thalamus between sleep and waking states [24, 46, 48, 79, 80]. The depolarizing effect of both glutamatergic and cholinergic signaling is modeled here by decreasing potassium conductances, *g_kl_* or *g_M_*, known downstream targets of activated mGlurRs and mAChRs [35, 42, 48]. Interestingly, it has been proposed that these two receptor types have different somatodendritic distributions that could partially account for some of the differences between glutamatergic and cholinergic signaling in the LGN [37]. In our model we find differences in neuromodulatory signaling could be based on the downstream ion channels that are impacted. Although *I_M_* and *I_kl_* both help control the level of depolarization in our model, unlike *g_k_l*, *g_M_* also controls the existence or absence of the alpha high-threshold bursting state and its associated filtering properties. Our results suggest that under different levels of neuromodulatory tone, an identical range of external excitation can span different subsets of the five dynamic states and the transitions among them (Figures 4 and 6). This observation helps provide two possible explanations for why high-threshold bursting was not observed experimentally in some thalamocortical LGN cells [28]: either *g_M_* in such cells is extremely low (due to poor expression or high neuromodulatory tone) such that states III and IV don’t exist for any value of *I_app_*, **or** the strength of the neuromodulatory tone is low enough that states III and IV are not reached by the tested levels of applied current (Figures 4 and 6).

While this work specifically emphasizes cholinergic and glutamatergic signaling, the thalamus also receives a variety of other neuromodulatory inputs including norepinephrine, histamine, serotonin and adenosine signals [18, 35, 81]. Some neuromodulators, such as histamine, reduce *g_kl_*, and thus are expected to exert a similar effect on the LGN as acetylcholine does on *I_kl_* [18, 82]. Neuromodulators can also target other ionic conductances, such as *g_H_* [41]; it has been shown that manipulations of *I_H_* conductance have the capacity to shift the rhythmic firing dynamics during the low-threshold bursting associated with sleep [77]. In total, state dependence on ionic conductance levels suggests that the full characterization of neuronal dynamics seen in vitro, and more importantly in vivo, requires neuromodulator application during experimental studies.

### 3.3 M and L-type calcium current help structure awake retinal transmission

In the natural wakefulness states, the M current works to prevent medium length (10 − 30ms) retinal ISIs from generating LGN spikes while simultaneously allowing some of the longest length retinal intervals (*ISI_l_ >* 30ms) to be transmitted. This implies the LGN in the resting awake state could prevent low gamma frequency inputs (30-80Hz) from being passed while allowing the passage of beta rhythms. These results help suggest a reason for why, in monkey LGN, it was observed that V1-LGN communication happens in the alpha/beta frequency bands [60, 61]. Within the model, we see a high percent passage of these low gamma frequency retinal inputs only when the model has reached the highest arousal state (Figure 3). This indicates that additional excitation (such as increased cholinergic tone or cortical feedback signals) are required to transmit gamma band activity [18], a key EEG signal of increased attention and arousal [40, 59].

The model also suggests that *I_CaL_*may be a key determining factor in the experimentally observed paired spike facilitation (increased probability of successful transmission when the incoming retinal input closely follows a previous input) [7]. Specifically, L-type calcium currents work to permit clustered LGN spikes in response to retinal inputs with the shortest ISIs (0-10ms) with almost 100% faithful transmission (Figure 7). This relationship of retinal ISIs with the L-type calcium current could offer further explanation about a noted difference between awake versus anesthetized animals: the duration of the facilitation effect of retinal spike pairs is shown to be shorter for awake animals compared with anesthetized ones [5]. One possible explanation for this is that the hyperpolarizing effect of anesthetics [83] deactivates the L-type calcium channels, preventing them from helping to pass clustered retinal spikes. Also some of the anesthetics used in retinogeniculate transfer studies, such as propofol, have the capability to selectively block L-type calcium currents [84, 85]. In the model, lowering the L-type calcium conductance (*g_CaL_*) decreases the percent of transmitted short ISIs (Figure 7). This, combined with an unexpected increase in the passage of long ISIs, results in an increased average transmitted ISI (Figure 7). The resulting impact of changing the transmitted retinal spike distribution due to decreased *g_Cal_* is a decrease in paired spike facilitation and a less dependent relationship between short retinal ISIs and successful transmission.

### 3.4 Translating an LGN model to other thalamic nuclei

Although our model is constrained based on available LGN data, the intuition gained here about the distinct dynamic states and their filtering properties may also be relevant for understanding how signals are processed in other thalamic nuclei. This includes investigating how auditory and somatosensory inputs are handled by the medial geniculate and ventral posterior thalamus respectively, as well as deciphering how higher order nuclei such as the pulvinar and mediodorsal thalamus process signals related to more complex cognitive functions like attention [86–88]. We expect similar results when extending (and modifying) our model to non-LGN thalamic nuclei as brain atlas databases [32, 33] confirm the presence of the currents used in our model in other thalamic nuclei across different species. This critically includes the M-current and L-type calcium current which, in the more depolarized states of our model, are responsible for creating dynamic states III and IV, selectively passing tightly clustered inputs and periodic inputs at a beta (or lower) frequency while simultaneously preventing the throughput of tonic low gamma frequency inputs.

However, there are some noted experimental differences between the LGN and other thalamic nuclei to consider for future modeling endeavors. It has been observed that the LGN has a higher density of cholinergic fibers compared to other primary sensory nuclei [89], indicating the relationship between *g_M_*, *g_kl_* and dynamic state may be altered when modeling the auditory and somatosensory thalamus. Furthermore, for translating to higher order nuclei it has been noted that, in comparison to lower order thalamus, these nuclei produce low-threshold bursts in the awake state more frequently, and in some cases hyperpolarize, instead of depolarize, in response to cholinergic signaling [88, 90]. The increased presence of low-threshold bursting in higher order thalamus is particularly interesting as our model indicates that this type of bursting prevents the transmission of non-rhythmic signals. This implies that the low-threshold bursts in higher order nuclei may be acting as a synchronizing or attention signal instead of facilitating active signal processing [88]. This connection between low-threshold bursting and attention was recently examined directly in the pulvinar nucleus using electrical microstimulation, where it was revealed that pulvinar bursting enhanced target detection and broadly increased control of cortical processing and behaviors [91]. This observed importance of low-threshold bursting in higher order cortical control may suggest that the baseline working state for higher order nuclei is state III (from which you can reach the low-threshold bursting state II more easily) and not state IV, as is proposed here for lower order nuclei. In total, the experimentally observed differences between higher order and lower order nuclei indicate that model alterations may be required to properly capture awake processing in higher order thalamus. This may entail adding additional ionic currents, changing the existing model ionic current conductances and/or including different (non-cholinergic/glutamatergic) neuromodulatory signals thereby allowing awake higher order nuclei to still enter state IV and enable the optimal signal processing and filtering predicted to occur there.

### 3.5 Re-framing the awake alpha rhythm

The awake alpha rhythm was the first recorded human brain rhythm and has been observed repeatedly in numerous species [36, 92–94]. The origins of the awake alpha rhythm are still hotly debated. Some studies have attributed the generation of alpha to the cortex [23, 95], others relate awake alpha power to distinct nuclei of the thalamus (such as the LGN) [36], and others consider alpha to arise as an interaction of the two [21, 96]. It remains an open question if and how these mechanisms interact [23]. Our model predicts that in the absence of external inputs the thalamus generates a clear alpha that may be influential in driving a cortical V1 alpha. We also predict that the thalamus will entrain to corticothalamic feedback at alpha frequencies depending on the shape (number of spikes per alpha rhythm peak) of the alpha input (Supplemental Figure 11). In tandem, these predictions suggest a context and signal dependent coexistence of the thalamic and cortical alpha rhythms that could be tested in a future cortico-thalamo-cortical model.

Functionally, alpha rhythms have been hypothesized to: act as an idling state, increase in power with memory load or memory maintenance, exhibit elevated power during task distractor stimuli presentation, have a causal role in the allocation of spatial attention, and be inversely correlated with neuronal spiking and gamma band activity [92, 97–101]. Here we propose a new functional understanding (consistent with the above interpretations) of the thalamic alpha rhythm: this rhythm is a marker that the thalamus is in a working state prepared to filter both rhythmic and non-rhythmic incoming input; the alpha rhythm itself is not explicitly processing, or suppressing, information through the thalamus. Instead, the model state IV (where the baseline rhythm is alpha) is where *I_M_*and *I_CaL_* are interacting to produce filtering and entrainment capabilities; it happens that in the absence of external signals this combination is producing an alpha.

Our model has the following important properties: it produces alpha in the absence of inputs, it exhibits decreasing alpha in response to sensory and non-alpha-frequency rhythmic signals, and it entrains to periodic alpha frequency inputs. These properties are in agreement with experimental observations and can be used to provide mechanistic explanations. Examples are: why the alpha rhythm is largest during periods of eyes closed relaxed wakefulness and is sharply decreased upon eyes opening [102] (an absence of sensory inputs into our model increases alpha); why alpha amplitude decreases when attending to a stimulus [22, 103] (an increase of external inputs to our model decreases alpha); and why alpha is increased during memory maintenance [104] (a top down cortical-produced alpha seen during memory maintenance entrains thalamic output in our model, thereby increasing alpha power).

The model decrease in baseline alpha in response to external signals may have an analogue in the decreases of baseline beta (11-35Hz) power found in macaque [105]. Again, it may not be the frequency that is causal, but the interactions between the currents *I_M_* and *I_CaL_* (possibly with altered time constants or conductance values) that are causal. We therefore predict that manipulations of the conductances and kinetic properties of *I_M_* and *I_CaL_* that disrupt the baseline alpha (or beta) rhythms will also disrupt many of these functional properties currently attributed to these rhythms as well.

### 3.6 Extending beyond retinogeniculate transmission: rhythms and disease

Our model TC cell is capable of generating a variety of experimentally observed rhythms at different frequencies including: low-threshold bursting delta (1-4Hz) and theta (4-8Hz) rhythms during sleep, single spike and high-threshold bursting theta/alpha (4-14 Hz) rhythms associated with natural wakefulness, and fast tonic spiking at beta/gamma (20-80Hz) [18, 27, 31, 37]. Alterations in these rhythms have been associated with diseased states. For example, a number of neurological disorders categorized as thalamocortical dysrhythmias (TCDs), including schizophrenia, Parkinson’s disease, absence epilepsy, tinnitus, and depression have been shown to exhibit decreases in thalamocortical alpha power with a corresponding increase of theta or delta power [21, 106]. While TCD associated disorders span multiple sensory modalities (motor, auditory and visual), thereby involving different thalamic nuclei, the underlying disrupted electrophysiological processes may be similar [107, 108]. In some cases, the increase in the slower rhythms has been associated with increased bursting, changes in HCN channel expression, and/or chronic deinactivation of calcium channels [25, 77, 108].

Our modeling results predict that, by slowing the underlying thalamic frequency from alpha to theta (or delta), neurological disorders will heavily impact the ability of the thalamus to process external stimuli. First, if the thalamus is perturbed towards state III from the alpha bursting state IV, the model exhibits a notable decrease in transmission of non-rhythmic excitatory inputs (Figure 3). Furthermore, in the awake theta generating state III, the model thalamus also displays a decreased ability (compared to state IV) to entrain to both excitatory and inhibitory rhythmic frequencies outside of the theta frequency band (Supplemental Figure 13, Figure 10). This further exacerbates the increase in theta power associated with thalamocortical dysrhythmias [21, 107], as theta becomes the primary thalamic output, regardless of input signal, in this underlying state. We observe a similar response to rhythmic inputs where the thalamic outputs are restricted to the theta band (Figure 11) upon perturbation of the L-type calcium current in a manner consistent with the schizophrenia literature [71, 74]. Limited thalamic flexibility to entrain to rhythmic feedback signals may help explain the observed attention deficits in schizophrenia patients [75, 76, 109] and why L-type calcium channel blockers such as isradipine have displayed success in improving cognitive function in schizophrenia [110]. Finally, if a thalamocortical dysrhythmia perturbs the baseline firing mode all the way to state II (low-threshold bursting at theta), the model predicts a complete inability to process non-rhythmic stimuli. Deciphering which baseline model theta-producing state (state III, L-type perturbed state, or state II) corresponds to the diseased phenotype in question will help uncover mechanisms behind the dysfunction and propose suggestions for restoring the thalamus to its optimal functional state.

The TCD slowing of baseline alpha to theta also affects how the thalamus processes nested oscillations. Coupled with the observed increase in theta, many of these dysrhythmias are also associated with increases in beta and gamma [25, 111, 112]. These observations are in agreement with our model results, as both the resting theta output in state III, and the theta output generated in response to incoming theta nested gamma rhythms, take the form of high frequency bursts (happening at beta and gamma frequencies) nested within theta oscillations (Figures 2 and 9). Interestingly, unlike in state IV (underlying alpha), in state III (underlying theta), when the thalamus is entraining to an excitatory theta nested gamma, the output gamma spikes do not match frequency, or entrain, to the input gamma signals. This inability to entrain to the incoming gamma in state III (the expected state seen during dysrhythmia phenotypes) could further negatively impact the ability of the thalamus to interpret external signals.

Outside of their relationships with TCDs, experimental work within and outside of the thalamus suggests that gamma nested slow oscillations may play an important role in thalamic communication and coordination with other brain areas. Theta-gamma oscillations are most prominently studied in the hippocampus for their positive relationship between cross-frequency-coupling and memory recall [113, 114]. Recent work has also shown evidence that theta-gamma coupling in the anterior thalamic nucleus is important for memory encoding [115], while pulvinar alpha can be coupled to cortical gamma influencing network synchrony and attentional selection [116]. Our modeling results suggest that gamma-nested slow oscillations (theta/alpha) may be possible within multiple thalamic nuclei, and that entrainment is likely dependent on the initial dynamic behavioral state. We find that entrainment success depends on the nested frequencies of the input signal as well as the phase alignment between the underlying dynamics and incoming signals, a feature controlled in state IV by the interactions of the M current and L-type calcium current (Supplemental Figure 10). Interestingly, these two currents are both expressed in brain areas such as the hippocampus and neocortex [32, 33], areas where theta-gamma coupling is prominent and has been attributed to properties of the M-current [117]. In addition to the theta-gamma coupling, in our model the interactions of the M and L-type calcium currents produce excitatory and inhibitory periodic entrainment in the alpha and beta bands; future studies could examine whether the interactions of these expressed currents in the hippocampus and neocortex might display similar results.

### 3.7 Comparison with previous modeling studies

A variety of different conductance-based thalamic models (cellular and network) have been used previously to study rhythmic oscillations under different arousal states. However, these models were limited in their ability to reproduce the full range of awake LGN states seen by the Crunelli group [26, 27]. The pioneering models by Destexhe and McCormick [77, 118] produce only a single tonic spiking awake mode. The later models by both Vijayan and Kopell [35], and Li et. al. [18], created two subtypes of TC cells, one of which could produce the alpha high-threshold bursting state, but missed the slow tonic firing regime. Each of these later high-threshold bursting models hypothesized the existence of a high-threshold T-type calcium current in a specialized thalamocortical cell that still lacks experimental validation. By comparison, our model employs currents that have been validated against genomic databases [32, 33], produces all experimentally found firing types, and correctly transitions between all five arousal states in a manner consistent with changes in external excitation and neuromodulatory signaling [21, 46, 47, 119]. Additionally, contrasted with the Li et. al. model [18], our TC cell, starting in the awake alpha state, exhibits a broader range of entrainment to excitatory frequencies, most notably exhibiting successful passage of the beta frequency band. LGN entrainment to beta is a particularly important feature of the model as beta frequency band feedforward and feedback interactions are essential to LGN-V1 communication [59, 61]. Finally, in this alpha state, our model reveals that TC cells are capable of relaying certain periodic inhibitory rhythms, as well as passing high frequency theta-nested gamma excitatory and inhibitory rhythms, types of rhythmic phenomena not explored in the previous thalamic models [18, 31, 35, 77].

Researchers have also used mathematical modeling to examine retinogeniculate transfer, although these studies have been mostly restricted to using firing rate, integrate-and-fire, or statistical models [8, 11–13]. Collectively, these models confirmed that retinal ISI length was the biggest determining factor in successful retina-LGN transmission in anesthetized animals, and that the process could be approximated with an exponential filter [13]. However, based on the employed model type and data used, these models were not intended to examine retinal spike transmission in awake animals and how it changes with arousal level. This limitation precludes addressing which components of the neuron physiology are responsible for determining the differences in transmitted retinal ISIs when comparing between awake and anesthetized data, namely: the higher average efficacy in wakefulness, the decreased duration of facilitating effects, and the increased transmission of longer retinal ISIs in wakefulness [5]. Here, our detailed conductance-based model addresses these issues by tuning parameter values to control arousal state while simultaneously examining the impact of other ionic conductances on retinal transmission. Specifically, our model framework helps reveal that modulation of the potassium leak and M-current conductances (*gkl* and *g_M_*) controls arousal level and thus the average transfer efficacy, while the timescales of *g_CaL_* (L-type calcium current conductance) and *g_M_*are responsible for the decreased spike facilitation and the increased transmission of longer ISIs in awake animals. One possible extension for this model to consider is the idea of directly modeling extra-retinal inputs instead of considering them to be the action of the parameter *I_app_*. These extensions could help elucidate the role of cortical (or TRN) feedback in retinogeniculate transfer, a process that is still under investigation, but has been discussed in previous works [13] with modeling attempts using some of the simplified frameworks listed earlier [17, 120].

### 3.8 Caveats and limitations

Our model used a minimal subset of expressed thalamic ion channels that can produce the experimentally observed in-vitro dynamics. While this chosen subset by no means exhausts what has been found experimentally [33], or used in previous computational models [18, 31], removing details (currents) we believe to be extraneous for our questions helped create better clarity for understanding what we propose are the underlying mechanisms generating the filtering dynamics. Although the kinetic equations for the channels in our model have been fit to experimental data, in some cases, these data and their corresponding fits, were derived from non-LGN thalamic nuclei [31, 121] or from different brain regions entirely such as the hippocampus, the olfactory bulb, or the cortex [18, 31, 122–125], with recordings performed across a wide range of species. This limitation highlights the need for experimentalists to continue performing in-vitro clamp recording experiments for fitting accurate detailed conductance-based models that produce mechanistic insights and experimentally testable predictions.

In this study we used a cellular model specifically to enable comparisons with the experimental retinogeniculate transfer studies (and their associated data). By using a single cell model, when examining the ability of the model to filter retinal signals, we can only consider postsynaptic effects and how a given incoming retinal spike interacts with the existing active ionic currents. Although this ignores the possible influence of presynaptic effects like facilitation and depression [9], it has been observed that retinogeniculate synaptic amplitude remains constant across different ISI lengths, implying that the presynaptic effects are likely minimal [8].

Previous network models of the thalamus have postulated that alpha-bursting TC cells can phase the outputs of non-bursting cells [30, 35]. Because the alpha bursting cells in these network models provide a driving alpha signal unaffected by feedback signals, we predict that the same results would emerge if our TC cells were used to produce the alpha in these network models instead. However, in general, these alpha producing TC cells are connected with gap junctions and are receiving feedback inhibition (from the TRN and local interneurons) and feedback excitation (from the cortex). These interactions introduce additional time scales that filter or amplify a given signal in a manner distinct from the proposed processing of rhythmic signals we investigated in section 2.7 [3, 13, 29]. To fully explore the impacts of these network level effects would require a network structure.

## 4 Conclusion

Past experimental work has examined two distinct research areas pertaining to the lateral geniculate nucleus: retinogeniculate transmission and the LGN’s role in rhythmic generation and rhythmic processing [3–5, 27]. In this work, we use a novel Hodgkin-Huxley style conductance-based thalamocortical cell model to unite the findings from these two areas of LGN research. Using our model, we propose that the awake alpha rhythmic dynamics are a marker that the thalamus is in a working state that is capable of (and critical for) processing sensory and non-retinal rhythmic inputs. Key features of these processes are determined by the interactions of the M current and L-type calcium current. Furthermore, our model shows that disease driven disruptions of these currents, in experimentally consistent manners, significantly alter thalamic brain state and processing power. These two currents are found ubiquitously throughout the thalamus [31–33, 49], but are lacking (either 1 or both) from commonly cited computational models [18, 31, 35, 77]. The dynamics of these two currents may be instrumental for higher order thalamic nuclei cognitive tasks and could be useful more generally for understanding the mechanisms underlying thalamocortical diseases.

## 5 Model and Methods

### 5.1 Conductance-based model experimental background

To examine properties of retinogeniculate transmission in different states of arousal (i.e., deep and light sleep, natural wakefulness, attentive awake), we need a model that distinguishes between different LGN states. Due to the historical focus of experimental literature [1–3, 77, 126], previous conductance-based computational models emphasized replicating sleep rhythms and their dynamics. These experimental studies prompted the development of the model by Destexhe et al. [77], that is still widely used as a modeling baseline today. Within this model, a low-threshold T-type Calcium current *I_CaT_* and a hyperpolarization-activated cation current *I_H_*combine to produce the low-threshold calcium bursting dynamics associated with sleep rhythms. The remaining currents in the model are the spiking sodium and potassium channels (*I_Na_*, *I_K_*) and two different leak currents *I_l_* and *I_kl_*. While this combination of currents suffices for sleep dynamics, it is not sufficient to capture the complexity of awake firing.

Originally, the awake state LGN (and other thalamic nuclei) were believed to exhibit primarily tonic spiking interspersed with occasional low-threshold bursts [1–4]. While attempting to understand the natural wakefulness associated alpha rhythm in the thalamus [21, 29, 36, 37], experimentalists found a novel awake state firing mode termed high-threshold bursting. In the LGN, it was shown that a subset (10-30 percent) of excitatory TC cells generated this novel form of bursting with an interburst interval in the alpha (8-14Hz) frequency range [21, 26, 27, 36]. Crunelli et al. reproduced this bursting phenomenon in slice experiments after the application of mGluR1 or mAChR agonists [21, 29, 30, 36], that are known to affect the conductances of the background potassium channels [42, 48, 49, 119]. Due to distinct properties of high-threshold bursting (discussed later in the methods), Crunelli et. al. hypothesized that a different (still unknown) mechanism is responsible for this bursting type versus the generators of low-threshold burst firing [27].

Using these experimental results, some computational models have incorporated this novel firing behavior into their investigations about awake thalamic dynamics [18, 35]. However, the lack of clarity surrounding the high-threshold bursting mechanism resulted in key model limitations [18, 35]. Specifically, these models required a phenomenologically constructed calcium current, and were able to replicate only a subset of the full spectrum of awake dynamics [18, 35]. These issues could lead to incorrect dynamic behavior when analyzing and interpreting visual signal processing. Based on these drawbacks, and our intention to study retinogeniculate transmission at different levels of arousal, there was a need for a novel thalamocortical cell model that has experimentally validated currents, and the capability to produce the full breadth of thalamic dynamics.

### 5.2 Model formulation

This subsection presents our single cell lateral geniculate thalamocortical conductance-based mathematical model:

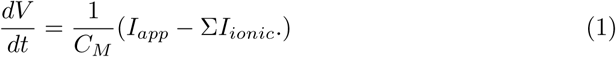

Equation 1 is formulated as a Hodgkin-Huxley style single compartment model [127]. The model dictates that changes in cellular membrane potential *V* are the result of fluxes through membrane bound ionic channels (*I_ionic_*) and external forcing/noise to the cell (*I_app_*) scaled by the inverse of the capacitance *C_M_*. To replicate known experimental observations, and generate experimentally testable and functionally relevant predictions, we constrain the selection of currents for *I_ionic_*using expression data sourced from the Allen Brain Institute and other public databases [32, 33]. More specifically, we look up the gene associated with each ion channel chosen to be included in the model and verify positive expression within the LGN and the thalamus more broadly. It is important to highlight that we are not modeling/including ALL channels expressed within the thalamus, but are instead searching for a minimal subset that reproduces the dynamics of interest.

Validating our selections against this expression data, we arrive at the following minimal model for Σ*I_ionic_*:

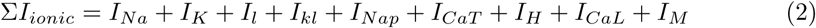

where the specific current kinetics are explicitly detailed in the Supplement (S1 Text). In Equation 2 there are four groups of current types; the first 3 are widely used across previous thalamic modeling studies [18, 31, 34, 35, 77, 128, 129]. First we have the spiking sodium (*I_Na_*) and potassium (*I_K_*) currents necessary to produce action potentials [127]. The model also includes 3 background currents: leak *I_l_*, potassium leak *I_kl_*, and persistent sodium *I_Nap_* to set baseline neuronal excitability levels [18, 127]. Then, to capture sleep dynamics, a the third set of necessary currents, *I_H_* (a hyperpolarization-activated cyclic nucleotide-gated current) and *I_CaT_* (T-type calcium channel), are included [1, 2, 77]. The final two channel types are the non-inactivating Muscarinic potassium current *I_M_*and a high-threshold L-type calcium channel *I_CaL_*. Based on previous modeling studies [34, 43], we hypothesized that these currents could be used to generate high-threshold bursting in the thalamus. Note that in comparison with existing models, *I_M_* and *I_CaL_* are relatively novel additions, and are not ubiquitous across LGN TC cell models [35, 49, 77, 129]. A summary diagram of our TC cell model and the primary purpose for each of the chosen ionic currents is provided in the supplement (Supplemental Figure 17). Note, this selection provides a minimal subset sufficient for capturing our dynamics of interest, but additional currents highlighted in other studies may be useful for future analyses [18, 31, 129].

### 5.3 Model fitting

To address retinal transmission across different arousal levels, the model first needs to properly reproduce the full observed range of experimental firing types. This is accomplished by first running a parameter search algorithm to find possible candidate models that produce the two most distinct firing behaviors: classic low-threshold bursting seen during sleep, and the novel high-threshold bursting phenomenon associated with the awake thalamic alpha rhythm (highlighted by the work of the Crunelli Lab) [28, 36, 37]. Critically, our algorithm requires that the candidate model generates both types of bursting from a single parameter set with variations only to external excitation levels. Good candidate models were determined based on their fit to an objective function that quantitatively described the low- and high- threshold bursting dynamics. Following the completion of the initial parameter search, each candidate model was then subjected to a series of stress tests including: varying *I_app_* to check for the existence of the other dynamic regimes and manipulations of key ionic conductances to replicate pharmacological blocking experiments (Figure 1 and Supplemental Figure 2). The results presented within this manuscript are for one candidate model produced via the parameter search algorithm that successfully reproduced the full set of experimental results of the Crunelli lab [28, 36, 37]. The remainder of this subsection describes the specific details associated with this parameter search algorithm including the parameter set chosen and the constructed objective function.

The parameter search was conducted by using MATLAB’s built in genetic algorithm [130]. This algorithm searches through a bounded, pre-defined parameter space by minimizing a given objective function that we define to have a minimum when the model produces low-threshold bursting for a small value of *I_app_* and high-threshold bursting for a larger value of *I_app_*. The parameter space used here consists of a subset of parameters from Σ*I_ionic_* of Equation 1. Note that the intrinsic ionic currents of Σ*I_ionic_*presented in Equation 2 have the general structure:

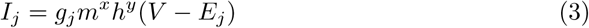

with maximal conductance *g_j_*, gating variables *m* and *h*, and reversal potential *E_j_*. By enforcing that the gating kinetics and reversal potentials to be fixed based on their experimentally fit values from literature [31, 121, 127, 131–133], our parameter space was restricted to the 9 ionic conductance values and *I_app_*. The upper and lower bounds for the conductances *g_j_*set during the parameter search were provided by previous modeling studies of the thalamus and the cortex (in the case of the M-current where previous thalamic modeling studies are lacking) [18, 31, 34].

The objective function that the algorithm minimized to search through parameter spaces was based on work by Marder et. al. (and others) [134–136]. More specifically, the weighted objective function is constructed to quantitatively describe both high- and low-threshold bursting dynamics. Based on experiments by the Crunelli group, three defining bursting characteristics were focused on for the construction of this objective function [27, 28, 37]. The first distinguishing feature is bursting frequency. While there are a variety of different rhythms that have been recorded during sleep (low-threshold bursting) and waking (high-threshold bursting) states [27, 28, 38], two well known frequency bands (delta (1-4Hz) for sleep and alpha (8-14Hz) for wakefulness [18, 38]) are used here. The second characteristic built into the objective function captures how the inter-spike intervals within a burst are different depending on burst type. High-threshold bursts are restricted to have fixed ISI lengths in the range of 10-20ms [27, 30, 36], while low-threshold burst ISIs are more variable and also much shorter [27]. The last piece of our objective function concerns a minimum observed membrane potential. For the awake high-threshold bursting, we restrict the membrane potential to stay above -60mV to agree with the experimental results [36]. This is a critical distinguishing feature as it dictates which currents are involved in the bursting mechanism and separates high-threshold bursting from low-threshold bursting because of the clear hyperpolarization events.

Combining these three descriptors of bursting qualities results in a weighted objective function consisting of 5 components: frequency of low-threshold bursts *F_LT B_*, frequency of high-threshold bursts *F_HT B_*, interspike interval within a low-threshold burst *ISI_LT B_*, interspike interval within a high-threshold burst *ISI_HT B_*, and the minimum observed voltage during high-threshold bursting *MinV_HT B_*. This combination produces the error function given below:

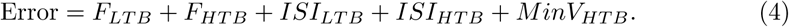

Each term of Equation 4 is constructed such that the objective function is minimized for a window of values for the quantitative feature: 1Hz-3Hz *F_LT B_*, 8Hz-12Hz *F_HT B_*, 5ms-9ms *ISI_LT B_*, 14ms-18ms *ISI_HT B_* and -57mV- -53mV *MinV_HT B_* to enable some flexibility. Specific functional forms can be found in the attached supplemental code.

### 5.4 Retinal spike transmission

Examining the LGN cell model response to visual stimuli required a mathematical formulation of retinal input. A simple spiking retinal ganglion cell (RGC) was synaptically connected to provide fast excitation to the LGN TC cell. An RGC current (*I_ret_* = *s_ret_g_ret_*(*V* − *E_ret_*)) was appended into Equation 1 where *g_ret_* is the strength of the synapse and *E_ret_* its reversal potential. The synapse is controlled by the gating variable *s_ret_*, whose dynamics are adopted from [69] to give:

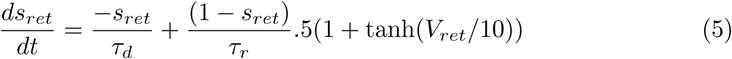

with decay constant *τ_d_*, rise constant *τ_r_*and retinal ganglion cell membrane potential *V_ret_*. For this study, the RGC’s membrane potential is set to a maximum *V_m_ax* on each spike, and then decays according to the following equation:

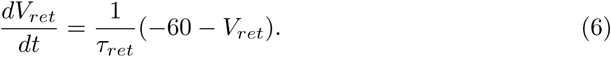

Retinal spike timing in this model was determined by choosing a mean firing rate (60 spikes per second unless noted otherwise) and then sampling spike times from a corresponding Gamma distribution with the given mean [5, 10–12].

Analyzing the input-output relationship between the spiking retinal cell and the LGN output spikes was done by recording a few key experimental metrics. The primary recorded metric is retinogeniculate efficacy (percent trasnmission) defined as the ratio of output LGN spikes to incoming retinal spikes [5, 7–9, 13]. To verify that that LGN spikes are being driven by the incoming retinal spikes the corresponding delay time between LGN spike time and the preceding retinal spike time is recorded. In accordance with experimental literature a RGC is considered to be driving LGN spiking provided the the delay time between spikes is between 0-5ms [7, 9, 13].

### 5.5 Rhythmic signal transmission

The retinal spiking Equations 5 and 6 are repurposed to investigate how the thalamic neuron responds to rhythmic signaling. Rhythmic excitatory feedback inputs are modeled using the fast excitatory synapse employed for the RGC spiking inputs. For inhibitory inputs, the parameters *τ_d_*, *τ_r_* and *E_ret_* were adjusted accordingly to reflect a fast inhibitory *gaba_a_* synapse [69]. Nested rhythmic inputs for excitatory and inhibitory signals are determined by dictating the inter- and intra- burst ISIs as well as the desired number of spikes per input burst.

### 5.6 Sensitivity and robustness analysis

Four representative models were chosen from within regions III and IV of the *I_app_*-*g_kl_* parameter space of Figure 2 to examine feature specific fluctuations in response to parameter variations. For all four chosen models, each of four key parameters (*g_CaL_, g_M_, g_H_* and *g_CaT_*) were varied individually by ±(10% of their original value (±(5% was used in instances where ±(10% destroyed high-threshold bursting) to determine model feature sensitivity. The feature values computed were the frequency of bursting, the ISI of the bursts, and the minimum observed voltage value: the three values emphasized within previous experimental studies for defining high-threshold bursting [27]. For each varied parameter, the percent change from the baseline value exhibited by the control parameter set was computed for all three electrophysiological features.

We also repeated the filtering and entrainment simulations for multiple representative models chosen from throughout regions III and IV of parameter space to show the relationship with underlying state. For retinal filtering, the percent transmission of simulated retinal spike trains for twelve representative models (from the *I_app_*-*g_kl_* parameter space) was recorded as a function of retinal ISI. A similar procedure was used for the entrainment simulations; each different model (according to position in *I_app_*-*g_kl_* parameter space) received the full range of periodic inhibitory and excitatory spike trains. The upper and lower limits of the frequency where pure 1:1 entrainment occurred were recorded. In this case, entrainment was defined to be where the difference between the input frequency and the primary model output frequency (recorded as spikes or burst events per second) was less than one.

### 5.7 Simulation replication and data availability

Except for the two-parameter bifurcation curves, all other simulations were run in MATLAB [130]. The ODE system was solved using MATLAB’s built in numerical solvers. The two parameter bifurcation curves were made using the numerical bifurcation and continuation package in XPP-AUTO [137]. Note that in all bifurcation diagrams (and associated colormap plots) the potassium conductances *g_kl_* and *g_M_* have been scaled (by .06 and .3 respectively) for numerical simulation and visualization purposes.

All code can be accessed at:

https://github.com/keesmcgahan/Thalamocortical-Highthreshold-Bursting-Cell-Model All of the ODEs, current equations, kinetic activation and inactivation rates and additional parameter values can be found in the supplement (S1 Text) as well as within the accompanying code files.

## Supporting information

**S1 Text. Supplemental methods including equations and parameter values.**

**Supplemental Table 1. Changes in recorded values of the frequency, ISI and minimum voltage for high-threshold bursting neurons in response to variations of ionic conductances.** The four varied ionic conductances are *g_CaT_, g_CaL_, g_H_* and *g_M_*. Conductances are increased and decreased by 10 percent (or 5 percent in select instances where 10 percent destroyed HTB) and the resulting bursting metric is recorded. If the row is blank, the neuron stopped producing high-threshold bursts. We investigated four different points in states III and IV (*I_app_*-*g_kl_*) parameter space to generalize across the excitation space.

**Supplemental Figure 1.** T**C cell model smoothly transitions between dynamic states.** Moving from left to right and top to bottom, the TC model (with fixed ionic conductances, *g_kl_* = .2) response to 12 different values of *I_app_* (-2.2, -1.8, -1, -.4, .2, .6, 1, 1.5, 2.0,.2.5, 2.9, 3.5) are presented. The low-threshold bursting frequency continues to increase until there is a sharp transition to the quiescent depolarized state. Then, there is a second transition where the single spiking begins and increases in firing frequency until the firing mode transitions to high-threshold bursting. The frequency of high-threshold bursting then continues to increase until there is a transition to fast tonic firing. Note that there are 5 distinct firing types (that can occur at multiple frequencies) presented here: low-threshold bursting, depolarized quiescence, slow tonic firing, high-threshold bursting, fast tonic firing.

**Supplemental Figure 2. Five different pharmacological application simulations showing how the model TC cell transitions out of high-threshold bursting in manners consistent with experimental findings.** Corresponding experimental manipulations with similar changes to firing type can be found in [30, 36, 37]. A) Baseline high-threshold bursting at alpha frequency for a TC cell starting in state IV (Conductance values from above supplemental methods and *I_app_* = 2.8*µA/cm*^2^) B) High-threshold bursting is removed and replaced with single spiking at a frequency slower than alpha with application of a partial Na+ blocker replicating effect of TTX (*g_Na_* = 3*mS/cm*^2^ and *g_NaP_*= .008*mS/cm*^2^) C) Simultaneous application ofcalcium and sodium blockers prevents the neuron from firing completely (*g_Na_*= 3*mS/cm*^2^, *g_CaL_* = .1*mS/cm*^2^ and *g_NaP_* = .008*mS/cm*^2^) D) Application of an partial L-type Ca++ blocker (*g_CaL_*= .18*mS/cm*^2^) results in a tonic spiking mode at a frequency faster than alpha E) Application of an HCN blocker (*g_H_* = .03*mS/cm*^2^) leaves the firing state of the model unchanged in a high-threshold alpha bursting mode indicating high-threshold bursting occurs at a membrane potentials too depolarized for HCN channels F) Mimicking the action of an mAchR antagonist by increasing leak potassium conductance (*g_kl_* = .018*mS/cm*^2^) transitions the neuron from high-threshold bursting to tonic firing at an alpha frequency.

**Supplemental Figure 3. The complete bifurcation diagram from Figure 2 shows which bifurcations occur in response to changes in *I_app_*and *g_kl_*.** A) Parameter space is separated into 6 distinct regions based on the number and type of solutions that exist. Each distinct region is separated by bifurcation curves that denote where the model undergoes changes in the stability and/or the numbers of solutions to the system of differential equations. The region meaning and curve colors are best understood by traversing through the plot for a fixed value of *g_kl_*and increasing *I_app_*. The corresponding observable model dynamics can be seen in greater detail in panel B. First, in region 0, there is a single stable steady state solution to the ODEs. Moving from region 0 to region I crossed a blue Hopf bifurcation (HB) corresponding to the changing of stability of the steady state solution and the appearance of a new stable periodic orbit/solution. This periodic solution in region I corresponds to the low-threshold bursting dynamics. Moving from region I into region II crosses the second blue (now subcritical) Hopf bifurcation curve. This results in the appearance of an additional unstable periodic orbit as the unstable steady state solution becomes stable. This is what creates the bistability of region II. Note, there is an additional bifurcation curve in region II not shown here which corresponds to a SNP bifurcation (see panel B). Moving from II to III crosses the magenta line (LP/HC) that represents a homoclinic bifurcation curve (meaning that a fold bifurcation and a hopf bifurcation are very close together). This global homoclinic bifurcation triggers the sleep to wake transition and results in the appearance of new periodic orbits (with different amplitudes than the low threshold bursting) where the period of the orbit has grown to infinity but decreases with increasing *I_app_*. In region III the model appears to have 3 solutions, a stable periodic orbit, a second stable periodic orbit corresponding to high-threshold bursting, and an unstable steady state solution. Thus region III exhibits bistability. Crossing the first period doubling yellow curve (PD) from III to IV results in the changing of stability of the non-bursting solution giving region IV a single stable high-threshold bursting solution and an unstable steady state. Finally, crossing the second yellow curve from region IV to region V, eliminates the stable high-threshold bursting periodic solution leading to a single stable periodic orbit and a single unstable steady state solution. B) A one-dimensional bifurcation diagram slice of panel A showing the actual TC cell membrane potential and thus dynamic behavior (recorded as membrane voltage) as a function of *I_app_* with fixed *g_kl_* = .7*mS/cm*^2^ The regions and the corresponding bifurcation points are labeled here according to the similarly colored dashed vertical lines (blue:hopf, magenta:homoclinic, yellow:period doubling). The steady state solutions are the red (stable) and black (unstable) curves. The existence of the periodic orbits is denoted by the green (stable) and blue (unstable) dotted (non-vertical) lines. Note 1: the amplitude of the stable low-threshold bursting periodic orbit of region I/II is not shown by the bifurcation software (XPP) due to the nature of the slow-fast oscillation. Note 2: the periodic orbit stability (green versus blue) labeled in regions III-V corresponds to the non-high-threshold bursting solution. Note 3: There is a bifurcation missing from panel A shown here in region II. This is a Saddle-node-periodic bifurcation (SNP) and corresponds to point where the periodic solution (shown by dotted blue curve) truncates finally eliminating the bistability. Depending on the fixed value of *g_kl_* the SNP bifurcation is closer or further to the magenta curve.

**Supplemental Figure 4. Low-threshold bursting frequency increases continuously moving from state I to state II.** Burst frequency (colorbar value) is recorded as a function of changes in *I_app_* and *g_kl_*. Slower than 1Hz frequencies are obtained at the far left end of region I, which is associated with slow-wave-sleep. The vast majority of region I produces bursting at frequencies between 1-4Hz which corresponds to the LGN Delta rhythm. Finally near and across the I/II border the LGN starts to produce higher than 4Hz bursting frequencies. These higher frequency bursts could correspond light sleep or the type of bursting that is engaged during spindling. As discussed in section 2.2, region 2 is bistable and this frequency of bursting may not be observed without external input or specific engagement of the T-type calcium current which may happen in vivo in response to inhibition. Note that the regions of parameter space and curves labeled here are reproductions of the regions/curves from Figure 2 in the main text.

**Supplemental Figure 5. Retinal input to the TC cell in state II can act like blanket excitation.** A/B) Simulations of the neuron parameterized in region II where the instantiation of retinal input (A) or the static increase of *I_app_* (B) is insufficient to transition the neuron out of a low-threshold bursting mode. Here retinal input does not pattern the LGN output. C/D) Simulations of the neuron parameterized in region II where the instantiation of retinal input (C) or the static increase of *I_app_* (D) is sufficient to switch the neuron out a low-threshold bursting mode. In this parameter regime (C) the model produces time periods of tonic spiking interspersed with low-threshold bursting depending on the retinal sequence. The application of retinal spikes also does not immediately switch the LGN to tonic firing mode: there is a lag delay for sufficient excitatory summation to occur (C). This lag before the switching of firing type is also observed with a simple fixed increase of *I_app_*(D). Across all simulations, prior to 2000ms, the neurons receive no additional input and after 2000ms they receive either random spiking input at time points denoted in red (A/C) or a fixed increase in *I_app_* by .8*uA/cm*^2^ (B/D). The parameter regimes use the standard parameters from the above table but with initial *I_app_* = .8*µA/cm*^2^ and *g_kl_* = .03*mS/cm*^2^ (A/B) or with initial *I_app_* = 1.6*µA/cm*^2^ and *g_kl_* = .03*mS/cm*^2^ (C/D).

**Supplemental Figure 6. Bistability in regions II and III is overcome by noise.** A/B) Response of the neuron in region II starting with two different initial conditions (pink/blue). C/D) Response of the neuron in region III starting with two different initial conditions (pink/blue). All parameter values are standard except *g_kl_*= .03*mS/cm*^2^ and *I_app_*= 1.3, 1.6, 2.1, 2.4*µA/cm*^2^ respectively for A/B/C/D. In all cases (A/B/C/D) the simulation protocol is with no noise (before green line), after the introduction of noise (between green and yellow), after the cessation of noise (between yellow and red), and after the reintroduction of noise (after red). In all instances, regardless of the initial condition and firing mode seen prior to the green line, the firing mode post noise is identical between overlapping plots (blue/pink). Different areas of regions II/III can have different output firing types after the cessation of noise (low-threshold bursting (A)/silent (B) in II, or single spike (C)/bursting (D) in III). The difference in what mode the neuron returns to has to do with the basin of attraction for the given bistable solutions. Initial investigations suggest that quiescence (II) and slow tonic spiking (III) are more likely to be observed close to the STW transition for higher values of *g_kl_*. These results indicate that certain firing types may not be found under certain experimental conditions, since they can only be found in the absence of noise.

**Supplemental Figure 7. The full bifurcation diagram corresponding to Figure 5 in the main text.** The diagram is reproduced in the same manner as the diagram in Supplemental Figure 2. Here we are plotting a two parameter bifurcation diagram using *I_app_* and *g_M_*. The curve color and region meanings are identical to those in Supplemental Figure 2. Although this bifurcation diagram possesses the same types of bifurcations as in Supplemental Figure 2, there is now a meeting point between the homoclinic and period doubling that suggests the possible existence of bifurcations of higher than codimension two.

**Supplemental Figure 8. Detailed depiction of the relationships between key ionic currents and the successful transmission of retinal spikes when the TC cell is in state III.** Changes in the transmission probability of different retinal ISIs as a result of increasing *g_M_* (orange *g_M_* = .165*mS/cm*^2^*, g_CaL_* = .23*mS/cm*^2^) or decreasing *g_CaL_* (pink, *g_M_* = .12*mS/cm*^2^*, g_CaL_* = .15*mS/cm*^2^) when compared with the response of the control parameter set (black, *g_M_* = .12*mS/cm*^2^*, g_CaL_* = .23*mS/cm*^2^). Alterations in ionic conductances reveals that *g_M_* heavily influences the transmission of intermediate ISIs while *g_CaL_* controls the passages of short ISIs. These relationships between ionic conductances and retinal filtering are consistent across the filtering dynamic regions III and IV. In total, 10000 retinal spikes were simulated with ISIs grouped into 3ms bins. For comparison with experimental recordings of retinogeniculate transmission probabilities see Figure 9 in [5].

**Supplemental Figure 9. Three distinct firing types that occur during thalamic 1:1 entrainment to periodic excitatory inputs.** Successful transmission of periodic inputs depends on the level of entrainment. Note, entrainment is measured by comparing the ratio of input frequency to output frequency. The output within a single cycle of a periodic output could be an input-driven burst or a single spike. This means that there may be more output spikes than input spikes, but the neuron could still be considered to be entrained. A an example, during 1:1 entrainment to excitatory periodic inputs the TC cell exhibits three output types: entrained output as doublets (left), entrained output as mixed mode, meaning alternating doublets and singlets (middle), entrained output as single spikes (right). The mixed mode entrainment is interesting as it represents a firing model shift of the TC cell and can result in minor deviations from perfect 1:1 entrainment. The incoming input frequencies are 14Hz (left), 16Hz (middle), and 20Hz (right). Red dots denote timing of excitatory input spikes.

**Supplemental Figure 10. Different theta timescales interact with the underlying alpha dynamics (produced by M and L current time constants) to determine passage of gamma inputs.** The response of the TC cell to theta-gamma nested excitatory inputs depends on the incoming theta frequency (time delay). A-D) Response of the neuron to a nested theta-gamma excitatory signal where the gamma is at 66Hz for all plots and the theta is at 8, 6.5, 5.1 and 4.5Hz for A-D respectively. The gamma input is faithfully transmitted in A, C and D, but there are additional spikes coming from the underlying alpha dynamics appearing in the TC output. The gamma input is disrupted in B where the underlying alpha spikes are appearing in the middle of the gamma timed inputs. E) Summary plot showing the ratio of total input spikes to output spikes as a function of different delay lengths (theta timescales) with a fixed incoming gamma frequency (66Hz) and a fixed number of incoming spikes (4). Notably, the ratio of input to output spikes increases as the delay length between input gamma bursts is increased due to the interaction with the underlying alpha rhythm.

**Supplemental Figure 11. Successful response of the TC cell to incoming nested excitatory gamma rhythms also depends on the number and frequency of the incoming gamma spikes.** A) Sample membrane potential trace of the neuron in state IV responding to 6 incoming gamma (66Hz or 15ms ISI) spikes per theta (6Hz) cycle where the thalamus fails to generate 100 percent passage of spiking. Red dots denote the timing of the input spikes. B) Summary plot describing the relationship between the gamma spikes generated by the thalamus (in state IV) in response to different spike numbers and frequencies. Note the delay length (90ms) here is fixed for all simulations. Blue symbols correspond to the left axis which showcases the ratio of input spikes to output spikes regardless of frequency. Red Symbols correspond to the right axis and denote the ratio of incoming gamma spikes to outgoing gamma spikes at a similar (±3ms ISI) gamma frequency. Different symbol shapes correspond to different speeds of gamma inputs as follows: Triangle 25ms ISI, Asterisk 20ms ISI, Square 15ms ISI, and Diamond 10ms ISI. This figure indicates that increasing the number of spikes per slow cycle, or decreasing the input gamma frequency, decreases the reliability of gamma transmission. In fact, for slower gamma inputs (50Hz or 40Hz), regardless of the number of input spikes per slow cycle, less than fifty percent of the output spikes occur at a matching gamma frequency. As a final note, for all input frequencies tested, fewer spikes per slow cycle helps maximize the total (irrespective of frequency) percent transmission of inputs. Together these results suggest that faster gamma frequencies with fewer incoming spikes per theta cycle, are likely to be relayed better.

**Supplemental Figure 12. Output firing mode of the TC cell in response to nested theta/gamma excitatory and inhibitory inputs depends on the spiking gamma frequency.** The left column presents the TC response to excitatory nested gamma rhythms at 66Hz (A), 50Hz (B), and 40Hz (C). The right column presents the TC response to inhibitory nested beta/gamma rhythms with a fast frequency input at 35Hz (A), 30Hz (B) and 27Hz (C). For a fixed delay length (80ms), but with decreasing gamma frequency, the TC cell spiking output in response to both excitatory and inhibitory nested gammas is: 1:1 entrainment to the theta rhythm with a single gamma frequency (top), 1:1 entrainment to the theta rhythm with an inconsistent gamma spiking output (middle), 1:2 partial entrainment to the gamma input, alternating single spikes and input-driven bursts (bottom).

**Supplemental Figure 13. Transmission of periodic rhythms is generally less successful in states III and V compared to state IV.** A) Number of thalamic input-driven burst (or single spike) outputs per second in response to periodic rhythmic excitatory input for a TC cell in state III. B) Number of thalamic input-driven bursts (or single spike) outputs per second in response to periodic rhythmic excitatory input for a TC cell in state V. C) Number of thalamic input-driven burst (or single spike) outputs per second in response to periodic rhythmic inhibitory input for a TC cell in state III. D) Number of thalamic input-driven burst (or single spike) outputs per second in response to periodic rhythmic inhibitiory input for a TC cell in state V. Red line corresponds to 1:1 entrainment. Periodic stimulation of the neuron in state III entrains to a smaller frequency range than when in state IV (A and C). Periodic excitatory stimulation of the model TC cell in state V can only produce a gamma frequency output, and overrides, or ignores, all lower frequency inputs (B). Periodic inhibitory inputs to the model in state V entrains the neuron over a broad range of frequencies, however, the outputs consist of fast gamma spiking outputs interspersed with minor silent delays (not shown here) for all frequencies. This would imply that even for theta or alpha inputs the TC cell would also produce a very stronger gamma that wouldn’t been seen during state IV entrainment.

**Supplemental Figure 14. Thalamus passes incoming excitatory theta-gamma rhythms best when the underlying dynamics are in the alpha high-threshold bursting mode.** When underlying dynamics are in state III (a slower high-threshold bursting at theta) (left), the output is a theta-gamma but not matching the input gamma frequency. In state IV (alpha bursting) (middle) the thalamus faithfully transmits both the theta and the gamma rhythms. When the thalamus is placed into dynamic state V, the thalamus generates predominately gamma, and not just in response to incoming inputs (right). However, note that in state V, a pause occurs after each input, thereby still producing a theta timescale in the output. Red dots denote the timing of the input spikes.

**Supplemental Figure 15. Thalamic response to incoming inhibitory theta-gamma (6Hz/33Hz) rhythms can differ depending on the initial underlying state.** In response to theta/gamma input, states III-V produce a consistent theta rhythm, but with different spiking responses. In state III, the spiking output alternates between doublet and triplets (Left). In state IV, the spiking output is consistently identical triplets (middle). In state V, the spiking output is a fast series of gamma spikes, followed by a slower set of spikes that occur at a frequency matching the underlying firing dynamics (right). Red dots denote the timing of the input spikes.

**Supplemental Figure 16. Retinal filtering is robust to changes in *g_H_* and *g_CaT_* and produces a similar filtering window across regions III and IV *I_app_* vs *g_kl_* excitation space.** A) Transmission probability as a function of retinal ISI plotted for the control parameter set and 4 variations of this parameter set which are: 10% decrease in *g_CaT_* (purple), 10% increase in *g_CaT_* (green), 10% decrease in *g_H_* (blue), and 10% increase in *g_H_*(yellow). The four variations of the control parameter space produce a similar transmission probability curve. B) Transmission probability as a function of retinal ISI plotted for 12 distinct parameter sets from within regions III and IV of *I_app_*vs *g_kl_*parameter space. Parameter sets were found by fixing a value of *g_kl_*and then taking four .5 increments of *I_app_*. This was repeated for 3 separate values of *g_kl_* yielding the total of 12 parameter sets. Changes in *I_app_* can be counteracted by changes in *g_kl_*, but throughout the entirety of the parameter space tested the 20-40ms window of retinal ISIs are the most likely to be filtered out as described in the main text. The lowest (smallest transmission probability) two groupings of curves (by line type) are from region III and the higher two groupings of curves are from region IV in parameter space.

**Supplemental Figure 17. Schematic depicting TC model structure and the contributions of the included ionic currents** A single cell, single compartment neuron model that has 9 ionic current types. These ionic currents are separated into 4 broad categories based on their primary functional role within the model: Spiking (*I_Na_* and *I_K_*), Low-threshold bursting (*I_CaT_* and *I_H_*), High-threshold bursting (*I_M_*and *I_CaL_*) and Leak/background (*I_kl_*, *I_l_* and *I_NaP_*).

