## Supplemental Document for "Awake alpha bursting emerges as the dynamic working state in a lateral geniculate thalamocortical cell model"

### Supplemental Results and associated Figures

#### Supplement for Section 2.1

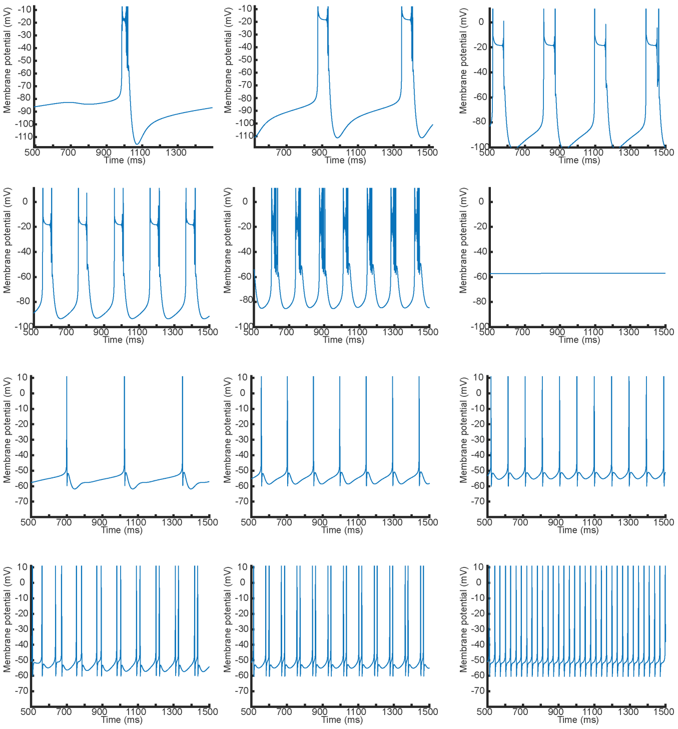

Supplemental Figure 1: **TC cell model smoothly transitions between dynamics states.** Moving from left to right and top to bottom, the TC model (with fixed ionic conductances, $g_{kl} = .2$) response to different 12 different values of $I_{app}$ (-2.2, -1.8, -1, -.4, .2, .6, 1, 1.5, 2.0,.2.5, 2.9, 3.5) are presented. The low-threshold bursting frequency continues to increase until there is a transition to the quiescent depolarized state. Then there is a second transition where the single spiking begins and increases in firing frequency until the firing mode transitions to high-threshold bursting. The frequency of high-threshold bursting then continues to increase until there is a transition to fast tonic firing. Note that there are 5 distinct firing types (that can occur at multiple frequencies) presented here: low-threshold bursting, depolarized quiescence, slow tonic firing, high-threshold bursting, fast tonic firing.

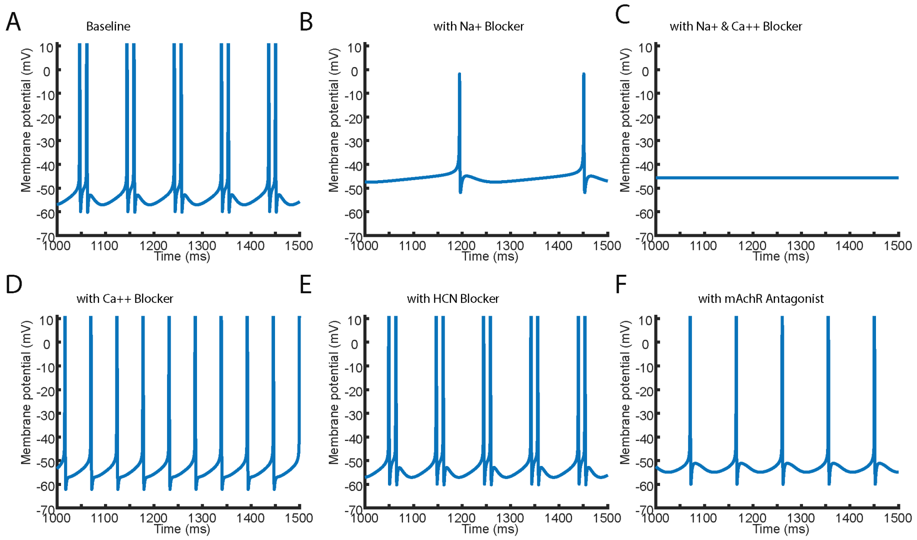


Supplemental Figure 2: **Five different pharmacological application simulations showing how the model TC cell transitions out of high-threshold bursting in manners consistent with experimental findings.** Corresponding experimental manipulations with similar changes to firing type can be found in [1, 2, 3]. A) Baseline high-threshold bursting at alpha frequency for a TC cell starting in state IV (Conductance values from above supplemental methods and $I_{app} = 2.8\mu A/cm^2$) B) Bursting is removed and replaced with single spiking at a frequency slower than alpha with application of a partial Na+ blocker replicating effect of TTX ($g_{Na} = 3mS/cm^2$ and $g_{NaP} = .008mS/cm^2$) C) Simultaneous application of calcium and sodium blockers prevents the neuron from firing completely ($g_{Na} = 3mS/cm^2$, $g_{CaL} = .1mS/cm^2$ and $g_{NaP} = .008mS/cm^2$) D) Application of an partial L-type Ca++ blocker ($g_{CaL} = .18mS/cm^2$) results in a tonic spiking mode at a frequency faster than alpha E) Application of an HCN blocker ($g_H = .03mS/cm^2$) leaves the firing state of the model unchanged in a high-threshold alpha bursting mode indicating high threshold bursting occurs at a membrane potentials too depolarized for HCN channels F) Mimicking the action of an mAchR antagonist by increasing leak potassium conductance ($g_{kl} = .018mS/cm^2$) transitions the neuron from high-threshold bursting to tonic firing at an alpha frequency.

#### Supplement for Section 2.2

This supplement subsection is intended for readers interested in the underlying mathematics used to define the different firing types (dynamic regimes) of the model.

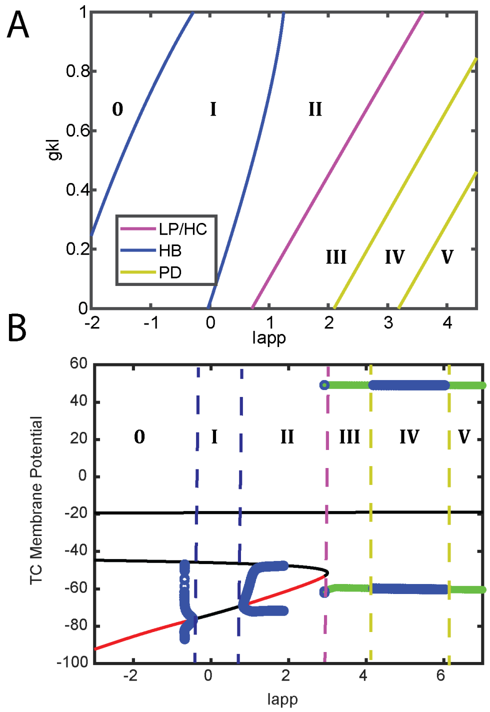


Supplemental Figure 3: **A) The complete bifurcation diagram from Figure 2 shows which bifurcations occur in response to changes in $I_{app}$ and $g_{kl}$.** Parameter space is separated into 6 distinct regions based on the number and type of solutions that exist. Each distinct region is separated by bifurcation curves that denote where the model undergoes changes in the stability and/or the numbers of solutions to the system of differential equations. The region meaning and curve colors are best understood by traversing through the plot for a fixed value of $g_{kl}$ and increasing $I_{app}$. The corresponding observable model dynamics can be seen in greater detail in panel B. First, in region 0, there is a single stable steady state solution to the ODEs. Moving from region 0 to region I crossed a blue Hopf bifurcation (HB) corresponding to the changing of stability of the steady state solution and the appearance of a new stable periodic orbit/solution. This periodic solution in region I corresponds to the low-threshold bursting dynamics. Moving from region I into region II crosses the second blue (now subcritical) Hopf bifurcation curve. This results in the appearance of an additional unstable periodic orbit as the unstable steady state solution becomes stable. This is what creates the bistablity of region II. Note, there is an additional bifurcation curve in region II not shown here which corresponds to a SNP bifurcation (see panel B). Moving from II to III crosses the magenta line (LP/HC) that represents a homoclinic bifurcation curve (meaning that a fold bifurcation and a hopf bifurcation are very close together). This global homoclinic bifurcation triggers the sleep to wake transition and results in the appearance of new periodic orbits (with different amplitudes than the low threshold bursting) where the period of the orbit has grown to infinity but decreases with increasing $I_{app}$. In region III the model appears to have 3 solutions, a stable periodic orbit, a second stable periodic orbit corresponding to high-threshold bursting, and an unstable steady state solution. Thus region III exhibits bistability. Crossing the first period doubling yellow curve (PD) from III to IV results in the changing of stability of the non-bursting solution giving region IV a single stable high-threshold bursting solution and an unstable steady state. Finally, crossing the second yellow curve from region IV to region V, eliminates the stable high-threshold bursting periodic solution leading to a single stable periodic orbit and a single unstable steady state solution. **B) A one-dimensional bifurcation diagram slice of panel A showing the actual TC cell membrane potential and thus dynamic behavior (recorded as membrane voltage) as a function of $I_{app}$ with fixed $g_{kl} = .7mS/cm^2$** The regions and the corresponding bifurcation points are labeled here according to the similarly colored dashed vertical lines (blue:hopf, magenta:homoclinic, yellow:period doubling). The steady state solutions are the red (stable) and black (unstable) curves. The existence of the periodic orbits is denoted by the green (stable) and blue (unstable) dotted (non-vertical) lines. Note 1: the amplitude of the stable low-threshold bursting periodic orbit of region I/II is not shown by the bifurcation software (XPP) due to the nature of the slow-fast oscillation. Note 2: the periodic orbit stability (green versus blue) labeled in regions III-V corresponds to the non-high-threshold bursting solution. Note 3: There is a bifurcation missing from panel A shown here in region II. This is a Saddle-node-periodic bifurcation (SNP) and corresponds to point where the periodic solution (shown by dotted blue curve) truncates finally eliminating the bistability. Depending on the fixed value of $g_{kl}$ the SNP bifurcation is closer or further to the magenta curve.

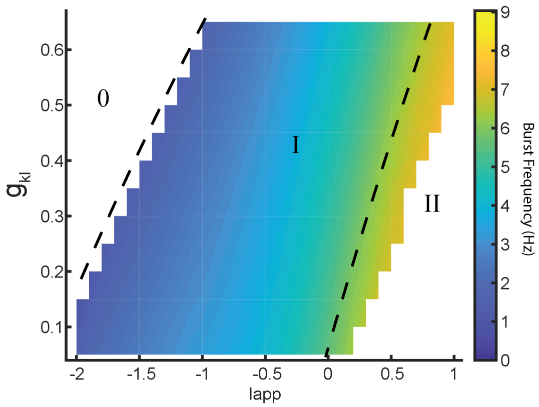


Supplemental Figure 4: **Low-threshold bursting frequency increases continuously moving from state I to state II.** Burst frequency (colorbar value) is recorded as a function of changes in $I_{app}$ and $g_{kl}$. Slower than 1Hz frequencies are obtained at the far left end of region I, which is associated with slow-wave-sleep. The vast majority of region I produces bursting at frequencies between 1-4Hz which corresponds to the LGN Delta rhythm. Finally near and across the I/II border the LGN starts to produce higher than 4Hz bursting frequencies. These higher frequency bursts could correspond light sleep or the type of bursting that is engaged during spindling. As discussed in section 2.2, region 2 is bistable and this frequency of bursting may not be observed without external input or specific engagement of the T-type calcium current which may happen in vivo in response to inhibition. Note that the regions of parameter space and curves labeled here are reproductions of the regions/curves from Figure 2 in the main text.

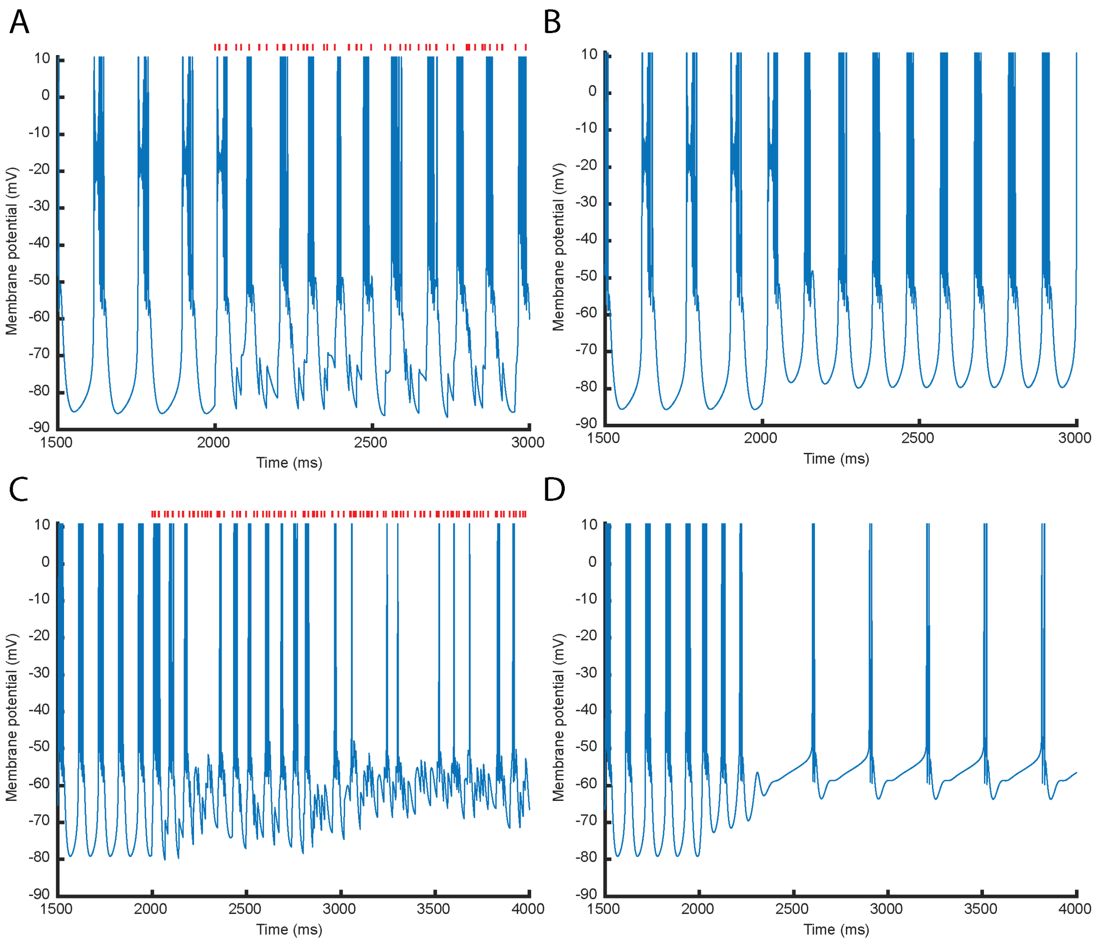


Supplemental Figure 5: **Retinal input to the TC cell in state II can act like blanket excitation.** A/B) Simulations of the neuron parameterized in region II where the instantiation of retinal input (A) or the static increase of $I_{app}$ (B) is insufficient to transition the neuron out of a low-threshold bursting mode. Here retinal input does not pattern the LGN output. C/D) Simulations of the neuron parameterized in region II where the instantiation of retinal input (C) or the static increase of $I_{app}$ (D) is sufficient to switch the neuron out a low-threshold bursting mode. In this parameter regime (C) the model produces time periods of tonic spiking interspersed with low-threshold bursting depending on the retinal sequence. The application of retinal spikes also does not immediately switch the LGN to tonic firing mode: there is a lag delay for sufficient excitatory summation to occur (C). This lag before the switching of firing type is also observed with a simple fixed increase of $I_{app}$ (D). Across all simulations, prior to 2000ms, the neurons receive no additional input and after 2000ms they receive either random spiking input at time points denoted in red (A/C) or a fixed increase in $I_{app}$ by $.8uA/cm^2$ (B/D). The parameter regimes use the standard parameters from the above table but with initial $I_{app} = .8\mu A/cm^2$ and $g_{kl} = .03mS/cm^2$ (A/B) or with initial $I_{app} = 1.6\mu A/cm^2$ and $g_{kl} = .03mS/cm^2$ (C/D)

#### Supplement for section 2.3

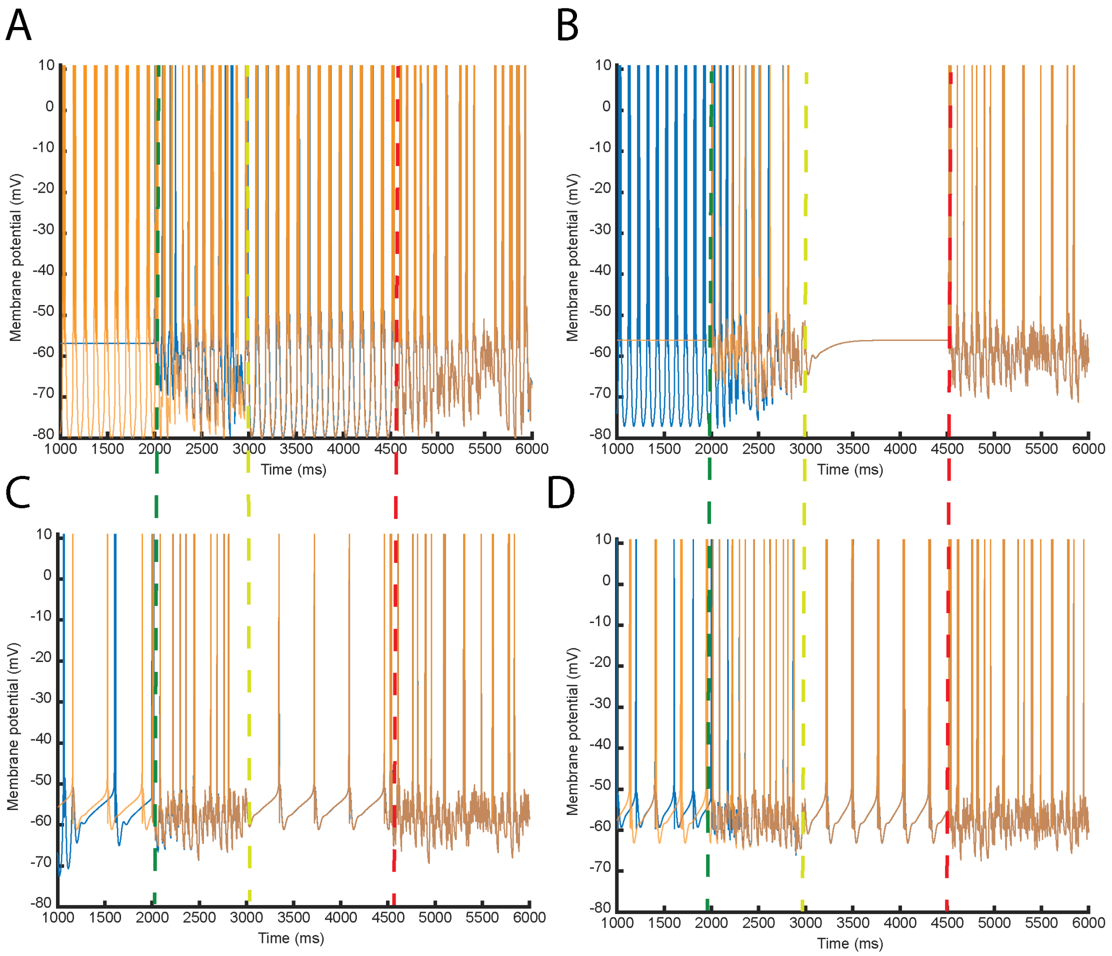


Supplemental Figure 6: **Bistability in regions II and III is overcome by noise.** A/B) Response of the neuron in region II starting with two different initial conditions (pink/blue). C/D) Response of the neuron in region III starting with two different initial conditions (pink/blue). All parameter values are standard except $g_{kl} = .03mS/cm^2$ and $I_{app} = 1.3, 1.6, 2.1, 2.4\mu A/cm^2$ respectively for A/B/C/D. In all cases (A/B/C/D) the simulation protocol is with no noise (before green line), after the introduction of noise (between green and yellow), after the cessation of noise (between yellow and red), and after the reintroduction of noise (after red). In all instances, regardless of the initial condition and firing mode seen prior to the green line, the firing mode post noise is identical between overlapping plots (blue/pink). Different areas of regions II/III can have different output firing type after the cessation of noise (bursting (A)/silent (B) in II, or single spike (C)/bursting (D) in III). The difference in what mode the neuron returns to has to do with the basin of attraction for the given bistable solutions. Initial investigations suggest that quiescence (II) and slow tonic spiking (III) are more likely to be observed close to the STW transition for higher values of $g_{kl}$. These results indicate that certain firing types may not be found under certain experimental conditions, since they can only be found in the absence of noise.

#### Supplement for section 2.6

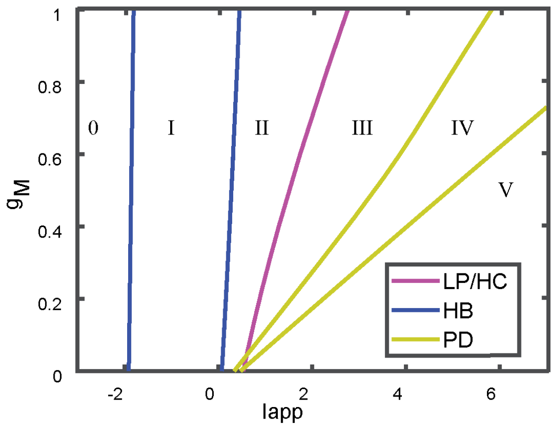


Supplemental Figure 7: **The full bifurcation diagram corresponding to Figure 5 in the main text.** The diagram is reproduced in the same manner as the diagram in Supplemental Figure 2. Here we are plotting a two parameter bifurcation diagram using $I_{app}$ and $g_M$. The curve color and region meanings are identical to those in Supplemental Figure 2. Although this bifurcation diagram possesses the same types of bifurcations as in Figure 3, there is now a meeting point between the homoclinic and period doubling that suggests the possible existence of bifurcations of higher than codimension two.

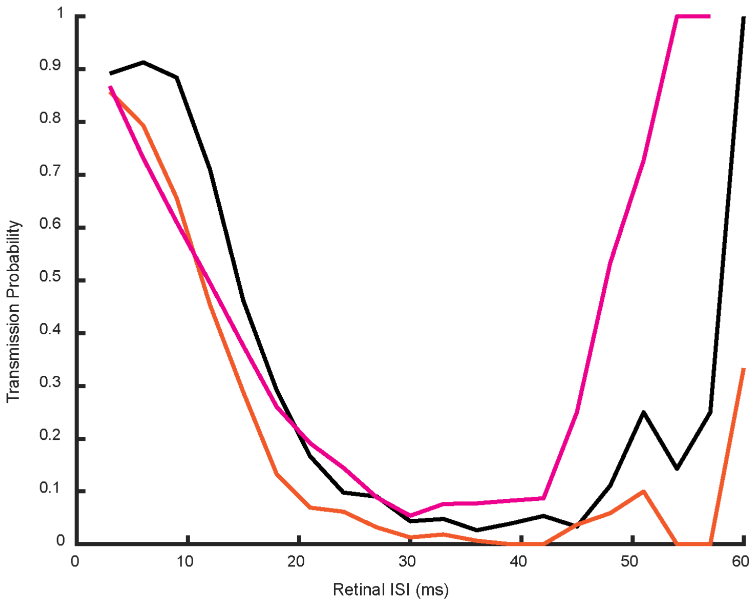


Supplemental Figure 8: **Detailed depiction of the relationships between key ionic currents and the successful transmission of retinal spikes when the TC cell is in state III.** Changes in the transmission probability of different retinal ISIs as a result of increasing $g_M$ (orange $g_M = .165mS/cm^2, g_{CaL} = .23mS/cm^2$) or decreasing $g_{CaL}$ (pink, $g_M = .12mS/cm^2, g_{CaL} = .15mS/cm^2$) when compared with the response of the control parameter set (black, $g_M = .12mS/cm^2, g_{CaL} = .23mS/cm^2$). Alterations in ionic conductances reveals that $g_M$ heavily influences the transmission of intermediate ISIs while $g_{CaL}$ controls the passages of short ISIs. These relationships between ionic conductances and retinal filtering are consistent across the filtering dynamic regions III and IV. In total, 10000 retinal spikes were simulated with ISIs grouped into 3ms bins.

#### Supplement for section 2.7

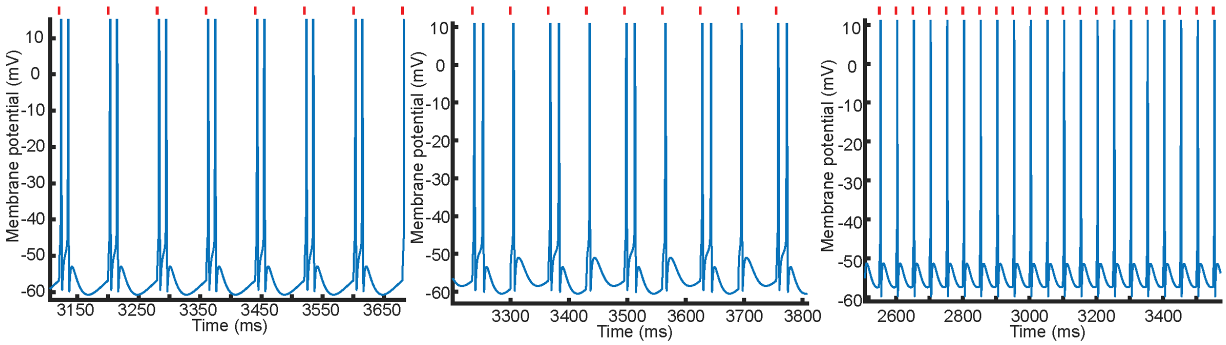


Supplemental Figure 9: **Three distinct firing types that occur during thalamic 1:1 entrainment to periodic excitatory inputs.** Successful transmission of periodic inputs depends on the level of entrainment. Note, entrainment is measured by comparing the ratio of input frequency to output frequency. The output within a single cycle of a periodic output could be a burst or a single spike. This means that there may be more output spikes than input spikes, but the neuron could still be considered to be entrained. As an example, during 1:1 entrainment to excitatory periodic inputs the TC cell exhibits three output types: entrained output as doublets (left), entrained output as mixed mode, meaning alternating doublets and singlets (middle), entrained output as single spikes (right). The mixed mode entrainment is interesting as it represents a firing model shift of the TC cell and can result in minor deviations from perfect 1:1 entrainment. The incoming input frequencies are 14Hz (left), 16Hz (middle), and 20Hz (right). Red tick marks denote timing of excitatory input spikes.

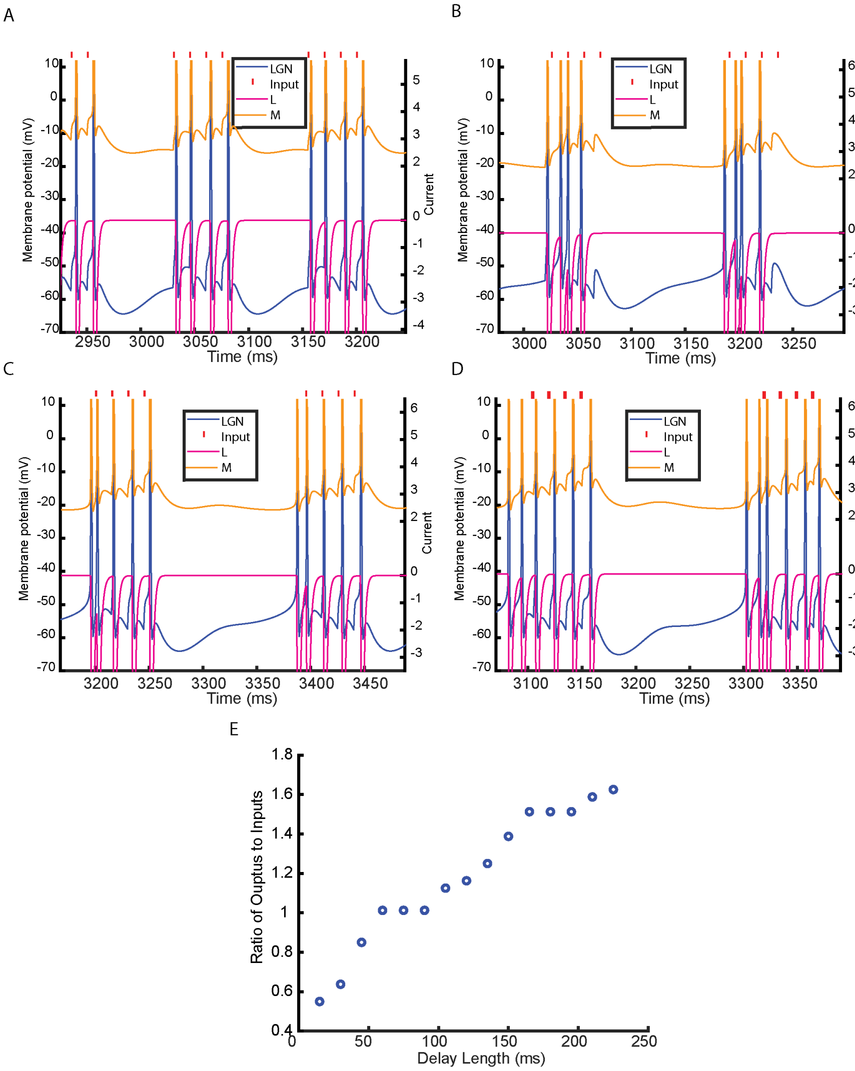


Supplemental Figure 10: **Different theta timescales interact with the underlying alpha dynamics (produced by M and L current time constants) to determine passage of gamma inputs.** The response of the TC cell to theta-gamma nested excitatory inputs depends on the incoming theta frequency. A-D) Response of the neuron to a nested theta-gamma excitatory signal where the gamma is at 66Hz for all plots and the theta is at 8, 6.5 ,5.1 and 4.5Hz for A-D respectively. The gamma input is faithfully transmitted in A, C and D but there are additional spikes coming from the underlying alpha dynamics appearing in the TC output. The faithful transmission of gamma input is disrupted in B where the underlying alpha spikes are appearing in the middle of the gamma timed inputs. E) Summary plot showing the ratio of total input spikes to output spikes as a function of different theta timescales with a fixed incoming gamma frequency (66Hz) and a fixed number of incoming spikes (4). Notably, the ratio of input to output spikes increases as the delay length between gamma bursts is increased due to the interaction with the underlying alpha rhythm.

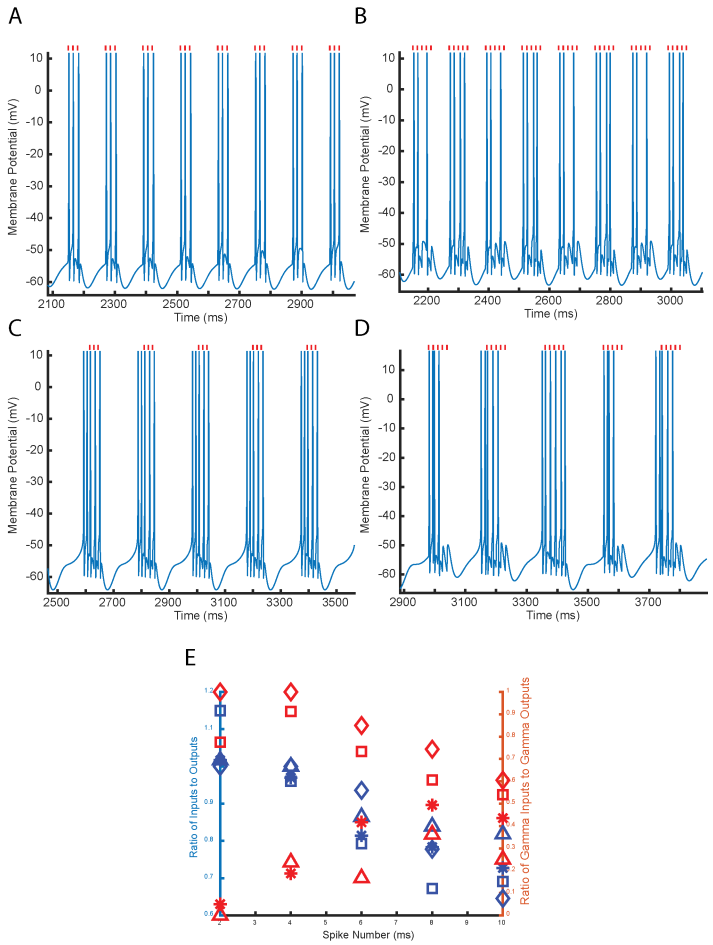


Supplemental Figure 11: **Successful response of the TC cell to incoming nested excitatory gamma rhythms also depends on the number and frequency of the incoming gamma spikes.** A) and B) Sample membrane potential trace of the neuron in state IV responding to 3 (A) or 5 (B) incoming gamma (66Hz or 15ms ISI) spikes per alpha (8Hz) cycle. The thalamus successfully passes all gamma spikes with 3 input spikes but fails to generate 100 percent passage of spiking for 5 spiking inputs. Red tick marks denote the timing of the input spikes. C) and D) Sample membrane potential trace of the neuron in state IV responding to 3 (C) or 5 (D) incoming gamma (66Hz or 15ms ISI) spikes per theta (5Hz) cycle. The thalamus successfully passes all gamma spikes with 3 input spikes but fails to generate 100 percent passage of spiking for 5 spiking inputs. Red tick marks denote the timing of the input spikes. Red tick marks denote the timing of the input spikes. E) Summary plot describing the relationship between the gamma spikes generated by the thalamus (in state IV) in response to different spike numbers and frequencies. Note the delay length (90ms) here is fixed for all simulations. Blue symbols correspond to the left axis which showcases the ratio of input spikes to output spikes regardless of frequency. Red Symbols correspond to the right axis and denote the ratio of incoming gamma spikes to outgoing gamma spikes at a similar ($\pm$3ms ISI) gamma frequency. Different symbol shapes correspond to different speeds of gamma inputs as follows: Triangle 25ms ISI, Asterisk 20ms ISI, Square 15ms ISI, and Diamond 10ms ISI. This figure indicates that increasing the number of spikes per slow cycle, or decreasing the input gamma frequency, decreases the reliability of gamma transmission. In fact, for slower gamma inputs (50Hz or 40Hz), regardless of the number of input spikes per slow cycle, less than fifty percent of the output spikes occur at a matching gamma frequency. As a final note, for all input frequencies tested, fewer spikes per slow cycle helps maximize the total (irrespective of frequency) percent transmission of inputs. Together these results suggest that faster gamma frequencies with fewer incoming spikes per slow cycle, are likely to be relayed better.

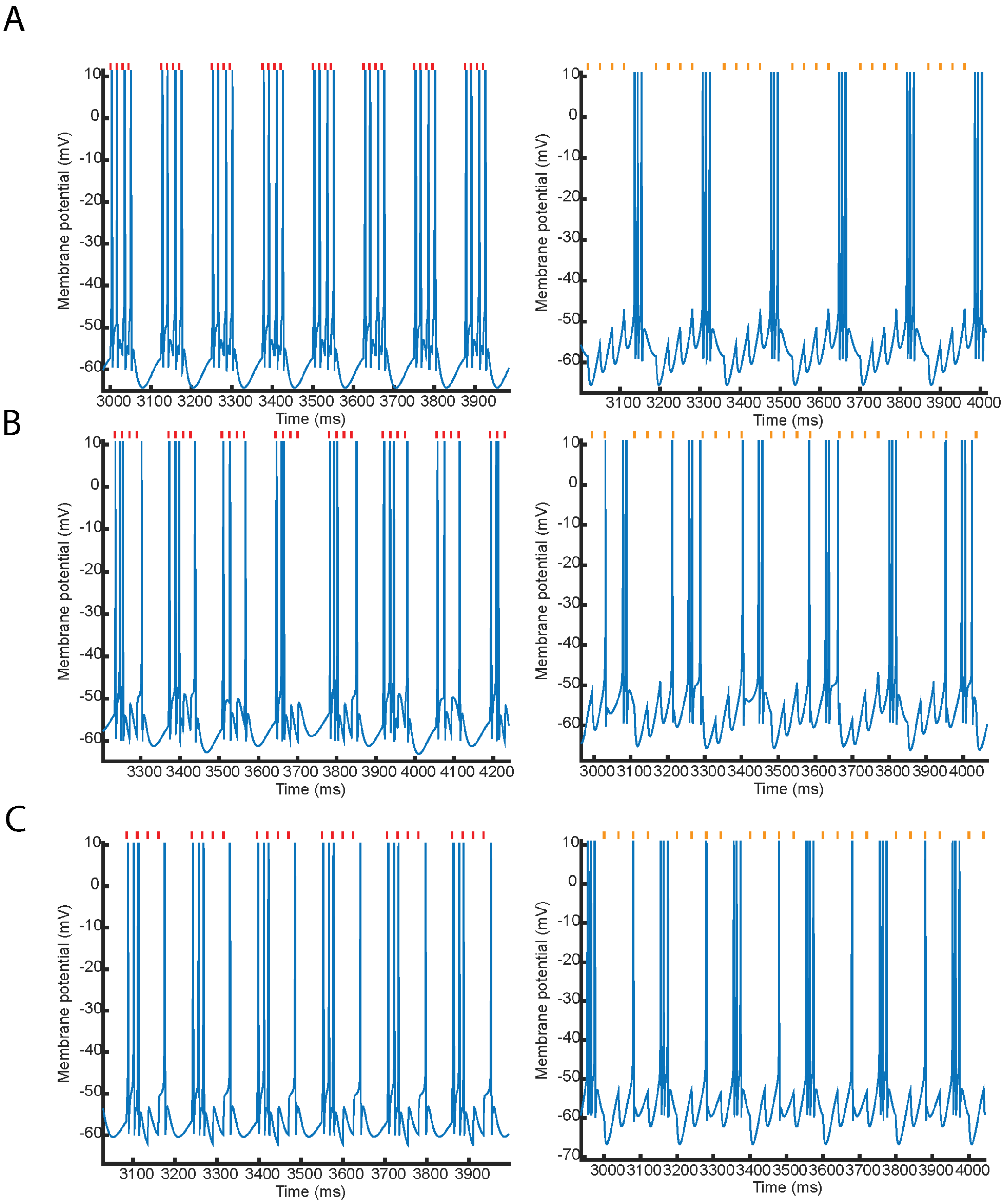


Supplemental Figure 12: **Output firing mode of the TC cell in response to nested theta/gamma excitatory and inhibitory inputs depends on the spiking gamma frequency.** The left column presents the TC response to excitatory nested gamma rhythms at 66Hz (A), 50Hz (B), and 40Hz (C). The right column presents the TC response to inhibitory nested beta/gamma rhythms with a fast frequency input at 35Hz (A), 30Hz (B) and 27Hz (C). For a fixed delay length (80ms), but with decreasing gamma frequency, the TC cell spiking output in response to both excitatory and inhibitory nested gammas is: 1:1 entrainment to the theta rhythm with a single gamma frequency (top), 1:1 entrainment to the theta rhythm with an inconsistent gamma spiking output (middle), 1:2 partial entrainment to the gamma input, alternating single spikes and bursts (bottom).

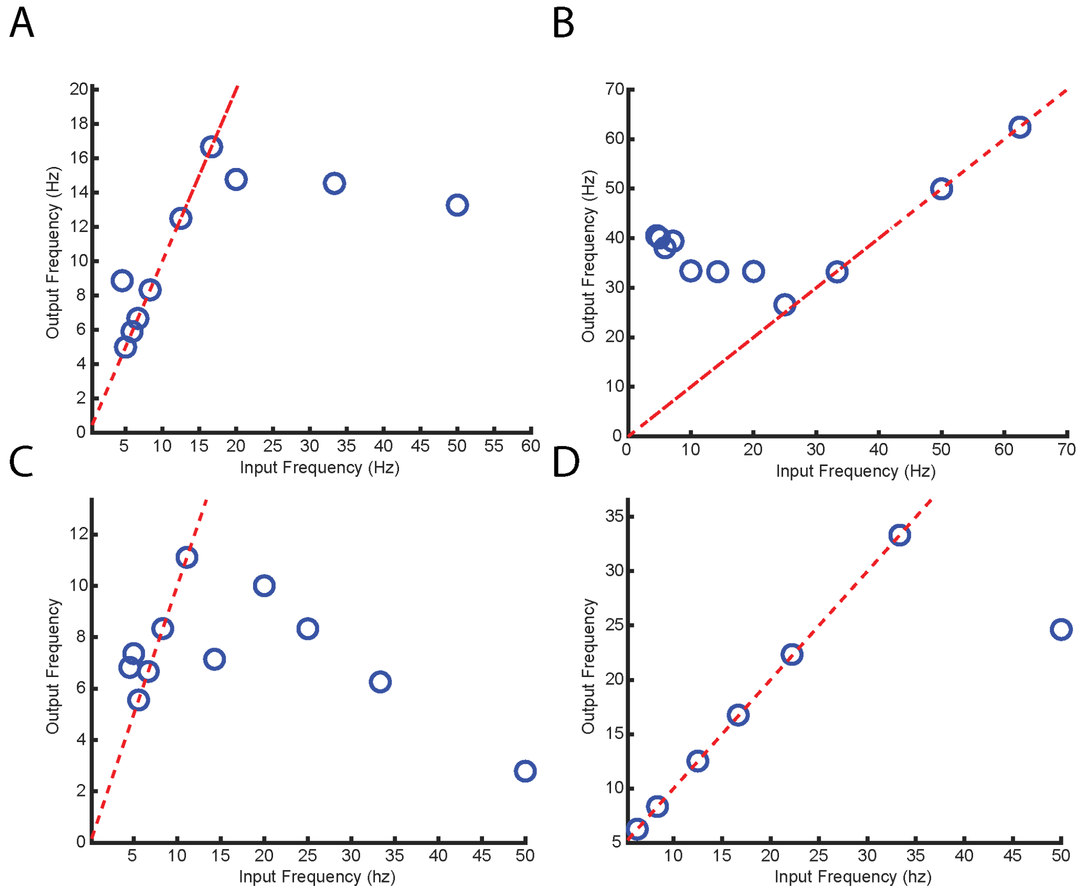


Supplemental Figure 13: **Transmission of periodic rhythms is generally less successful in states III and V compared to state IV.** A) Number of thalamic burst (or single spike) outputs per second in response to periodic rhythmic excitatory input for a TC cell in state III. B) Number of thalamic bursts (or single spike) outputs per second in response to periodic rhythmic excitatory input for a TC cell in state V. C) Number of thalamic burst (or single spike) outputs per second in response to periodic rhythmic inhibitory input for a TC cell in state III. D) Number of thalamic burst (or single spike) outputs per second in response to periodic rhythmic inhibitiory input for a TC cell in state V. Red line corresponds to 1:1 entrainment. Periodic stimulation of the neuron in state III entrains to a smaller frequency range than when in state IV (A and C). Periodic excitatory stimulation of the model TC cell in state V can only produce a gamma frequency output, and overrides, or ignores, all lower frequency inputs (B). Periodic inhibitory inputs to the model in state V entrains the neuron over a broad range of frequencies, however, the outputs consist of fast gamma spiking outputs interspersed with minor silent delays (not shown here) for all frequencies. This would imply that even for theta or alpha inputs the TC cell would also produce a very stronger gamma that wouldn't been seen during state IV entrainment.

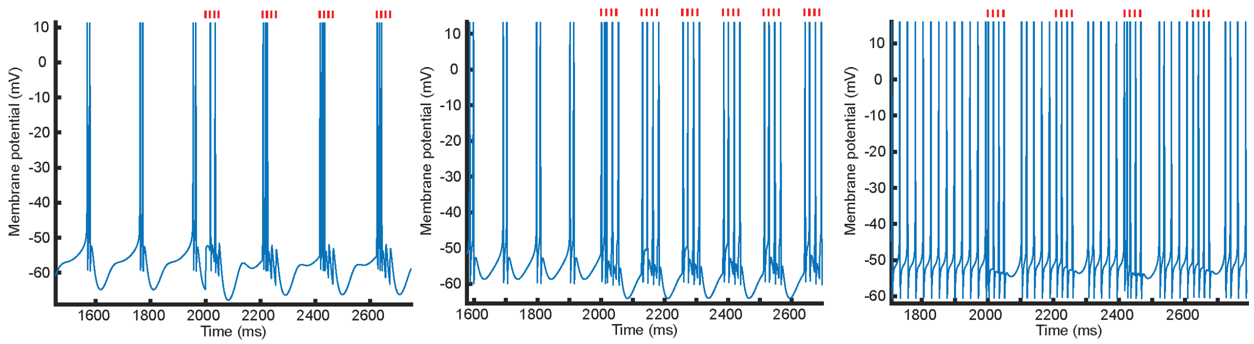


Supplemental Figure 14: **Thalamus passes incoming excitatory theta-gamma rhythms best when the underlying dynamics are in the alpha bursting mode.** When underlying dynamics are in state III (a slower high-threshold bursting at theta) (left), the output is a theta-gamma but not matching the input gamma frequency. In state IV (alpha bursting) (middle) the thalamus faithfully transmits both the theta and the gamma rhythms. When the thalamus is placed into dynamic state V, the thalamus generates predominately gamma, and not just in response to incoming inputs (right). However, note that in state V, a pause occurs after each input, thereby still producing a theta timescale in the output. Red dots denote the timing of the input spikes.

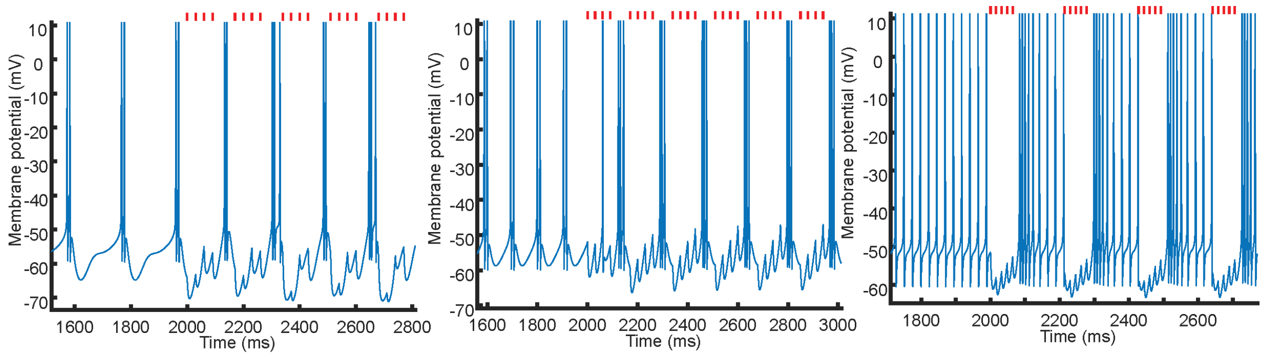


Supplemental Figure 15: **Thalamic response to incoming inhibitory theta-gamma (6Hz/33Hz) rhythms can differ depending on the initial underlying state.** In response to theta/gamma input, states III-V produce a consistent theta rhythm, but with different spiking responses. In state III, the spiking output alternates between doublet and triplets (Left). In state IV, the spiking output is consistently identical triplets (middle). In state V, the spiking output is a fast series of gamma spikes, followed by a slower set of spikes that occur at a frequency matching the underlying firing dynamics (right). Red dots denote the timing of the input spikes.

#### Supplement for section 2.9

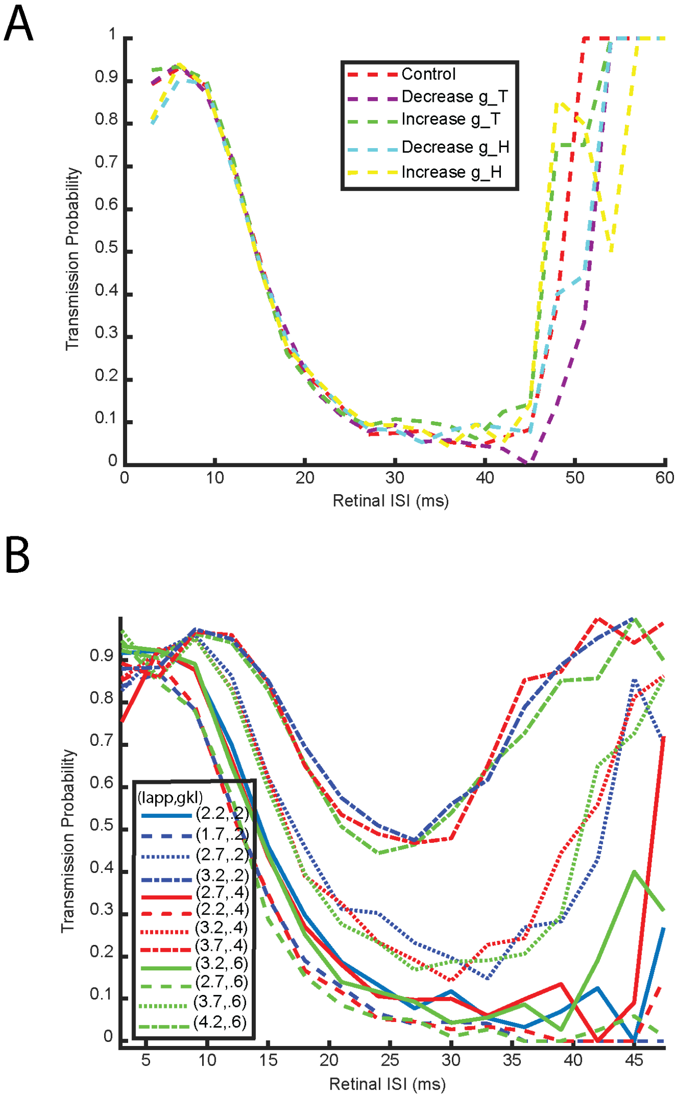


Supplemental Figure 16: **Retinal filtering is robust to changes in $g_H$ and $g_{CaT}$ and produces a similar filtering window across regions III and IV $I_{app}$ vs $g_{kl}$ excitation space.** A) Transmission probability as a function of Retinal ISI plotted for the control parameter set and 4 variations of this parameter set which are: 10% decrease in $g_{CaT}$ (purple), 10% increase in $g_{CaT}$ (green), 10% decrease in $g_H$ (blue), and 10% increase in $g_H$ (yellow). The four variations of the control parameter space produce a similar transmission probability curve. B) Transmission probability as a function of Retinal ISI plotted for 12 distinct parameter sets from within regions III and IV of $I_{app}$ vs $g_{kl}$ parameter space. Parameter sets were found by fixing a value of $g_{kl}$ and then taking four .5 increments of $I_{app}$. This was repeated for 3 separate values of $g_{kl}$ yielding the total of 12 parameter sets. We find that the effect of changing $I_{app}$ can be counteracted by changes in $g_{kl}$, and that throughout the entirety of the parameter space tested, the 20-40ms window of retinal ISIs are the most likely to be filtered out as described in the main text. The lowest (smallest transmission probability) two groupings of curves (by line type) are from region III and the higher two groupings of curves are from region IV in parameter space.

| Parameter Set 1 | Frequency | ISI | Minimum Voltage |
|---|---|---|---|
| **Control Parameters** | 9.25 | 10.52 | -59.75 |
| Inc T | 9.37 | 10.77 | -59.79 |
| Dec T | 9.13 | 10.24 | -59.72 |
| Inc L | 7.38 | 8.55 | -61.22 |
| Dec L | 8.97 | 17.99 | -60.27 |
| Inc M | 8.49 | 10.66 | -59.90 |
| Dec M | 10.03 | 10.67 | -59.85 |
| Inc H | 9.26 | 10.65 | -59.75 |
| Dec H | 9.25 | 10.40 | -59.75 |

| Parameter Set 3 | Frequency | ISI | Minimum Voltage |
|---|---|---|---|
| **Control Parameters** | 6.04 | 9.40 | -62.93 |
| Inc T | 6.04 | 10.55 | -62.74 |
| Dec T | 6.05 | 9.20 | -63.12 |
| Inc L by 5% | 10.33 | 8.09 | -69.39 |
| Dec L by 5% | 5.97 | 12.91 | -63.00 |
| Inc M | No HTB | No HTB | No HTB |
| Dec M | 6.93 | 9.49 | -61.94 |
| Inc H | 6.05 | 8.76 | -62.93 |
| Dec H | 6.04 | 9.10 | -62.93 |

| Parameter Set 2 | Frequency | ISI | Minimum Voltage |
|---|---|---|---|
| **Control Parameters** | 9.68 | 12.80 | -59.93 |
| Inc T | 9.76 | 13.09 | -59.95 |
| Dec T | 9.60 | 12.46 | -59.91 |
| Inc L | 9.74 | 9.62 | -59.57 |
| Dec L | No HTB | No HTB | No HTB |
| Inc M | 8.63 | 15.73 | -59.97 |
| Dec M | 10.88 | 13.63 | -59.99 |
| Inc H | 9.70 | 12.82 | -59.93 |
| Dec H | 9.68 | 12.75 | -59.92 |

| Parameter Set 4 | Frequency | ISI | Minimum Voltage |
|---|---|---|---|
| **Control Parameters** | 5.74 | 11.29 | -59.68 |
| Inc T | 5.78 | 11.63 | -59.71 |
| Dec T | 5.70 | 10.97 | -59.65 |
| Inc L by 5% | 5.77 | 9.00 | -59.44 |
| Dec L by 5% | No HTB | No HTB | No HTB |
| Inc M | 5.21 | 11.81 | -60.30 |
| Dec M | 6.40 | 11.06 | -59.69 |
| Inc H | 5.75 | 11.30 | -59.68 |
| Dec H | 5.74 | 11.28 | -59.68 |

Supplemental Table 1: **Changes in recorded values of the frequency, ISI and minimum voltage for high-threshold bursting neurons in response to variations of ionic conductances.** The four varied ionic conductances are $g_{CaT}, g_{CaL}, g_H$ and $g_M$. Conductances are increased and decreased by 10 percent (unless specified otherwise) and the resulting bursting metric is recorded. If the row is blank, the neuron stopped producing high-threshold bursts. We investigated four different points in states III and IV ($I_{app}$-$g_{kl}$) parameter space to generalize across the excitation space.

#### Supplement for section 5.2

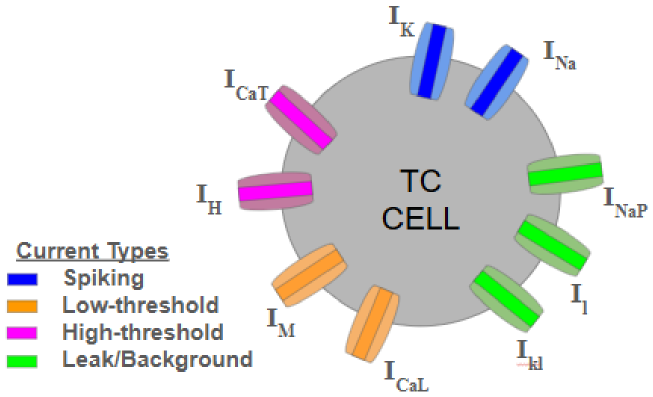


Supplemental Figure 17: **Schematic depicting the TC cell model structure and the contributions of the included ionic currents.** This model is a single cell, single compartment neuron model that has 9 ionic current types. These ionic currents are separated into 4 broad categories based on their primary functional role within the model: Spiking ($I_{Na}$ and $I_K$), Low-threshold bursting ($I_{CaT}$ and $I_H$), High-threshold bursting ($I_M$ and $I_{CaL}$) and Leak/background ($I_k l$, $I_l$ and $I_{NaP}$).

### Supplemental Methods

Differential equations, kinetic rate constants, ionic current equations and all associated parameter values for the computational TC cell are provided here. The equations representing the ionic current kinetics are taken from a variety of different modeling studies that can each be traced back to experimentally fit kinetic equations. An example modeling study using the given gated currents can be found at the following: $I_{Na}$ [4], $I_K$ [4], $I_{Nap}$ [5], $I_{CaT}$ [6] $I_{CaL}$ [4], $I_h$ [6], and $I_M$ [4]. These modeling studies base their kinetic equations from the following experimental works: $I_{Na}$ [7], $I_K$ [7], $I_{Nap}$ [8], $I_{CaT}$ [9] $I_{CaL}$ [10], $I_h$ [9], and $I_M$ [11]. All parameter values provided within these methods (including baseline excitation $I_{app} = 2.7\mu A/cm^2$) are set to produce high-threshold bursting at an alpha frequency. External excitation (or different ionic conductances) can then be varied to produce the other firing modes as desired/described in the main text.

The following differential equation describes a conductance based model where the change in membrane potential (V) of the TC cell is a combination of the ionic currents $\Sigma I_{ionic}$ and the applied current $I_{app}$ all scaled by 1 over the membrane capacitance $C_M$ (in units $\mu F/cm^2$):

$$\frac{dV}{dt} = \frac{1}{C_M}(I_{app} - \Sigma I_{ionic}). \tag{1}$$

The summed ionic currents are:

$$\Sigma I_{ionic} = I_{Na} + I_K + I_l + I_{kl} + I_{Nap} + I_{CaT} + I_{CaL} + I_h + I_M. \tag{2}$$

These currents are: spiking sodium $I_{Na}$, spiking potassium $I_K$, standard leak $I_l$, potassium leak $I_{kl}$, persistent sodium $I_{Nap}$, T-type calcium $I_{CaT}$, L-type calcium $I_{CaL}$, hyperpolarization activated H current $I_h$, and non-inactivating slowly decaying potassium M current $I_M$. The kinetics for these currents are adapted from previously published studies (see above) and are reproduced here for the readers. Each current type $j$ adheres to the Hodgkin-Huxley formulation:

$$I_j = g_j m_j^x h_j^y (V - E_j), \tag{3}$$

where $g_j$ is the maximal conductance for current type $j$, there are $x$ and $y$ numbers of activation $m$ and inactivation $h$ gates respectively, and the current has reversal potential $E_j$. The kinetics of each ionic gating variable have been fit based on voltage clamp experiments and follow one of two identical (a transformation from one to the other exists [12]) schemes. These schemes are provided here for an arbitrary gating variable $m$:

$$\frac{dm}{dt} = (m_{inf} - m)/\tau_m \tag{4}$$

$$\frac{dm}{dt} = \alpha_m(1 - m) - \beta_m m \tag{5}$$

where the kinetics are described either by the steady state open probability $m_{inf}$ and time constant functions $\tau_m$, or via opening ($\alpha_m$) and closing ($\beta_m$) kinetic rates. Using these gating equation formulations, the chosen modeled ionic currents are:

$$I_{Na} = g_{Na} m_{Na}^3 h_{Na} (V - E_{Na}) \tag{6}$$

$$I_K = g_K m_K^4 (V - E_K) \tag{7}$$

$$I_L = g_L (V - E_L) \tag{8}$$

$$I_{KL} = g_{KL} (V - E_{KL}) \tag{9}$$

$$I_H = g_H r (V - E_H) \tag{10}$$

$$I_{CAT} = g_{TLT} m_{TLT}^2 h_{TLT} (V - E_{CA}) \tag{11}$$

$$I_{CAL} = g_{cal} q_{cal}^2 r_{cal} (V - E_{CA}) \tag{12}$$

$$I_M = g_M p (V - E_K) \tag{13}$$

$$I_{NAP} = g_{NAP} m_{NAPinf} (V - E_{Na}). \tag{14}$$

(15)

Gating variables for the ionic currents are listed below:

| Current | Activation Open Prob | Activation time constant | Inactivation Open Prob | Inactivation time constant |
|---|---|---|---|---|
| $I_{Na}$ | $\frac{\alpha_m}{\alpha_m+\beta_m}$ | $\frac{1}{\alpha_m+\beta_m}$ | $\frac{\alpha_h}{\alpha_h+\beta_h}$ | $\frac{\alpha_h}{\alpha_h+\beta_h}$ |
| $I_K$ | $\frac{\alpha_K}{\alpha_K+\beta_k}$ | $\frac{1}{\alpha_K+\beta_k}$ | NA | NA |
| $I_L$ | NA | NA | NA | NA |
| $I_{KL}$ | Na | NA | NA | NA |
| $I_H$ | $\frac{1}{1+e^{((V+75)/5.5)}}$ | $\frac{1}{e^{(-.086V-14.59)}+e^{(.07V-1.87)}}$ | NA | NA |
| $I_{CAT}$ | $\frac{1}{1+e^{(\frac{-(V+62)}{6.2})}}$ | $.612 + \frac{1}{e^{(\frac{-(V+135)}{16.7})}+e^{(\frac{(V+19.8)}{18.2})}}$ | $\frac{1}{1+e^{((V+86)/4)}}$ | $30.8 + \frac{211.4+e^{((V+115.2)/5)}}{1+e^{((V+86)/3.2)}}$ |
| $I_{CAL}$ | $\frac{\alpha_q}{\alpha_q+\beta_q}$ | $\frac{1}{\alpha_q+\beta_q}$ | $\frac{\alpha_r}{\alpha_r+\beta_r}$ | $\frac{1}{\alpha_r+\beta_r}$ |
| $I_M$ | $\frac{1}{1+e^{(-(V+35)/10)}}$ | $\frac{1000}{3.3e^{((V+35)/20)}+e^{(-(V+35)/20)}}$ | NA | NA |
| $I_{NAP}$ | $\frac{1}{1+e^{(-(V+50)/5)}}$ | NA | NA | NA |

Supplemental Table 2: Open probability and time constant functional forms for both the activation and inactivation gating variables.

For instances where the open probability and the time constants are given in terms of opening ($\alpha$) and closing ($\beta$) rates, these rate constants are listed below. Here $Vt$ is a voltage shift (in mV) defined to as $Vt = V + 55$.

| Current | Activation $\alpha$ | Activation $\beta$ | Inactivation $\alpha$ | Inactivation $\beta$ |
|---|---|---|---|---|
| $I_{Na}$ | $\frac{0.32(13-Vt)}{e^{((13-Vt)/4)}-1}$ | $\frac{0.28(Vt-40)}{e^{((Vt-40)/5)}-1}$ | $0.128e^{((17-Vt)/18)}$ | $\frac{4}{1+e^{((40V-Vt)/5)}}$ |
| $I_K$ | $\frac{0.032(15-Vt)}{e^{((15-Vt)/5)}-1}$ | $0.5e^{((10-Vt)/40)}$ | NA | NA |
| $I_{CAL}$ | $\frac{.055(-27-V)}{e^{((-27-V)/3.8)}-1}$ | $.94e^{((-75-V)/17)}$ | $.000457e^{((-13-V)/50)}$ | $\frac{.0065}{e^{((-15-V)/28)}+1}$ |

Supplemental Table 3: Rate constants for gating variables that are defined in terms of opening and closing rates.

Finally, the maximal conductances and reversal potentials for the 9 membrane bound ionic currents and the 1 synaptic current (described in detail in the main text) are provided below:

| Current $I_j$ | Maximal Conductance $g_j$ (in $mS/cm^2$) | Reversal Potential $E_j$ (in mV) |
|---|---|---|
| $I_{Na}$ | 90 | $E_{na} = 50$ |
| $I_K$ | 4.4 | $E_k = -100$ |
| $I_L$ | .0195 | $E_l = -85$ |
| $I_{KL}$ | .012 | $E_k = -100$ |
| $I_H$ | .06 | $E_h = -43$ |
| $I_{CAT}$ | 4.04 | $E_{Ca} = RT/(zF)ln(Ca_e/Ca_i)$ |
| $I_{CAL}$ | .222 | $E_{Ca} = RT/(zF)ln(Ca_e/Ca_i)$ |
| $I_M$ | .12 | $E_k = -100$ |
| $I_{NAP}$ | .013 | $E_{na} = 50$ |
| $I_{syn}$ | .5 | Excitatory: $E_{syn} = 0$, Inhibitory:$E_{syn} = -80$ |

Supplemental Table 4: An initial set of maximal conductance values that produces high-threshold bursting and the fixed reversal potentials for all of incoming currents. Note, for all bifurcation diagrams plotting $I_{app}$ vs $g_{kl}$ (or $g_M$), the maximal conductances are divided by .06 and .3 respectively for numerical purposes.

For the two calcium currents, $I_{CaL}$ and $I_{CaT}$, the calcium flux and the corresponding reversal potential are explicitly time dependent and not just given as fixed parameter values. Notably $E_{Ca}$ varies according to the dynamic variable of internal Calcium $Ca_i$ whose changes are described by the ODE taken from [6] reproduced here:

$$\frac{dCa_i}{dt} = ((-I_{CaL} - I_{Ca_T})/(zFw)) - (Ca_i - Ca_r)/\tau_{Ca}. \quad (16)$$

All remaining parameters with their physiological meaning and value are given below:

| Parameter | Description | Value |
|---|---|---|
| T | Temperature | $310.35K$ |
| R | Gas Constant | 8.314 J/molK |
| z | Ca++ ion valence | 2 |
| F | Faraday Constant | 96485.3 C/mol |
| $Ca_e$ | external calcium concentration | 2mM |
| w | perimembrane shell thickness | $.5\mu m$ |
| $\tau_{Ca}$ | Ca++ removal rate | 10ms |
| $Ca_r$ | resting Ca++ concentration | .00005mM |
| $C_m$ | Capacitance | $1\mu F/cm^2$ |
| $\tau_d$ | Synaptic decay constant | Excitatory: .7ms, Inhibitory: 5ms |
| $\tau_r$ | Synaptic Rise constant | Excitatory: .125ms, Inhibitory: .25ms |
| $V_{max}$ | Synaptic cell max membrane potential | 10mV |

Supplemental Table 5: Additional fixed parameter values.
